# Interpretable Deep Learning for De Novo Design of Cell-Penetrating Abiotic Polymers

**DOI:** 10.1101/2020.04.10.036566

**Authors:** Carly K. Schissel, Somesh Mohapatra, Justin M. Wolfe, Colin M. Fadzen, Kamela Bellovoda, Chia-Ling Wu, Jenna A. Wood, Annika B. Malmberg, Andrei Loas, Rafael Gómez-Bombarelli, Bradley L. Pentelute

## Abstract

There are more amino acid permutations within a 40-residue sequence than atoms on Earth. This vast chemical search space hinders the use of human learning to design functional polymers. Here we couple supervised and unsupervised deep learning with high-throughput experimentation to drive the design of high-activity, novel sequences reaching 10 kDa that deliver antisense oligonucleotides to the nucleus of cells. The models, in which natural and unnatural residues are represented as topological fingerprints, decipher and visualize sequence-activity predictions. The new variants boost antisense activity by 50-fold, are effective in animals, are nontoxic, and can also deliver proteins into the cytosol. Machine learning can discover functional polymers that enhance cellular uptake of biotherapeutics, with significant implications toward developing therapies for currently untreatable diseases.

**One sentence summary:** Deep learning generates de novo large functional abiotic polymers that deliver antisense oligonucleotides to the nucleus.

## Main text

De novo design of functional polymers poses a significant sequence complexity challenge: for example, there are more permutations of 20 monomers over a 40 residue-long chain than there are atoms on Earth. This vast search space evades human intuition and prevents systematic investigation. Machine learning enables interpolation in high-dimensional search spaces by bridging the gaps between experimental training data points. Such strategies show promise to optimize functional molecules, but are typically hindered by datasets of inadequate size, consistency, and chemical feature representation.(*1*, *2*) For example, advancements in small molecule representation and generation have permitted de novo discovery of kinase inhibitors and identification of new antibiotics in a commercial database.(*3*, *4*) Prediction of peptide vaccines also benefitted from machine learning.(*5*) However, in all of these cases, the models use binary classifiers to predict candidates as a “hit” or “not a hit”. These binary classifiers are required because the available training data are not compatible with quantitative activity predictions because of experimental inconsistency. Initial attempts to address this limitation focused on the use of consistent data obtained with a standardized assay, but were still limited to a simple classifier because of insufficient size of the input dataset.(*6*)

We set forth to build a first-in-class model able to simultaneously design new functional polymers and quantitatively predict their activity. To attain this goal, a much larger standardized dataset and an advanced input representation was required.(*7*) We combined standardized, quantitative experimentation with versatile input representations and deep learning, focusing on tackling a major challenge in therapeutic development: delivery of functional macromolecules to the cytosol and nucleus. The cell membrane acts as a barrier to cytosolic delivery of extracellular proteins and foreign materials. While this feature is critical for the protection of the cell, it also prevents entry of many promising therapeutic macromolecules, stifling the development of biopharmaceuticals for diseases with intracellular targets.(*8*–*12*)

Antisense oligonucleotides have emerged as valuable tools for functional genomics, target validation, and more recently as therapeutics.(*13*) Structural modifications to the DNA backbone imparted high hybridization affinity to mRNA targets as well as greater stability towards nucleases. A prominent class of therapeutic antisense oligonucleotides with these features are charge-neutral phosphorodiamidate morpholino oligonucleotides (PMO). The first commercially-available PMO therapeutics were Eteplirsen (Exondys 51) and Golodirsen (Vyondys 53), which were approved by the FDA in 2016 and 2020, respectively, for the treatment of Duchenne muscular dystrophy.(*14*) A primary obstacle for clinical advancement and broad accessibility of these therapies is their poor cellular permeability.(*8*) High doses of PMO of up to 50 mg/kg are required for in vivo efficacy.(*15*) Approaches to deliver PMO using traditional vehicles such as liposomes and nanoparticles rarely advance to clinical trials, often suffering from endosomal entrapment or toxicity.(*16*)

One strategy to improve internalization of macromolecular therapeutic cargos is by covalently attaching them to cell-penetrating peptides (CPPs). CPPs typically range from 5 to 40 residues in length, have hydrophobic, hydrophilic, or amphipathic properties, and improve nuclear delivery of PMOs in vitro and in vivo.(*17*–*19*) Several CPPs are currently being investigated in clinical trials for delivery of therapeutic cargo, including PMO.(*1*, *2*, *20*, *21*) Cellular uptake of PMO typically occurs by energy-dependent uptake, meaning that endosomal escape presents an additional challenge. While some peptides can efficiently escape the endosome, designing a novel CPP sequence for this task is nearly impossible. In addition to the diversity of physicochemical properties of CPPs, variation in experimental design has resulted in inconsistent and sometimes contradictory datasets.(*22*) These inconsistencies preclude establishing sequence-activity relationships to guide the design of next-generation CPPs and can be remedied by testing PMO-CPP conjugates in a nuclear delivery-based assay that provides quantitative activity data and selects for sequences that can escape the endosome. In order to uncover CPP design principles for PMO delivery, it is necessary to have a standardized, biologically relevant dataset with which to train machine learning models.

Here we report a deep learning strategy with high predictive power fueled by robust input data containing unnatural residues and structures. A library containing 600 unique PMO-CPP conjugates was constructed using bioconjugation of peptide modules. Encoding residues as fingerprints allows us to expand beyond evolutionary big data and include abiotic monomers. Quantitative activity readout was achieved using an in vitro assay in which nuclear delivery of PMO results in enhanced green fluorescent protein (EGFP) fluorescence. Sequence matrices and corresponding activity data were used to train a predictor neural network. A “CPP thesaurus” dataset was used to train a generator neural network to produce novel “CPP-like” sequences. These novel sequences were optimized in the predictor-optimizer loop to increase predicted activity while minimizing length and arginine content to mitigate toxicity.(*23*) The output is hundreds of de novo designed sequences with a broad spectrum of predicted activity.

The model is also interpretable: we can visualize the decision-making process and identify structures that are consistent with empirical observations. From these predictions, we discovered best-in-class abiotic “Mach” (<u>Machine</u> Learning) peptides that improve PMO delivery by 50-fold and are safe for use in animals. Mach peptides are nontoxic and noninflammatory, and are able to deliver macromolecules other than PMO to the cytosol. Our model is application agnostic; with a dataset of sufficient size and consistency, our approach can be extended to the design of other functional materials. This strategy produces a broad spectrum of predicted activities and abiotic sequences that Nature is unable to design.

## Assembly of a Standardized Dataset

Recently, we demonstrated that linear combinations of known CPP sequences can synergistically improve delivery of PMO compared to each CPP alone.(*24*) We hypothesized that expanding this approach to a larger, more diverse library of linear combinations of CPPs would leverage chemical space to access a wide range of sequences and activities. We designed a synthetic method to assemble this library via bioconjugation of peptide “modules” into hundreds of novel PMO-CPPs. The resulting library diversity and spectrum of activities would be ideal to train machine learning algorithms (Fig. 1).

**Fig. 1.**
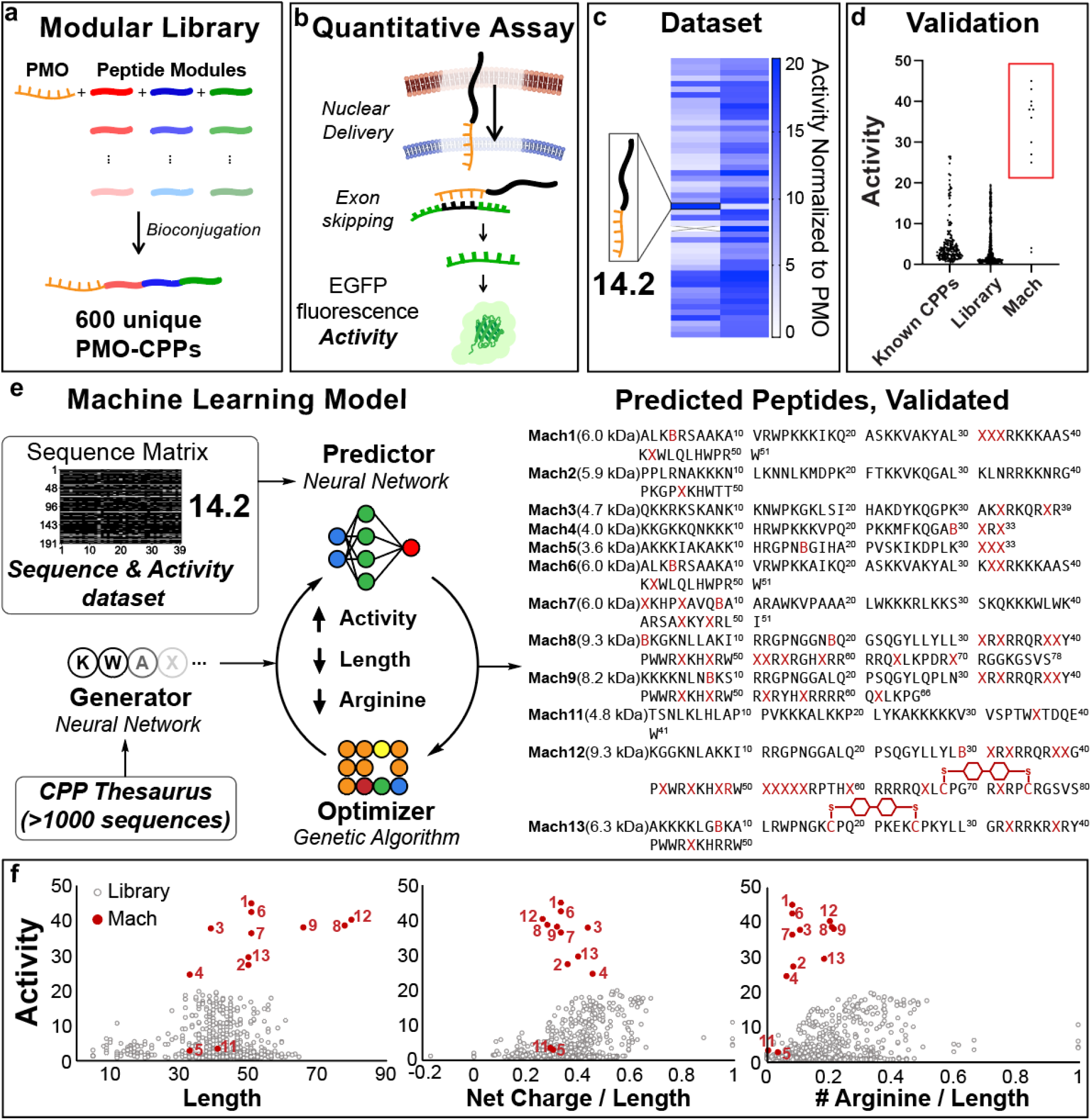
Machine learning model based on directed evolution predicts highly active abiotic peptides for macromolecule delivery. (a) A 600-membered library of PMO-CPP conjugates was synthesized using linear combinations of abiotic peptide modules. (b) A standardized in vitro activity assay tests for nuclear delivery using a quantitative fluorescence readout. (c) Members of the modular library exhibit a broad spectrum of activities. Each bit corresponds to a PMO-peptide in the library and its corresponding activity. (d) Peptides designed in this work have higher activity than the peptides in the modular library and known CPPs tested using the same assay. (e) Sequences are encoded into a matrix, labeled with experimental activity, and used to train a machine learning model. The model designs novel sequences in a loop based on directed evolution. X = aminohexanoic acid, B = β-alanine, C = cysteine macrocycles linked through decafluorobiphenyl. (f) Twelve predicted “Mach” peptides were synthesized and tested in the same activity assay (red) and compared to the library (grey) in relation to physicochemical properties.

Our synthesis strategy employs four modules: one for PMO and three for distinct pools of peptide sequences. Each pool contains peptides with diverse function and structure, including nuclear-targeting peptides and peptides containing noncanonical residues and cysteine-linked macrocycles (Table S1). To synthesize the constructs, we employed a convergent approach in which four modules are linked in a series of bioconjugation reactions that are chemoselective and irreversible, yielding products of sufficient crude purity for direct testing in vitro (SM 2.5).(*25*) The resulting library contained 600 members, composed of combinations of 57 total peptides.

The library exhibited a broad activity spectrum, quantified by a high-throughput functional readout assay.(*17*) In this assay, HeLa cells stably transfected with an EGFP gene interrupted by a mutated intron of β-globin (IVS2-654) produce a non-fluorescent EGFP protein. Successful delivery of PMO IVS2-654 to the nucleus results in corrective splicing and EGFP synthesis. The amount of PMO delivered to the nucleus is therefore correlated with EGFP fluorescence, quantified by flow cytometry. Activity is reported as mean fluorescence intensity (MFI) relative to PMO alone (Fig.1d, Fig. S1). The most active construct improved PMO delivery by nearly 20-fold, while the median activity was 3-fold. The resulting dataset was broad in terms of both peptide sequences and range of activity (Fig. S2).

## Developing the Machine Learning Model

Inspired by directed evolution, we leveraged fingerprint sequence representations to develop a machine learning-based generator-predictor-optimizer triad (Fig. 1e). In this framework, the generator produces novel cell-penetrating peptide sequences, the predictor quantitatively estimates the activity for a given sequence, and the optimizer evolves towards the most optimal sequence.

The standardized dataset of activity-labeled sequences from the modular library allowed for development and training of a quantitative regressor algorithm. This approach enabled us to overcome the limitations of other efforts in the literature which employed binary classifiers of active versus inactive sequences.(*3*–*5*, *26*–*28*) These previous predictors were mostly trained using physicochemical descriptors, with datasets obtained from non-standardized experiments and containing only canonical residues.(*6*, *29*–*31*) Inclusion of chemically diverse unnatural moieties is challenging because such physicochemical descriptors may not be readily available. The ability to encode for unnatural residues, however, would greatly expand the chemical search space, and may enhance macromolecule delivery.(*32*) Therefore, to predict activity of de novo-designed abiotic peptides, we used topological representations that capture atomic connectivity for residues and row matrix representations for sequences. Combined with quantitative experimental readouts, this novel polymer representation allows us to access the diverse pool of noncanonical residues and quantitatively predict activity.

Peptide sequences are represented as row matrices comprised of residue fingerprints. Individual residue fingerprints are bit-vectors based on the molecular graph of the residue, where atoms are treated as nodes and bonds as edges (Fig. 2a, SM Appendix III).(*33*) Each bit in the vector corresponds to a substructure, and is active/inactive depending on the presence/absence of the particular substructure. Representing residues as chemical structures, rather than discrete choices, allows for the encoding of both canonical and noncanonical residues and leverages chemical similarity between residues. The fingerprints are then compiled into a row matrix to encode the amide backbone of the peptide sequence (Fig. 2b).

**Fig. 2.**
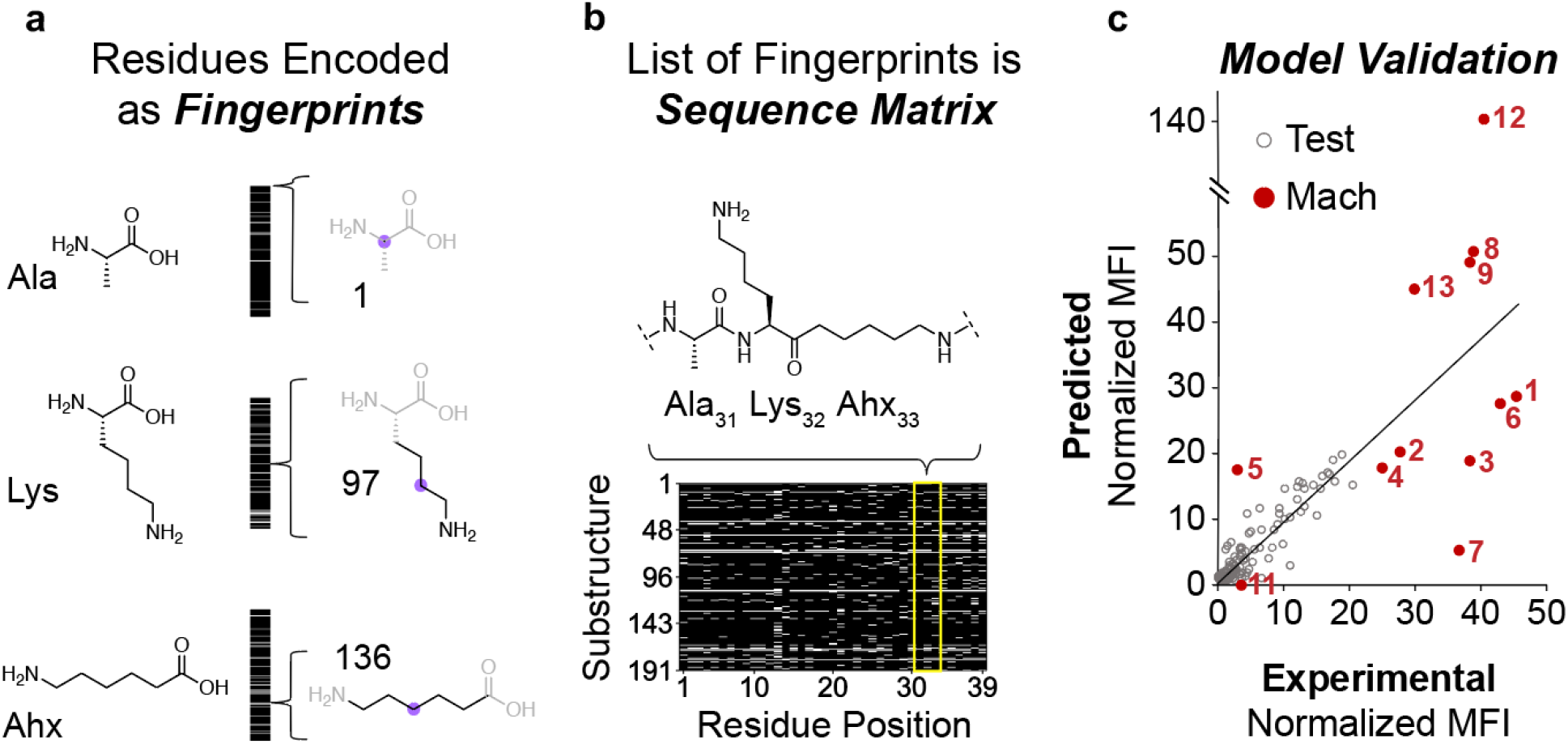
Machine learning-based generator-predictor-optimizer loop predicts highly active, novel abiotic CPPs. (a) Each amino acid residue is represented as a unique fingerprint, constructed as a bit-vector encoding for the presence or absence of 191 possible substructures in the residue. (b) Sequences are represented as residue fingerprints stacked in a row matrix. (c) Performance of machine learning model, comparing the predicted and experimental activity values for the holdout test set and novel Mach sequences.

The predictor neural network quantitatively estimated normalized MFI for a given sequence. Pairs of sequence representations and corresponding experimental activities were used to train a convolutional neural network (CNN). The training dataset consisted of PMO-CPPs from the modular library as well as other conjugates previously tested in the same assay.(*6*) A randomly-selected 20% of the dataset was saved for validation of the predictive accuracy of the algorithm. The root mean squared error on the validation set was 0.4 of the standard deviation of the training data. The prediction accuracy was found to be 89% as long as the predicted activity fell within the range of training values (normalized activity of 0.32-19.5) (Fig. 2c). Finally, we benchmarked our model against six other representations and model architectures and found that our strategy has the highest prediction power (Table S2).

We developed a generator based on a recurrent neural network (RNN) that captured the ontology of CPPs and generated CPP-like sequences. We trained the generator using a nested long short-term memory (LSTM) neural network architecture, which is better able to capture long-range correlations in sequence data.(*34*) We trained the algorithm using a “CPP thesaurus,” a collection of sequences from both our modular library and the literature.(*35*) Because the model is learning sequence grammar and has no role in activity predictions, no quantitative labels are necessary and we can use a large dataset of available sequences.

The optimizer completed the loop based on directed evolution. Sequences from the generator were randomly mutated and evaluated against an objective function, which maximized activity as predicted by the CNN model, and minimized length and arginine content while retaining water solubility estimated with net charge of the sequence. After 1000 iterations over each sequence, the model delivered hundreds of unique sequences with a wide range of predicted activity values. Along with highly active sequences, we predicted inactive sequences as negative control. By directing the evolution of the optimizer in the opposite direction, i.e., minimizing MFI, but keeping other constraints the same, we were able to generate an inactive sequence (Mach11) that appeared similar in amino acid composition to the active predictions. After synthesis, the Mach11 conjugate demonstrated minimal experimental activity, demonstrating the robustness of the model in accurately predicting the activity of an unknown sequence.

We confirmed that the predicted sequences were unique compared to known peptides and proteins (Fig. S3). We found that the abiotic peptides were unique compared to training set using text distance algorithms, which calculates how many steps are required to transform one sequence into another. In addition, by replacing unnatural residues to the closest natural residue, we found no similarity between Mach peptides and naturally occurring peptides and proteins as determined by a BLASTp homology search against the UniProt database.(*36*)

We can interpret the predictor CNN by visualizing the residue substructures that are important in its decision-making process. This type of visualization was a longstanding challenge that was recently addressed for image classification.(*37*) We developed an analogous tool to correlate the input sequence representation with predicted activity. This process generated bit-wise positive and negative activation values for each chemical substructure in the sequence. Bits with higher activation indicated the features that most strongly influence the final activity prediction.

As an example, for the predicted Mach3 sequence the two C-terminal aminohexanoic acid (Ahx) residues were the most positively activated (Fig. 3a), followed by arginine (Arg). The alkyl backbone in Ahx was the most activated substructure (Fig. 3b). A similar trend was observed for active sequences and substructures in the training dataset (Fig. S4, S5).

**Fig. 3.**
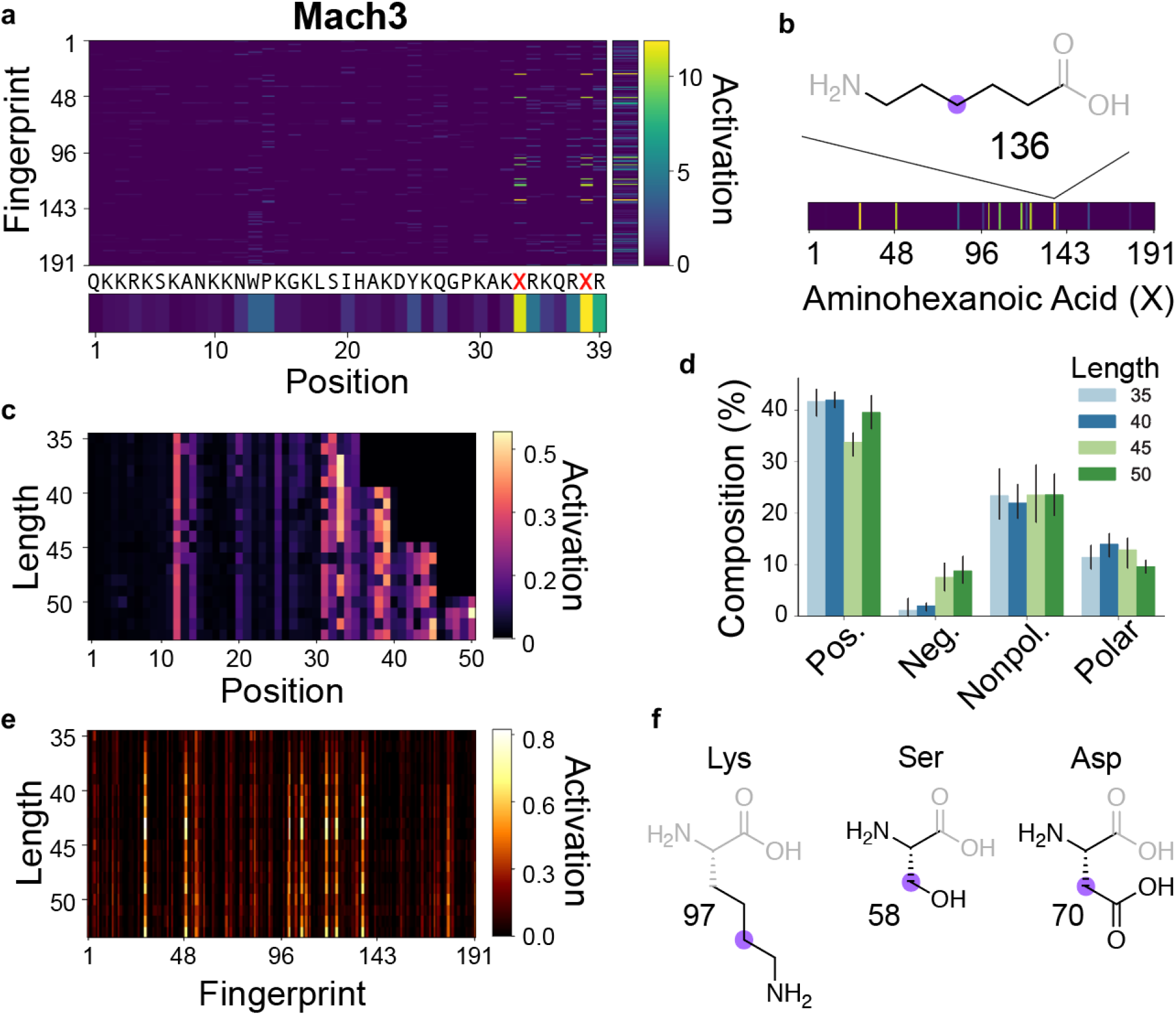
Interpretation of predictor CNN unveils activated substructures. (a) CNN positive activation gradient map was calculated for input sequence representation of Mach3. The averaged activation values over fingerprint indices and residue positions are shown. Fingerprint index corresponds to a corresponding substructure. (b) The alkyl backbone of aminohexanoic acid was the most positively activated substructure in Mach3 (c) Gradient maps of predicted sequences with lengths 35, 40, 45 and 50 showed that C-terminal residues are the most activated. (d) Composition of each type of residue remains unchanged regardless of sequence length of predicted peptides. (e) Several residues and substructures are consistently activated across all sequence lengths, including the amine side chain of Lys, polar side chain of Ser, and the carboxylic acid side chain of Asp.

We used this visualization approach to better understand how the trained model designed sequences. We chose five random sequences of different lengths, seeded them in the predictor-optimizer loop to maximize activity contingent upon other design constraints, and visualized the activations for the best predictions. Again, a higher activation can be seen for C-terminal residues (Fig. 3c). We also observed that the general composition of charged and hydrophobic residues remained unchanged across different sequence lengths (Fig. 3d). Particular residue fingerprints were activated irrespective of the sequence length, such as the side chains of Lys, Ser, and Asp (Fig. 3e-f). Consistent with earlier observations, a strong preference for polar and charged side chains as well as for Ahx was evident.

## Mach peptides enhance PMO delivery

We synthesized and characterized twelve candidates from hundreds of peptides predicted by the model, selecting diverse sequences and predicted activities. Using automated fast-flow solid-phase peptide synthesis, we reliably synthesized each predicted sequence on a 100 μmol scale.(*38*) Subsequently, each peptide was capped with 5-azidopentanoic acid for click chemistry (Fig. S6). If the predicted sequence contained a cysteine-linked macrocycle, we utilized SNAr chemistry to link the two cysteine residues with decafluorobiphenyl as previously reported.(*17*) Reverse-phase HPLC purification afforded our “Mach” peptides with a functional handle for conjugation to alkyne-labeled biomolecules. Finally, conjugation of azido-Mach peptides to PMO IVS2-654 was achieved in the same manner as in the library. The final PMO-Mach constructs are described in Table S3.

Experimental activity fell within the error range of predicted activity as long as the sequences were in the dynamic range of the training dataset (Fig. 2c). The PMO-Mach constructs were first tested for PMO delivery in the HeLa 654 assay at 5 μM and analyzed by flow cytometry as was done with the library (Fig. S7). We compared the resulting activity to the training dataset and found that nearly all sequences predicted to have activity greater than 20-fold did indeed surpass the highest performing library peptide, with the exception of Mach5.

Physicochemical properties of validated predictions show little correlation with PMO activity. Mach1 through Mach11 are linear peptides. Mach12 and 13 contain two cysteines linked by decafluorobiphenyl to form an internal macrocycle (Fig. S6). Circular dichroism suggested the peptides have limited secondary structure (Fig. S8). We compared the activities of Mach peptides to library peptides in relation to various physicochemical properties (Fig. 1f). While library peptides clearly show an increase in activity with an increase in length, arginine content relative to length and net charge relative to length, there is no obvious correlation between activity of Mach peptides and these same properties. These observations suggest that the model is taking advantage of sequence-activity relationships that go beyond sequence length and charge.

## PMO-Mach peptides have large therapeutic indices

PMO-Mach constructs have greater potency than previously characterized PMO-CPPs, while remaining nontoxic. This type of macromolecular delivery is a historic challenge, often suffering from either membrane disruption or endosomal entrapment. We first verified that PMO-Mach constructs enter cells via energy-dependent uptake using a panel of chemical endocytosis inhibitors and the HeLa 654 assay (Fig. S9). We then performed dose-response experiments (Fig. 4a-c, Fig. S10). PMO-Mach2, 3, 4, and 7 each had an EC_50_ value near 1 μM and were nontoxic at the concentrations tested, as determined by viability staining with propidium iodide (PI) and lactate dehydrogenase (LDH) release assay (Fig. S11). We compared these results to a previously well-performing CPP for PMO delivery, Bpep-Bpep.(*24*) This peptide has similar activity, but is composed of mostly Arg residues and exhibits cytotoxicity at higher doses (Fig. S11).

**Fig. 4.**
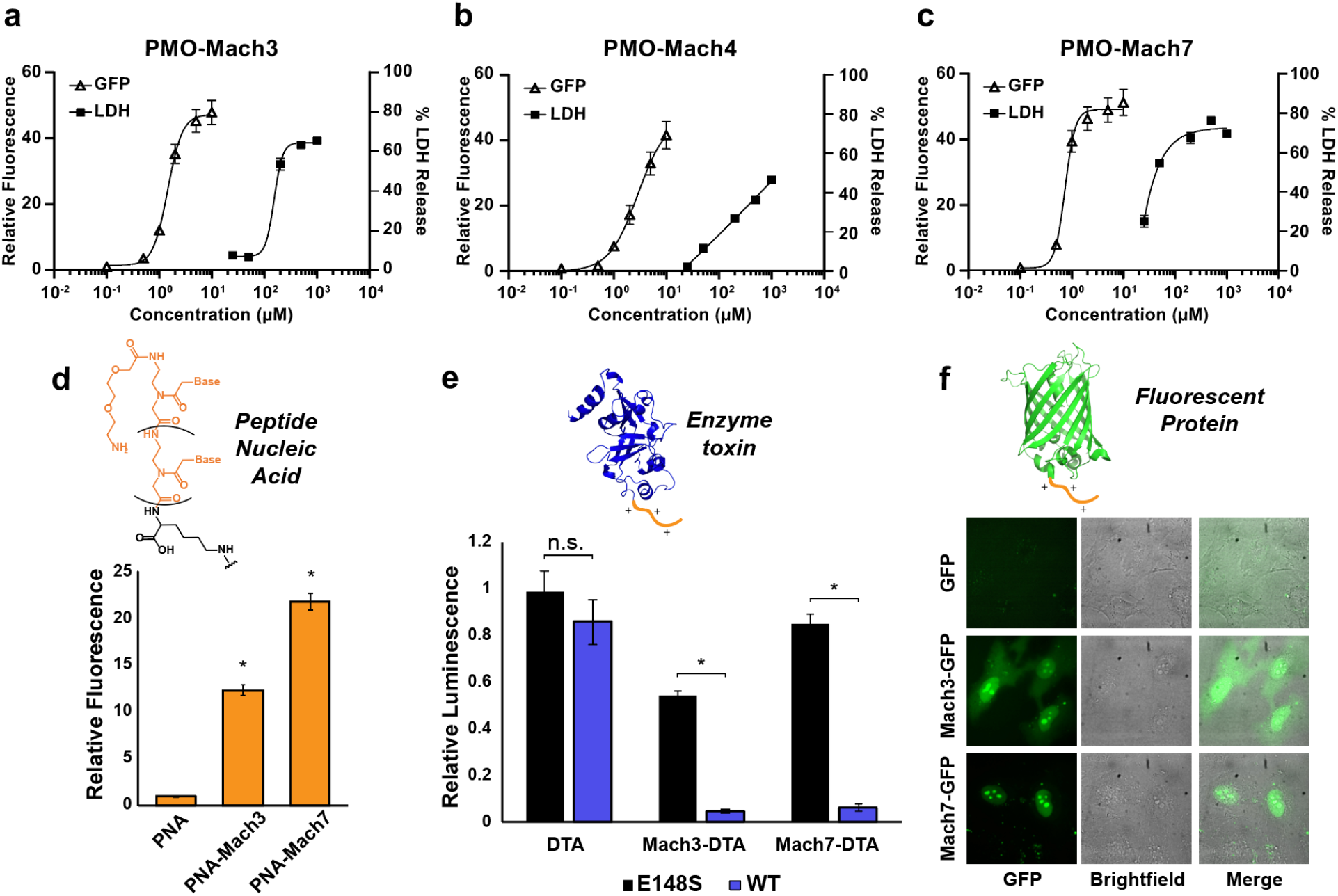
Mach peptides are highly active and deliver other biomacromolecules into the cytosol of mammalian cells. (a-c) Several PMO-Mach constructs have a large therapeutic index in vitro. Activity was determined using the EGFP assay: HeLa 654 cells were incubated with PMO-Mach constructs for 22 h before analysis by flow cytometry. Results are shown as fold increase relative to PMO alone. Toxicity was determined using renal epithelial cells (RPTEC TH1) treated in the same fashion and analyzed using LDH release assay (mean ± SD, for EGFP assay n = 5 for PMO-Mach3 and PMO-Mach7, and n = 6 for PMO-Mach4. For LDH assay, n = 4). (d) Mach peptides conjugated to PNA 654 significantly enhance PNA activity compared to PNA alone. HeLa 654 cells were incubated with PNA-Mach construct at 5 μM for 22 h before analysis by flow cytometry. (e) Mach3 and Mach7 attached to the N-terminus of Diphtheria toxin A (DTA) enhance the delivery of the toxin into the cytosol. HeLa cells were treated at 800 nM for 48 h before cell proliferation was measured with the CellTiter-Glo assay. (f) EGFP, Mach3-EGFP, or Mach7-EGFP were incubated with HeLa cells at 10 μM for 3 h. The treatment media was replaced with fresh media before imaging by confocal microscopy. Each bar in (d) and (e) represents group mean ± SD, n = 3, (*) p < 0.01, student’s two-tailed t-test.

PMO-Mach2, 3, 4, and 7 have a large therapeutic index as determined by dose-response cytotoxicity experiments in renal cells. We tested PMO-Mach constructs in human renal proximal tubule epithelial cells (RPTEC TH1, ECH001, Kerafast). Treatment supernatant was analyzed for LDH release, and no toxicity was observed even at the highest concentration needed for maximum PMO activity in HeLa 654 cells (Fig. 4a-c, Fig. S12). In contrast, Bpep-Bpep showed high toxicity above 10 μM. PMO-Mach conjugates have high activity, low arginine content, and a wide therapeutic window, highlighting their suitability for cytosolic and nuclear delivery.

## Mach peptides deliver other biomacromolecules

Mach peptides are versatile in that they can deliver other large biomolecules to the cytosol. Peptide nucleic acid (PNA) is a class of synthetic antisense oligonucleotides that has the same mechanism of action as PMO but has a highly flexible backbone structure.(*39*) We tested for delivery of a PNA variant of PMO 654 that is compatible with the EGFP assay. Each of the four Mach peptides tested was able to significantly enhance PNA delivery (Fig. 4d, Fig. S13).

In addition to antisense oligonucleotides, Mach peptides can also deliver charged proteins, such as Diphtheria toxin A (DTA). DTA is a 21 kDa anionic protein segment containing the catalytic domain of the toxin but lacking the portions that endow cell entry.(*40*) Delivery of this enzyme can be monitored using a cell proliferation assay as it inhibits protein synthesis in the cytosol. We found that Mach-DTA constructs were delivered into the cell cytosol significantly more efficiently than protein alone, and that covalent linkage was required for delivery (Fig. 4e, S14). Furthermore, we confirmed that toxicity is due to the cytosolic delivery of active DTA by comparing the wild-type constructs to those containing DTA(E148S), a mutant with 300-fold lower activity that the wild-type.(*41*) As expected, the mutant DTA conjugates lead to significantly reduced toxicity. Finally, investigation into the uptake mechanism of Mach-DTA suggested energy-dependent uptake similar to the PMO-Mach constructs (Fig. S15).

Conjugation to Mach peptides also improves the delivery of EGFP, a fluorescent protein commonly used as a reporter. After incubation of HeLa cells with 10 μM Mach-GFP, confocal micrographs displayed diffuse green fluorescence in the cytosol and intense fluorescence in the nucleus after incubation with Mach-EGFP (Fig. 4f). This observation is in contrast with the EGFP alone condition, in which no diffuse fluorescence was observed in either location, indicating reduced uptake.

## PMO-Mach peptides are nonimmunogenic and restore protein synthesis in vivo

After verifying Mach peptides propensity for in vitro macromolecule delivery, we looked towards in vivo antisense applications. We confirmed that PMO-Mach constructs are not immunogenic in vitro and restore EGFP protein synthesis in transgenic mice. In vitro tests with human macrophages suggested that the constructs are not inflammatory (Fig. S16). Existing predictive models also suggest that Mach peptides would not be T cell epitopes (Fig. S17).

Lastly, we demonstrated that PMO-Mach constructs correct protein synthesis in animals. Transgenic mice containing the same EGFP IVS2-654 gene as the one used in cell assays were given a single intravenous injection of varying doses of PMO-Mach3 or PMO-Mach4 and evaluated after seven days. Both constructs exhibited a dose-dependent increase in EGFP expression in quadriceps, diaphragm, and heart (Fig. 5a-c). PMO delivery to the heart is a critical but challenging objective. Here we observe similar levels of protein synthesis in both skeletal and cardiac tissue. In addition, there were no significant changes in the level of renal function biomarkers seven days post-treatment (Fig. 5d-f). The lack of renal biomarkers elevation suggests limited nephrotoxicity of the PMO-Mach conjugates and their viability as delivery materials for PMO to muscle tissue.

**Fig. 5.**
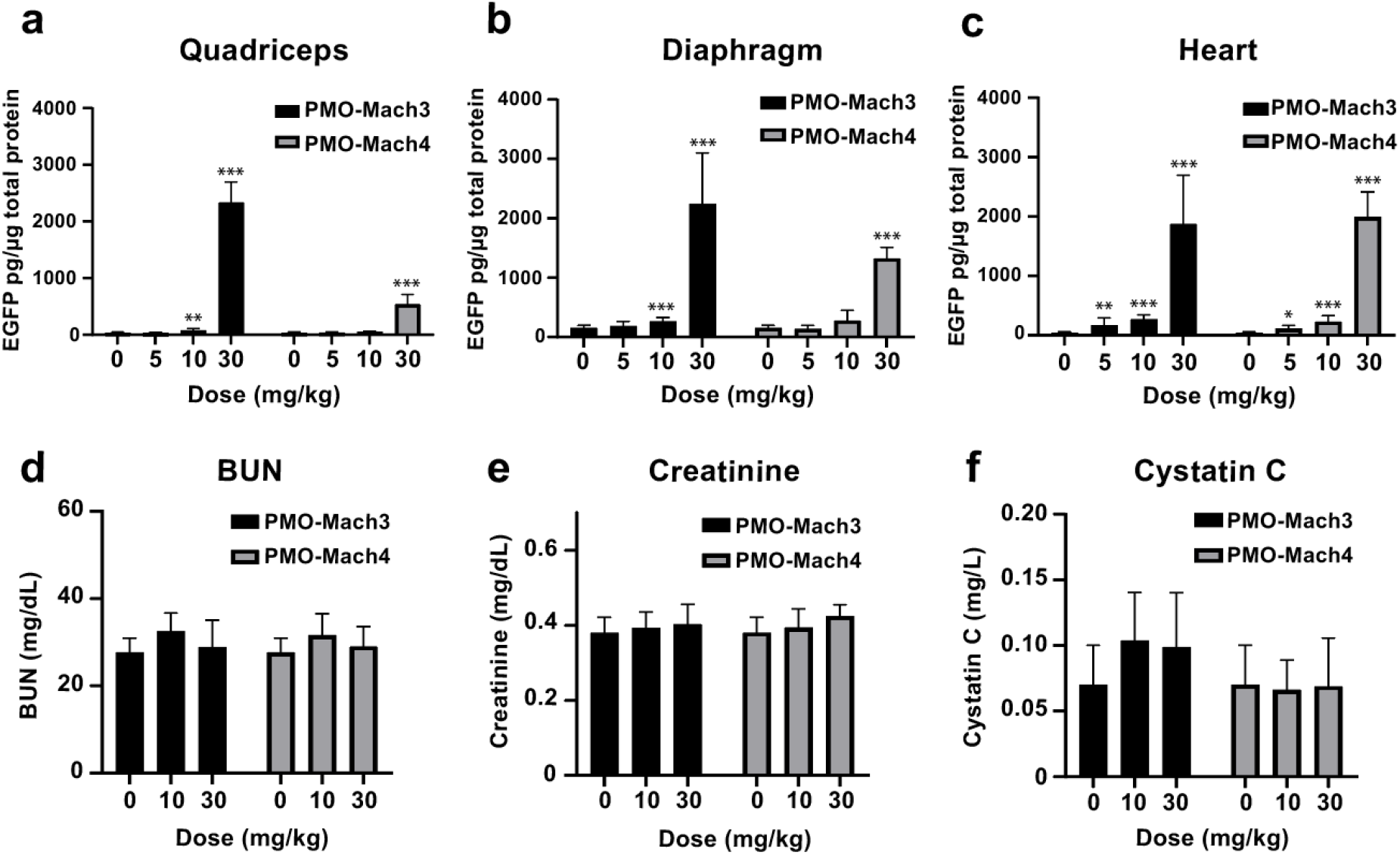
Machine learning-designed peptides are high-performing, safe delivery materials in mice containing the same transgene as HeLa 654 cells. Dose-response EGFP protein level in (a) quadriceps (b) diaphragm and (c) heart. Levels of (f) blood urea nitrogen (BUN), (g) creatinine, and (h) cystatin C remained unchanged. Saline (n = 6), Mach3 and Mach4 at 5 mg/kg (n = 4), all other n = 8. Each bar represents group mean ± SD. Mann-Whitney U test. *p<0.002 **p<0.0002 ***p<0.0001

## Discussion

We demonstrate a method to access the vast chemical search space of functional polymers using deep learning and standardized experimentation. Our model was applied to the design of abiotic peptides that can deliver an antisense PMO to the nucleus with greater efficiency than any previously known variant. The core strengths of our model lie in: (1) the topological representation of residues and sequences, (2) standardized quantitative activity data, and (3) a visualization tool to interpret the decision-making process of the model.

Synthesis and testing of the modular PMO-CPP library produced a broad spectrum of sequence and activity data with which we trained the model. By representing peptide sequences as topological fingerprints rather than descriptors like molecular weight, charge, and hydrophobicity, the model has access to more information to learn from during training and can be transferred to new chemistries without requiring physicochemical descriptors. The standardized data allowed us to use a quantitative regressor to design novel sequences with a broad spectrum of activity predictions. The Mach peptides predicted in this study are the most effective sequences for PMO delivery known to date, improving antisense activity by up to 50-fold while remaining nontoxic.

The interpretability of the model is an additional advantage. By overlaying the output of the predictor with the sequence matrix of a given peptide, we can visualize the activated residues and substructures important for the decision-making process. Several observations from the interpretations match our current understanding of CPP motifs, such as the benefit of cationic residues. The model also identified Ahx as an important residue, one which has only been investigated in the context of endosomal escape in Arg-rich sequences.(*42*)

In addition to PMO, Mach peptides deliver other antisense oligonucleotides as well as functional proteins into the cell cytosol. Delivery of EGFP reveals diffuse green fluorescence in the cytosol and clear accumulation of EGFP to the nucleus. We believe that Mach peptides may contain nuclear localization sequences (NLS), which have been described previously and are typically lysine-rich.(*43*) Nuclear localization is not solely due to the cationic charge, as shown by a previous study.(*44*) The model likely selected for such NLS sequences because the activity used in the training was acquired from a nuclear delivery-based assay.

While in vitro delivery efficiency is important, a greater challenge remains toward in vivo delivery to target tissues. In Duchenne muscular dystrophy, PMO must access the nucleus of muscle cells to have therapeutic effect. Targeting to cardiac tissue is a primary concern given that the leading cause of death from this disease is heart failure. PMO-Mach conjugates effected a dose-dependent increase in protein synthesis in all three examined muscle tissues including heart after a single i.v. injection. A remaining gap in this field is the translation of in vitro experiments to in vivo applications. More sophisticated experimental methods, such as organoids, could improve predictions of therapeutic materials.

This method illustrates how deep learning can be applied to de novo design of functional abiotic peptides. Our machine learning framework is application agnostic and can be repurposed to discover different sequence-optimized polymers with other desired activities. The only limitations lie in the quality of the input dataset and the sequence and activity space it accesses. The resulting Mach peptides are the most effective PMO delivery constructs developed to date and are effective for use in animals. We envision that this strategy will greatly impact the future design of functional polymers.

## Acknowledgments

We thank Alexander R. Loftis and Jacob Rodriguez for assistance with recombinant protein expression, Dr. Coralie Backlund for assistance with immunoassays, Wendy C. Salmon at the W. M. Keck Microscopy Facility at the Whitehead Institute for help with imaging, and Bryan Mastis and Same Foley for help with in vivo studies at Sarepta Therapeutics. We also thank Zi-Ning Choo for igniting our interest in machine learning.

## Author contributions

C.K.S., S.M., J.M.W., B.L.P., and R.G.B. conceptualized the research; J.M.W. and C.M.F. synthesized and tested the modular library; S.M. and R.G.B. developed the machine learning model with input from C.K.S. and B.L.P.; C.K.S. synthesized Mach peptides and constructs, performed experiments and analyzed the results. K.B., C.L.W., and J.A.W. performed the in vivo studies with input from A.B.M.; C.K.S., S.M., A.L., B.L.P., and R.G.B. wrote the manuscript with input from all authors.

## Data and materials availability

Computer code, training data, and trained models are available at https://github.com/learningmatter-mit/peptimizer. All other data is available in the main text or the supplementary materials.

## Supplementary Materials

Materials and Methods

Supplementary Text

Figs. S1 to S17

Tables S1 to S3

References 45-52

