## Supplementary materials for "Interpretable Deep Learning for De Novo Design of Cell-Penetrating Abiotic Polymers"

**This PDF file includes:**

Materials and Methods
Supplementary Text
Figs. S1 to S17
Tables S1 to S3

### 2 Materials and General Methods

#### 2.1 Reagents and Solvents

H-Rink Amide-ChemMatrix resin was obtained from PCAS BioMatrix Inc. (Saint-Jean-sur-Richelieu, Canada). 1-[Bis(dimethylamino)methylene]-1H-1,2,3-triazolo[4,5-b]pyridinium-3-oxid-hexafluorophosphate (HATU), 4-pentynoic acid, 5-azidopentanoic acid, Fmoc-β-Ala-OH, Fmoc-6-aminohexanoic acid, and Fmoc-L-Lys($N_3$) were purchased from Chem-Impex International (Wood Dale, IL). PyAOP was purchased from P3 BioSystems (Louisville, KY). Fmoc-protected amino acids (Fmoc-Ala-OHx$H_2O$, Fmoc-Arg(Pbf)-OH; Fmoc-Asn(Trt)-OH; Fmoc-Asp-(O*t*-Bu)-OH; Fmoc-Cys(Trt)-OH; Fmoc-Gln(Trt)-OH; Fmoc-Glu(O*t*-Bu)-OH; Fmoc-Gly-OH; Fmoc-His(Trt)-OH; Fmoc-Ile-OH; Fmoc-Leu-OH; Fmoc-Lys(Boc)-OH; Fmoc-Met-OH; Fmoc-Phe-OH; Fmoc-Pro-OH; Fmoc-Ser(But)-OH; Fmoc-Thr(*t*-Bu)-OH; Fmoc-Trp(Boc)-OH; Fmoc-Tyr(*t*-Bu)-OH; Fmoc-Val-OH), were purchased from the Novabiochem-line from Millipore Sigma. Peptide synthesis-grade *N,N*-dimethylformamide (DMF), $CH_2Cl_2$, diethyl ether, and HPLC-grade acetonitrile were obtained from VWR International (Radnor, PA). All other reagents were purchased from Sigma-Aldrich (St. Louis, MO). Milli-Q water was used exclusively in all experiments.

#### 2.2 Liquid chromatography-mass spectrometry

LC-MS analyses were performed on either an Agilent 6520 Accurate-Mass Q-TOF LC-MS (abbreviated as 6520) or an Agilent 6550 iFunnel Q-TOF LC-MS system (abbreviated as 6550) coupled to an Agilent 1260 Infinity HPLC system. Mobile phases were: 0.1% formic acid in water (solvent A) and 0.1% formic acid in acetonitrile (solvent B). The following LC-MS methods were used for characterization:

**Method A: 1-61% B over 9 min, Zorbax C3 column (6520)**

<u>LC</u>: Zorbax 300SB-C3 column: 2.1 × 150 mm, 5 μm, column temperature: 40 °C, gradient: 0-2 min 1% B, 2-11 min 1-61% B, 11-12 min 61-95% B, 12-15 min 95% B; flow rate: 0.8 mL/min.

<u>MS</u>: Positive electrospray ionization (ESI) extended dynamic range mode in mass range 300–3000 *m*/*z*. MS is on from 4 to 11 min.

**Method B: 1-61% B over 10 min, Phenomenex Jupiter C4 column (6550)**

LC: Phenomenex Jupiter C4 column: 1.0 × 150 mm, 5 μm, column temperature: 40 °C, gradient: 0-2 min 1% B, 2-12 min 1-61% B, 12-16 min 61-90% B; 16-20 min 90% B; flow rate: 0.1 mL/min.

MS: Positive electrospray ionization (ESI) extended dynamic range mode in mass range 100–1700 *m*/*z*. MS is on from 4 to 12 min.

**Method C: 1-61% B over 10 min, Agilent EclipsePlus C18 column (6550)**

LC: Agilent EclipsePlus C18 RRHD column: 2.1 × 50 mm, 1.8 μm, column temperature: 40 °C, gradient: 0-2 min 1% B, 2-12 min, 1-61% B, 12-13 min, 61% B, 13-16 min, 1% B; flow rate: 0.1 mL/min.

MS: Positive electrospray ionization (ESI) extended dynamic range mode in mass range 300–3000 m/z. MS is on from 4 to 12 min. This method was used exclusively for characterization of the modular library.

All data were processed using Agilent MassHunter software package. Y-axis in all chromatograms shown represents total ion current (TIC) unless noted.

#### 2.3 General method for peptide preparation

Automated Fast-flow Peptide Synthesis: Peptides were synthesized on a 0.1 mmol scale using an automated fast-flow peptide synthesizer.(*38*) A 100 mg portion of ChemMatrix Rink Amide HYR resin was loaded into a reactor maintained at 90 ºC. All reagents were flowed at 40 mL/min with HPLC pumps through a stainless-steel loop maintained at 90 ºC before introduction into the reactor. For each coupling, 10 mL of a solution containing 0.4 M amino acid and 0.38 M HATU in DMF were mixed with 600 μL diisopropylethylamine and delivered to the reactor. Fmoc removal was accomplished using 10.4 mL of 20% (v/v) piperidine. Between each step, DMF (15 mL) was used to wash out the reactor. For peptides in the modular library, special coupling conditions were used for arginine, in which 10 mL of a solution containing 0.4 M Fmoc-L-Arg(Pbf)-OH and 0.38 M PyAOP in DMF were mixed with 600 μL diisopropylethylamine and delivered to the reactor. For Mach peptides, additional special coupling conditions were used according to the optimized peptide synthesis protocol.(*45*) To couple unnatural amino acids or to cap the peptide (e.g., with 4-pentynoic acid), the resin was incubated for 30 min at room temperature with 4-pentynoic acid (1 mmol) dissolved in 2.5 mL 0.4 M HATU in DMF with 500 μL diisopropylethylamine. After completion of the synthesis, the resin was washed 3 times with DCM and dried under vacuum.

Peptide Cleavage and Deprotection: Each peptide was subjected to simultaneous global side-chain deprotection and cleavage from resin by treatment with 5 mL of 94% trifluoroacetic acid (TFA), 2.5% 1,2-ethanedithiol (EDT), 2.5% water, and 1% triisopropylsilane (TIPS) (v/v) for 7 min at 60 °C or at room temperature for 2 to 4 h. For arginine-rich sequences, the resin was treated with a cleavage cocktail consisting of 82.5% TFA, 5% phenol, 5% thioanisole, 5% water, and 2.5% EDT (v/v) for 14 h at room temperature. For peptides containing azide, EDT was substituted for thioanisole. The cleavage cocktail was first concentrated by bubbling $N_2$ through the mixture, and cleaved peptide was precipitated and triturated with 40 mL of cold ether (chilled in dry ice). The crude product was pelleted by centrifugation for three minutes at 4,000 rpm and the ether was decanted. This wash step was repeated two more times. After the third wash, the pellet was dissolved in 50% water and 50% acetonitrile containing 0.1% TFA, filtered through a fritted syringe to remove the resin and lyophilized.

Peptide Purification: The peptides were dissolved in water and acetonitrile containing 0.1% TFA, filtered through a 0.22 μm nylon filter and purified by mass-directed semi-preparative reverse-phase HPLC. Solvent A was water with 0.1% TFA additive and Solvent B was acetonitrile with 0.1% TFA additive. A linear gradient that changed at a rate of 0.5% B/min was used. Most of the peptides were purified on an Agilent Zorbax SB C3 column: 9.4 × 250 mm, 5 μm. Extremely hydrophilic peptides, such as the arginine-rich sequences were purified on an Agilent Zorbax SB C18 column: 9.4 × 250 mm, 5 μm. Using mass data about each fraction from the instrument, only pure fractions were pooled and lyophilized. The purity of the fraction pool was confirmed by LC-MS.

##### 2.4 PMO-DBCO Synthesis

PMO IVS-654 (50 mg, 8 μmol) was dissolved in 150 μL DMSO. To the solution was added a solution containing 2 equivalents of dibenzocyclooctyne acid (5.3 mg, 16 μmol) activated with HBTU (37.5μL of 0.4 M HBTU in DMF, 15 μmol) and DIEA (2.8 μL, 16 μmol) in 40 μL DMF (Final reaction volume = 0.23 mL). The reaction proceeded for 25 min before being quenched with 1 mL of water and 2 mL of ammonium hydroxide. The ammonium hydroxide hydrolyzed any ester formed during the course of the reaction. After 1 h, the solution was diluted to 40 mL in water/acetonitrile and purified using reverse-phase HPLC (Agilent Zorbax SB C3 column: 21.2 × 100 mm, 5 μm) and a linear gradient from 2 to 60% B (solvent A: water; solvent B: acetonitrile) over 58 min (1% B / min). Using mass data about each fraction from the instrument, only pure fractions were pooled and lyophilized. The purity of the fraction pool was confirmed by LC-MS (Appendix II).

##### 2.5 Library Synthesis Conditions

The library was synthesized in a combinatorial fashion and analyzed by LC-MS.(*46*) The 600-member library was synthesized using 50 peptide members in module 4 (Table S1).

Reaction 1: PMO-DBCO was dissolved in water to 10 mM concentration (determined by UV-Vis). The module 2 peptides were dissolved in water containing 0.1% trifluoroacetic acid at 10 mM concentration (determined gravimetrically; the molecular weight was calculated to include 0.5 trifluoroacetate counter ions per lysine, arginine, and histidine residue). In a microcentrifuge tube, 50 μL each of PMO-DBCO solution and module 2 peptide solution were mixed and incubated for 1 h. The product was analyzed by LC-MS and dried by lyophilization. Lastly, the product was resuspended in 100 μL of DMSO to provide a 5 mM solution and stored at −20 ºC.

Reaction 2: Stock solutions were prepared by dissolving module 3 peptides and module 4 peptides in water at 10 mM concentration (determined gravimetrically). For each reaction, 4 μL of module 3 peptide was mixed with 4 μL of module 4 peptide in a PCR tube. Separately, the copper bromide solution was prepared by mixing 1 mL of degassed DMSO with 2.8 mg copper(I) bromide under $N_2$ to afford a 20 mM solution. Under ambient conditions, 4 μL of the CuBr solution was added to the mixture of module peptides 3 and 4. The reaction was capped and the reaction was allowed to proceed for 2 h; the small amount of $O_2$ present during reaction setup does not substantially impede reaction progress. After 2 h, 2 μL of a 100 mM solution of $Na_2HPO_4$ was added. The PCR tube was then sonicated, vortexed, and centrifuged. To remove the solvent, the PCR tube was centrifuged under vacuum using a Savant SPD121P Speed-Vac set at 35 ºC for 2 h. Lastly, the product was resuspended in 16 μL of DMSO to provide a 5 mM solution and stored at −80 ºC. The product was analyzed by LC-MS.

Reaction 3: The final modular construct was synthesized through the combination of module 1-2 and module 3-4. First, 1.6 μL of reaction 2 was added to a 384-well plate. Separately, 30 μL of reaction 1 was mixed with 15 µL of TCEP solution (100 mM TCEP·HCl in 50/50 water/DMSO containing 400 mM NaOH) and 75 µL DMSO. Then, 1.6 µL of the reaction 1 solution was added to reaction 2 in the 384 well plate. Each individual reaction ultimately contained 0.4 µL of reaction 1 (at 5 mM in DMSO), 1.6 µL of reaction 2 (at 5 mM in DMSO), 0.2 µL TCEP solution (at 100 mM in water/DMSO), and 1 µL DMSO. Excess reaction 2 was used to force the reaction to go to completion; the presence of copper hinders the efficiency of this conjugation. Reaction 1 was used as a limiting reagent to avoid excess PMO, which is the active component for the cell culture assays. The reaction was allowed to proceed for 2 h, and then the plate was stored at −80 ºC. The reaction was analyzed by LC-MS.

**2.6 PMO and PNA-Peptide Conjugation**

Mach peptides were conjugated to PMO via strain-promoted azide-alkyne cycloaddition. PMO-DBCO (5 mM in water) was stoichiometrically combined with azide-peptide (5 mM in water) and incubated at room temperature until reaction completed (between 2 and 12 h), monitored by LC-MS. The reaction was purified using reversed-phase HPLC (Agilent Zorbax SB C3 column: 21.2 × 100 mm, 5 µm) and a linear gradient from 2 to 60% B (solvent A: 100 mM ammonium acetate in water pH 7.2; solvent B: acetonitrile) over 58 min (1% B / min). pure fractions were pooled as determined by LC-MS and lyophilized (Appendix II).

50 nmol of PNA 654 (O-GCTATTACCTTAACCCAG-Lys(DBCO)) was purchased from PNABio. PNA-DBCO (1 mM in water) was stoichiometrically combined with azide-peptide (1 mM in water) and incubated at 4 °C for 12 h. The product was then used in cell assays without purification. Conversion was checked by LC-MS (Appendix II).

**2.7 HeLa-654 eGFP Assay**

HeLa 654 cells obtained from the University of North Carolina Tissue Culture Core facility were maintained in MEM supplemented with 10% (v/v) fetal bovine serum (FBS) and 1% (v/v) penicillin-streptomycin at 37 °C and 5% $CO_2$. 18 h prior to treatment, the cells were plated at a density of 5,000 cells per well in a 96-well plate in MEM supplemented with 10% FBS and 1% penicillin-streptomycin.

For testing of the library, on the day of the experiment, the 384 well plate containing the crude reaction mixtures in DMSO was diluted to 100 μM by the addition of 16.8 μL of PBS to the 3.2 μL reaction mixture. Then, each construct was diluted to 5 μM in MEM supplemented with 10% FBS and 1% penicillin-streptomycin. For individual peptide testing, PMO-peptides were dissolved in PBS without $Ca^{2+}$ or $Mg^{2+}$ at a concentration of 1 mM (determined by UV) before being diluted in MEM. Cells were incubated at the designated concentrations for 22 h at 37 °C and 5% $CO_2$. Next, the treatment media was removed, and the cells were washed once before being incubated with 0.25 % Trypsin-EDTA for 15 min at 37 °C and 5% $CO_2$. Lifted cells were transferred to a V-bottom 96-well plate and washed once with PBS, before being resuspended in PBS containing 2% FBS and 2 μg/mL propidium iodide (PI). Flow cytometry analysis was carried out on a BD LSRII flow cytometer at the MIT Koch Institute. Gates were applied to the data to ensure that cells that were positive for propidium iodide or had forward/side scatter readings that were sufficiently different from the main cell population were excluded. Each sample was capped at 5,000 gated events (Fig. S1).

Analysis was conducted using Graphpad Prism 7 and FlowJo. For each sample, the mean fluorescence intensity (MFI) and the number of gated cells was measured. To report activity, triplicate MFI values were averaged and normalized to the PMO alone condition.

#### 3 Development of the machine learning model

##### 3.1 Training of Generator

The generator is a data-driven tool to generate new peptide sequences that follow the ontology of cell penetrating peptides, and is based on recurrent neural network (RNN) - Nested LSTM architecture.(*34*) It was trained using one-hot encoding representations of the amino acids. The training dataset comprised of 1150 sequences, including unique (non-modular) sequences used in the creation of the library and sequences from CPPSite2.0.(*35*) The training was performed using 80% of this dataset, and validated using the remaining 20%. A validation accuracy of 76% was obtained in the training. For the model, multiple combinations of Nested LSTM and vanilla-LSTM layers were tried with different cell sizes.(*34*) The final model was chosen after the optimization of hyperparameters. All hyperparameters were optimized using SigOpt.(*47*)

##### 3.2 Training of Predictor

The predictor, based on convolutional neural network (CNN), estimates the normalized fluorescence intensity from PMO delivery by a given peptide sequence, as measured in the HeLa 654 assay. The model was trained on a linear graph of peptide sequences with a row matrix of residue fingerprints. The representation of the peptide sequences for the Conv1D model is similar to an image with 1 color channel. The row matrix of 2048-bit vectors (vector of 0s and 1s) represents the arrangement of the residues along the backbone of the peptide chain. This is analogous to a 2D black and white image. All fingerprints were generated using RDKit.(*48*) By combining the CPP library from this work as well as the collection of CPPs from previous work, we compiled 640 PMO-peptides sequences for training.(*6*) All hyperparameters were optimized using SigOpt.(*47*) We used fingerprints and one-hot encodings to train non-CNN models such as those based on support vector regression, Gaussian process regression, kernel ridge regression, k-nearest neighbors regression and XGBoost regression.

##### 3.3 Setup of Optimizer

The optimization was done using genetic algorithm (GA), where single residue mutations involved insertion, deletion and swapping, and multi-residue mutation was done using hybridization. Single residue mutation involved choosing the index of the residue, and deleting, or in the case of insertion/swapping, adding another residue, with all the processes being random. For hybridization, the sequence length and position to be hybridized, and the hybridized sequence (from the list of all CPPs) were all chosen randomly. In the case of hybridization mutation, the selection and replacement of motifs was done at random without conservation of the sequence length. For the case of mutations with cysteine macrocycles, explicit conditions were built in to keep the number and position of cysteine residues separate in the case of a single through-space covalent bond or bicycle. A constrained hybridization condition conserving the sequence length was also set-up for specific optimization tasks. In the case of cysteine macrocycles, different fingerprints were used to denote the residues. The GA was implemented for the following objective function for all LSTM generated sequences for 1000 evolution steps:

$$GA\ Score = \frac{1}{2}\ Intensity - \frac{1}{2}\left(\frac{1}{2}\ R_{count} + \frac{1}{5}\ Length - \frac{1}{10}\ Net\ Charge + Similarity\right)$$

##### 3.4 Set-up of Generator-Predictor-Optimizer Loop

The generator was seeded with random sequences from the training dataset. After multiple rounds of sampling, the randomly sampled sequences were set-up for optimization. The directed evolution of the generated sequences was carried using the predictor-optimizer feedback loop. Each sequence was mutated by the optimizer. Post mutations, the normalized fluorescence values for the new sequence was predicted by the Predictor and the optimization parameters (% arginine, length, net charge) were calculated. The objective function (equation with optimization parameters) was evaluated for both the old and mutated sequences. If the value for the mutated sequence was higher for the mutated than the older, then the old sequence was replaced by the mutated sequence. 1000 such optimization rounds were conducted for each sequence. The output was hundreds of sequences with varying predicted activity.

##### 3.5 Similarity of Training and Validation Sequences

Similarity among sequences in each training (for generator and predictor) and validation dataset was analyzed using Jaro-Winkler distance metric (Fig. S3).(*49*) Each sequence was compared with the rest of the library to evaluate the string similarity.

For the sequences used to train the generator, it is observed that the sequences have a mean similarity of 47% enabling us to capture a combinatorial chemical space of cell-penetrating peptide sequences. For the sequences used to train the predictor, the modularity of the sequences based on the combinatorial library is evident in the visualization of sequence similarity. The four highlighted squares along the diagonal correspond to module 2 of the sequences. Similarly, the four lighter colored boxes correspond to module 3. The non-modular sequences, which are dissimilar from one another, are on the bottom of the visualization. The mean similarity of the sequences is 66%.

Similarity of Mach sequences was first compared to the library using mean Jaro-Winkler distance (Fig. S3). All Mach sequences had a mean similarity less than 60% when compared to the training dataset. Then to compare Mach peptides to the existing proteome, we used BLASTp on the online server.(*50*) The search was done using default values to search the UniProt database. There was no sequence homology between Mach sequences and known proteins for significant E-values less than 0.01. For the unnatural residues, B (β-Alanine) and X (Aminohexanoic acid) were replaced by A (alanine) and L (leucine) respectively for the search operation. Mach sequences containing cysteine macrocycles were excluded from the search.

##### 3.6 Immunogenicity of Validation Sequences

The likelihood of being a T-cell epitope was calculated for all sequences using an online server (Fig. S17).(*51*) The score is an arbitrary number, where a higher positive value indicates a higher probability of the peptide to be immunogenic and vice-versa. For the unnatural residues, B (β-Alanine) and X (Aminohexanoic acid) were replaced by A (Alanine) and L (Leucine) respectively for the search operation. The Mach sequences were compared to the sequences used in the training of the predictor.

##### 3.7 Interpretability of Training Dataset using Conv1D Model

Using the conceptual framework used to understand activation of neural networks for image classification, we developed a toolkit to visualize the decision making process of our model.(*37*) This framework, in the case of images, allows users to understand which regions activated the decision making process of the neural network both in a positive and negative context. For instance, in a cat image, the whiskers positively activate, while the wall paint in the background negatively activate the decision engine of the machine learning model.(*37*)

We chose the first convolution layer of the model to access the fingerprint indices. Taking the gradient, first-order differential, of the model output (normalized fluorescence intensity) with respect to the input representation (row matrix of fingerprints), we were able to get the positive and negative activation values element-wise for each fingerprint index (representing chemical substructure) and by how much they affected the prediction of the model. To separate the analysis of positive and negative gradients, we multiplied the respective Jacobian, i.e., gradients, with the activated fingerprints (1s in the matrix) individually. A comprehensive visualization was obtained by individually averaging over the residue positions and fingerprint indices, and obtaining the particular substructures in individual amino acids (Fig. S4 and S5).

### 4 Evaluation of PMO-Mach constructs

#### 4.1 LDH release Assay

Cytotoxicity assays were performed in both HeLa 654 cells and human RPTEC (Human Renal Proximal Tubule Epithelial cells, TH-1, ECH001, Kerafast). RPTEC were maintained in high glucose DMEM supplemented with 10% (v/v) fetal bovine serum (FBS) and 1% (v/v) penicillin-streptomycin at 37 °C and 5% $CO_2$. Treatment of RPTEC was performed as with the HeLa 654 cells. After treatment, supernatant was transferred to a new 96-well plate. To each well of the 96-well plate containing supernatant, described above, was added CytoTox 96 Reagent (Promega). The plate was shielded from light and incubated at room temperature for 30 min. Equal volume of Stop Solution was added to each well, mixed, and the absorbance of each well was measured at 490 nm. The blank measurement was subtracted from each measurement, and % LDH release was calculated as: % cytotoxicity = 100 × Experimental LDH Release (OD490) / Maximum LDH Release (OD490).

#### 4.2 Inflammation panel

The inflammatory response to treatment was assayed by profiling inflammatory cytokine release by THP-1-derived macrophages. THP-1 cells (ATCC TIB-202) were grown in RPMI 1640 media supplemented with 10% (v/v) FBS, 1% (v/v) penicillin-streptomycin, L-glutamine, non-essential amino acids, sodium pyruvate at 37 °C and 5% $CO_2$. Two days before the experiment, THP-1 cells (~450,000/mL) were treated with 25 nM phorbol 12-myristate 13-acetate (PMA) at 37 °C and 5% $CO_2$ for 24 h to trigger differentiation into macrophages. Then, media was replaced with fresh RPMI media and the cells were incubated for another 24 h. At this time the phenotype changed from suspension cells to strongly adherent cells. In the morning of the experiment, the supernatant was removed and macrophages were lifted by incubating in enzyme-free cell dissociation buffer (Thermo) for 5 min. Cells were then collected, spun down, and brought up in complete RPMI media to a cell density of ~500,000/mL. ~100,000 cells were plated in each well of a 96-well plate, leaving the first two columns empty. Cells were allowed to re-adhere before treatment. Duplicate wells were treated with varying concentrations of the PMO-peptide conjugates at 37 °C and 5% $CO_2$ for 2 h. Media-only and no treatment wells were used as negative controls, and 10 ug/mL bacterial lipopolysaccharide (LPS) treatment was used as a positive control. Following treatment, each well was washed three times, given fresh media, and incubated for 12 h. Supernatant was transferred to a V-bottom plate and spun down at 4000 rcf to remove debris. Inflammatory cytokines in the supernatant were assayed using LEGENDplex Human Inflammation panel (BioLegend), a fluorescent bead-based immunoassay. Cytokines assayed were: IL-1β, IFN-α2, IFN-γ, TFN-α, MCP-1, IL-6, IL-8, IL-10, IL-12p70, IL-17A, IL-18, IL-23, and IL-33. Analysis was carried out on a BD LSRII flow cytometer and data was analyzed using BioLegend's accompanying software.

##### 4.3 Endocytosis Inhibition Assays

Chemical endocytosis inhibitors were used to probe the mechanism of delivery of PMO and DTA in a pulse-chase format. For the PMO constructs, HeLa 654 cells were preincubated with various chemical inhibitors or incubated at 4 °C for 30 min before treatment with PMO-Mach constructs for three hours. Treatment media was then replaced with fresh media for 22 h. Cells were then lifted as previously described and GFP synthesis was measured by flow cytometry (Fig. S9).

For the DTA constructs, HeLa cells were preincubated with the same chemical endocytosis inhibitors for 30 min before treatment with DTA constructs for 2 h. Treatment media was then replaced with fresh media, and the plate was incubated at 37 °C for 48 h. Cell proliferation was determined using the CellTiter-Glo assay.

##### 4.4 Circular Dichroism

Peptides were dissolved in PBS buffer to obtain stock solutions of 1 mM. The circular dichroism (CD) spectra was obtained from 195 to 250 nm using an AVIV 420 circular dichroism spectrometer with a 1 mm path length quartz cuvette. Peptides in PBS buffer at 20 μM, with or without 10 mM sodium dodecyl sulfate (SDS) were used in the measurement.

### 5 Delivery of other biomacromolecules

#### 5.1 Recombinant expression

$His_6$-SUMO-$G_5$-DTA(C186S), $His_6$-SUMO-$G_5$-DTA(C186S, E148S) and $G_5$-eGFP-$His_6$ were overexpressed in *E. coli* BL21 (DE3) cells. Approximately 10 g of cell pellet was lysed by sonication in 50 mL of 20 mM Tris, 150 mM NaCl, pH 7.5 buffer containing 30 mg lysozyme, 2 mg DNAase I, and 1 tablet of cOmplete™ Protease Inhibitor Cocktail. The suspension was centrifuged at 16,000 rpm for 30 min to remove cell debris. The supernatant was loaded onto a 5 mL HisTrap FF Ni-NTA column (GE Healthcare, UK) and washed with 30 mL of 100 mM imidazole in 20 mM Tris, 150 mM NaCl, pH 8.5. Protein was eluted from the column with buffer containing 300 mM imidazole in 20 mM Tris, 150 mM NaCl, pH 8.5. Imidazole was removed from protein via centrifugation in Millipore centrifugal filter unit (10K).

For the DTA constructs, the $His_6$-SUMO tag was then cleaved from the protein with SUMO protease (previously recombinantly expressed) by incubating a 1:1000 protease:protein ratio in 20 mM Tris, 150 mM NaCl, pH 7.5 overnight at 4 °C. Desired protein was separated from $His_6$-SUMO tag by flowing the mixture through a 5 mL HisTrap FF Ni-NTA column. Finally, purified protein was isolated by size exclusion chromatography using HiLoad 26/600 Superdex 200 prep grade size exclusion chromatography column (GE Healthcare, UK) in 20 mM Tris, 150 mM NaCl, pH 7.5 buffer.

For the GFP construct, purified protein was isolated by anion exchange chromatography using HiTrap Q HP anion exchange chromatography column (GE Healthcare, UK) in (0-40% B over 20 CV) where A: 20 mM Tris, pH 8.5 buffer and B: 1 M NaCl, 20 mM Tris, pH 8.5 buffer.

Proteins were analyzed using SDS-PAGE gel. In addition, proteins were analyzed by ESI-Q-TOF LC-MS to confirm molecular weight and purity. The protein charge-state envelope was deconvoluted using Agilent MassHunter Bioconfirm using maximum entropy (Appendix II).

#### 5.2 Synthesis and Testing of Mach-DTA

Mach-LPSTGG peptides were synthesized and purified by standard protocol as described. $G_5$-DTA (50 μM) was incubated with either Mach3-LPSTGG (250 μM) or Mach7-LPSTGG (750 μM) and SrtA* (2.5 μM) for 90 min at 4 °C in SrtA buffer (10 mM $CaCl_2$, 50 mM Tris, 150 mM

NaCl, pH 7.5). The reaction was monitored by LCMS and gel electrophoresis. After 90 min, Mach-DTA conjugate was isolated using HiLoad 26/600 Superdex 200 prep grade size exclusion chromatography column (GE Healthcare, UK) in 20 mM Tris, 150 mM NaCl, pH 7.5 buffer. Fractions containing the pure product as determined by LC-MS and gel electrophoresis were concentrated using a centrifugal filter unit (10K, Millipore) (Appendix II).

To test for DTA delivery to the cytosol, HeLa cells were plated at 5,000 cells/well in a 96-well plate the day before the experiment. Wild-type and mutant constructs of $G_5$-DTA, Mach3-DTA, and Mach7-DTA, as well as Mach3-LPSTGG and Mach7-LPSTGG were prepared at varying concentrations in complete media and transferred to the plate. Cell proliferation was measured after 48 h using the CellTiter-Glo assay.

##### 5.3 Synthesis and Testing of Mach-eGFP

$G_5$-EGFP (60 μM) was incubated with either Mach3-LPSTGG (1000 μM) or Mach7-LPSTGG (1000 μM) and SrtA* (5 μM) in SrtA buffer (10 mM $CaCl_2$, 50 mM Tris, 150 mM NaCl, pH 7.5) for 90 min at room temperature under exclusion of light. The reaction was monitored by LCMS and gel electrophoresis. After 90 min, Mach-eGFP conjugate was isolated using cation exchange chromatography using HiTrap SP HP cation exchange chromatography column (GE Healthcare, UK) in (0-100% B over 20 CV) where A: 50 mM NaCl, 20 mM Tris, pH 7.5 buffer and B: 1 M NaCl, 20 mM Tris, pH 12 buffer. Fractions containing the pure product as determined by LC-MS and gel electrophoresis were immediately desalted and concentrated using a centrifugal filter unit (10K, Millipore) (Appendix II).

To visualize delivery of EGFP into Hela cells, HeLa cells were plated at 5,000 cells/well in a coverslip glass-bottomed 96-well plate the day before the experiment. Mach3-EGFP, Mach7-EGFP, or EGFP at varying concentrations were added to each well and incubated at 37 °C and 5% $CO_2$ for 1 h. Treatment media was replaced with fresh media 1 h before being imaged in the W.M. Keck microscopy facility on an RPI Spinning Disk Confocal microscope on brightfield and GFP setting (488 nm 150 mW OPSL excitation laser, 525/50 nm emission).

#### 6 In Vivo Studies

EGFP-654 transgenic mice were first obtained from Dr. Ryszard Kole's laboratory.(*52*) This mouse model ubiquitously expresses EGFP-654 transgene throughout body under chicken β-actin promoter. A mutated nucleotide 654 at intron 2 of human β-globin gene is contained in the EGFP-654 sequence which interrupts EGFP-654 coding sequence and prevents proper translation of EGFP protein. The antisense activity of PMO blocks aberrant splicing and resulted in EGFP expression, the same as in the HeLa 654 assay. In this study, 6- to 8-week-old male EGFP-654 mice bred at Charles River Laboratory were shipped to the vivarium at Sarepta Therapeutics (Cambridge, MA). These mice were group housed with ad libitum access to food and water. All animal protocols were approved by the Institutional Animal Care and Use Committee (IACUC) of Sarepta Therapeutics.

After 3-days of acclimation, mice were randomized into groups to receive a single *i.v.* tail vein injection of either saline or PMO-peptide (PMO-Mach3 or PMO-Mach4) at the indicated doses; 5, 10 and 30 mg/kg. Seven days after the injection, the mice were euthanized for serum and tissue sample collection. Quadriceps, diaphragm, heart were rapidly dissected, snap-frozen in liquid nitrogen and stored at -80 °C until analysis.

Serum from all groups were collected 7-days post-injection and tested for kidney injury markers using a Vet Axcel Clinical Chemistry System (Alfa Wassermann Diagnostic Technologies, LLC). Specifically, serum BUN, creatinine, and cystatin C levels were measured using ACE® Creatinine Reagent (Alfa Wassermann, Cat# SA1012), ACE® Blood Urea Nitrogen Reagent (Alfa Wassermann, Cat# SA2024) and Diazyme Cystatin C immunoassay (Diazyme Laboratories, Cat# DX133C-K), respectively, per manufacturer's recommendation.

20-25 mg of mouse tissue was homogenized in RIPA buffer (Thermo Fisher, Cat# 89900) with protease inhibitor cocktail (Roche, 04693124001) using a Fast Prep 24-5G instrument (MP Biomedical). Homogenates were centrifuged at 12,000 g for 10 min at 4 °C. The resultant supernatant lysates were quantified by Pierce BCA Protein Assay Kit (Thermo Fisher, Cat# 23225) and saved for EGFP expression measurement. Specifically, 80 µg of lysates were aliquoted in each well in a black-wall clear-bottom 96-well microplate (Corning). EGFP fluorescent intensity of each sample was measured in duplicates using a SpectraMAx i3x microplate reader (Molecular devices) by default setting. The average EGFP fluorescent intensity of each sample was then plotted against a standard curve constructed by recombinant EGFP protein (Origen, Cat#TP790050) to quantify EGFP protein level per µg protein lysate.

#### Appendix I: Supplementary Figures

**Table S1: List of peptides used for each module in the 600-member library.** Z refers to 4-pentynoyl. "**C**" refers to cysteines that are linked with decafluorobiphenyl. "*C*" refers to cysteines that are linked with 1,3,5-trisbromomethylbenzene. Module 4 included fifty CPPs, including a mixture of chimeric peptides, cyclic peptides, and bicyclic peptides that we have previously reported to improve PMO delivery.

| Module 1 | Name | Sequence |
|---|---|---|
| | PMO IVS2-654 | GCT ATT ACC TTA ACC CAG |
| **Module 2** | **Name** | **Sequence** |
| | Penetratin | RQIKIWFQNRRMKWKK |
| | pVec | LLIILRRRIRKQAHAHSK |
| | TP10 | AGYLLGKINLKALAALAKKIL |
| | DPV6 | GRPRESGKKRKRKRLKP |
| **Module 3** | **Name** | **Sequence** |
| | KRVK (NLS) | KRVK |
| | SV40 (NLS) | PKKKRKV |
| | AAV-PHP.eB | SDGTLAVPFKA |
| **Module 4** | **Name** | **Sequence** |
| | DPV6 | ZGRPRESGKKRKRKRLKP |
| | PPC3 | ZKKYRGRKRHPR |
| | PPC5 | ZGRKAARAPGRRKQ |
| | R12 | ZRRRRRRRRRRRR |
| | R12 full cycle | Z**C**RRRRRRRRRRRR**C** |
| | R12 N-cycle | Z**C**RRRRRR**C**RRRRRR |
| | R12 C-cycle | ZRRRRRR**C**RRRRRR**C** |
| | R12 benzyl bicycle | Z*C*RRRRRR*C*RRRRRR*C* |
| | R12 double cycle | Z**C**RRRRRR**CC**RRRRRR**C** |
| | Bpep | ZRXRRBRRXRRBR |
| | Bpep full cycle | Z**C**RXRRBRRXRRBR**C** |
| | Bpep C-cycle | ZRXRRBR**C**RXRRBR**C** |
| | Penetratin (nle) | ZRQIKIWFQNRRMKWKK |
| | Engrailed N-cycle | Z**C**QIKIWF**C**NKRAKIKK |
| | Engrailed C-cycle | ZSQIKIWFQ**C**KRAKIK**C** |
| | Engrailed full cycle | Z**C**SQIKIWFQNKRAKIKK**C** |
| | pVEC | ZLLIILRRRIRKQAHAHSK |
| | pVEC-Bpep | ZLLIILRRRIRKQAHAHSKRXRRBRRXRRBR |
| | AIP6 full cycle | Z**C**RLRWR**C** |
| | Melittin-Bpep | ZGIGAVLKVLTTGLPALISWIKRKRQQRXRRBRRXRRBR |

| | |
|---|---|
| Bh3 helix | ZIWIAQELRRIGDEFNAYYARR |
| Bac7 | ZRRIRPRPPRLPRPRPRPLPFPRPG |
| Buforin 2 | ZTRSSRAGLQWPVGRVHRLLRK |
| Melittin | ZGIGAVLKVLTTGLPALISWIKRKRQQ |
| SynB1 | ZRGGRLSYSRRRFSTSTGR |
| S413-PVrev | ZALWKTLLKKVLKAPKKKRKV |
| Ribotoxin2 L3 | ZKLIKGRTPIKFGKADCDRPPKHSQNGMGK |
| PreS2-TLM | ZPLSSIFSRIGDP |
| MAP | ZKLALKALKALKAALKLA |
| W/R | ZRRWWRRWRR |
| MAP12 | ZLKTLTETLKELTKTLTEL |
| SAP | ZVRLPPPVRLPPPVRLPPP |
| SVM1 | ZFKIYDKKVRTRVVKH |
| SVM3 | ZKGTYKKKLMRIPLKGT |
| SVM4 | ZLYKKGPAKKGRPPLRGWFH |
| YTA4 | ZIAWVKAFIRKLRKGPLG |
| 439a | ZGSPWGLQHHPPRT |
| HoxA13 serine2 | ZRQVTIWSQNRRVKSKK |
| Bip | ZVSALK |
| PPR3 | ZPPRPPRPPR |
| PPR4 | ZPPRPPRPPRPPR |
| AIP6 | ZRLRWR |
| DPV15b | ZGAYDLRRRERQSRLRRRERQSR |
| TAT | ZRKKRRQRRR |
| Penetratin | ZRQIKIWFQNRRMKWKK |
| R9 | ZRRRRRRRRR |
| HoxA13 serine1 | ZRSVTIWFQSRRVKEKK |
| KRVK TP10 | ZKRVKAGYLLGKINLKALAALAKKIL |
| TP10 KRVK | ZAGYLLGKINLKALAALAKKILKRVK |
| SV40 TP10 | ZPKKKRKVAGYLLGKINLKALAALAKKIL |

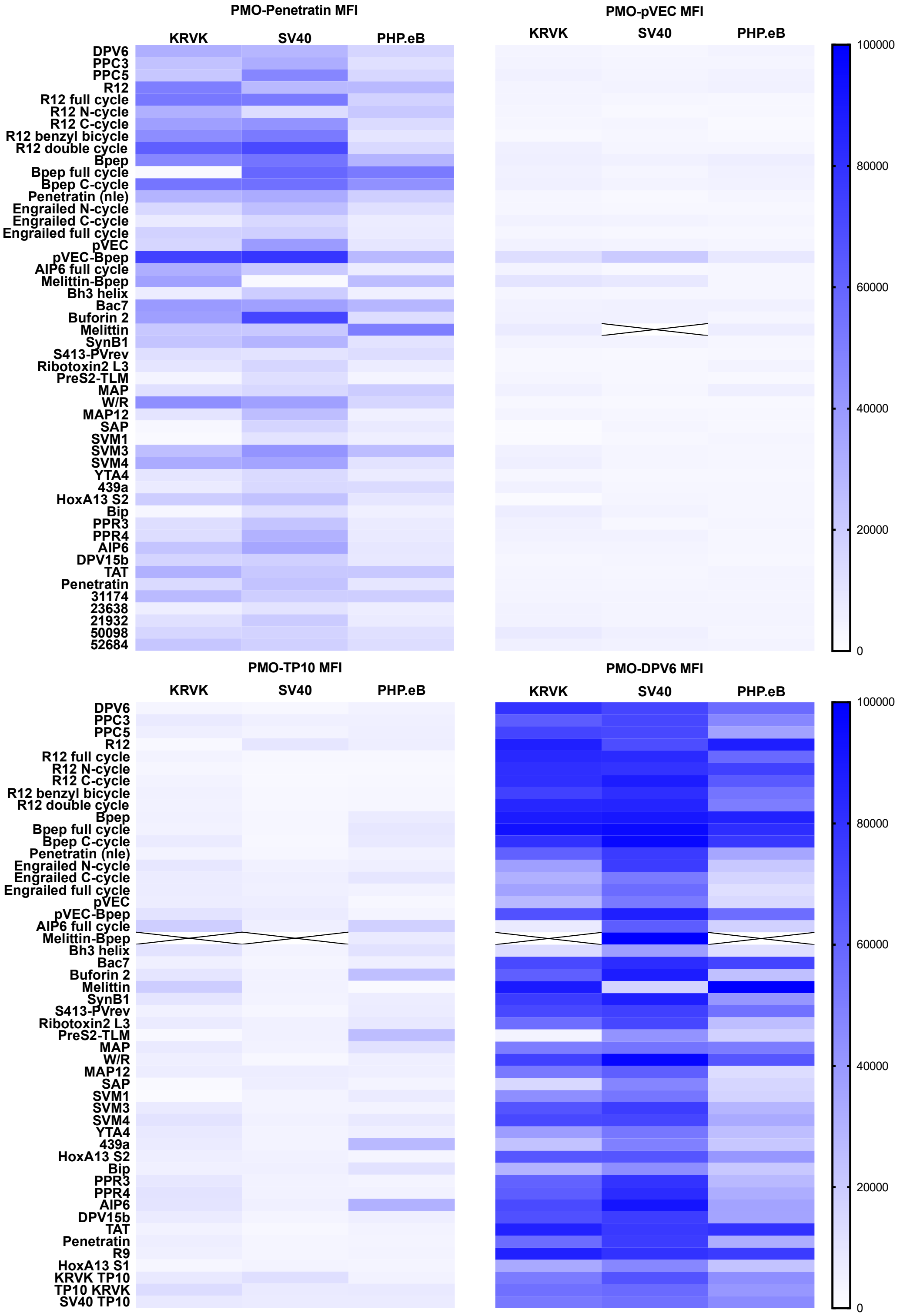

**Fig. S1. Mean fluorescence intensity of the 600-member modular library.** The heat maps show the mean fluorescence intensity of the 600 constructs tested in the HeLa-654 assay (n = 1 replicate well). Boxes marked with an "X" are constructs in which the gated cell count was zero.

**Table S2. Benchmarks for model performance for different representations**. We used fingerprints and one-hot encodings to train non-CNN models such as those based on support vector regression, Gaussian process regression, kernel ridge regression, k-nearest neighbors regression and XGBoost regression.

| Method | Input Feature | Validation Loss (RMSE) | Absolute Error Fold over PMO | % Accuracy* within range of training values | Correlation Coefficient, $R^2$ |
|---|---|---|---|---|---|
| **Conv1D** | **2048-bit FP** | **0.434** | **2.15** | **88.8** | **0.917** |
| Conv1D | 1-hot | 0.461 ± 0.009 | 2.28 | 88.1 | 0.889 |
| Support Vector Regression | 128-bit FP | 1.023 ± 0.005 | 5.08 | 73.5 | -0.031 |
| | 1-hot | 0.764 ± 0.005 | 3.78 | 80.3 | 0.705 |
| Gaussian Process Regression | 128-bit FP | 0.942 ± 0.142 | 5.16 | 73.1 | -0.061 |
| | 1-hot | 0.711 ± 0.020 | 3.45 | 82.0 | 0.741 |
| Kernel Ridge Regression | 128-bit FP | 0.933 ± 0.001 | 8.92 | 53.5 | -0.045 |
| | 1-hot | 0.655 ± 0.033 | 3.36 | 82.5 | 0.775 |
| k-Nearest Neighbors Regression (k=13) | 128-bit FP | 1.082 ± 0.036 | 5.48 | 71.5 | -0.008 |
| | 1-hot | 0.999 ± 0.078 | 5.22 | 72.8 | 0.041 |
| XGBoost Regression (max_depth = 5) | 128-bit FP | 0.683 ± 0.061 | 5.77 | 69.9 | -0.138 |
| | 1-hot | 0.655 ± 0.001 | 3.24 | 83.1 | 0.822 |

* % Accuracy of the model within range of training values, is defined as –

$$\%\, Accuracy = \frac{Absolute\ Error}{Max\ Expt\ Intensity - Min\ Expt\ Intensity} \times 100\%$$

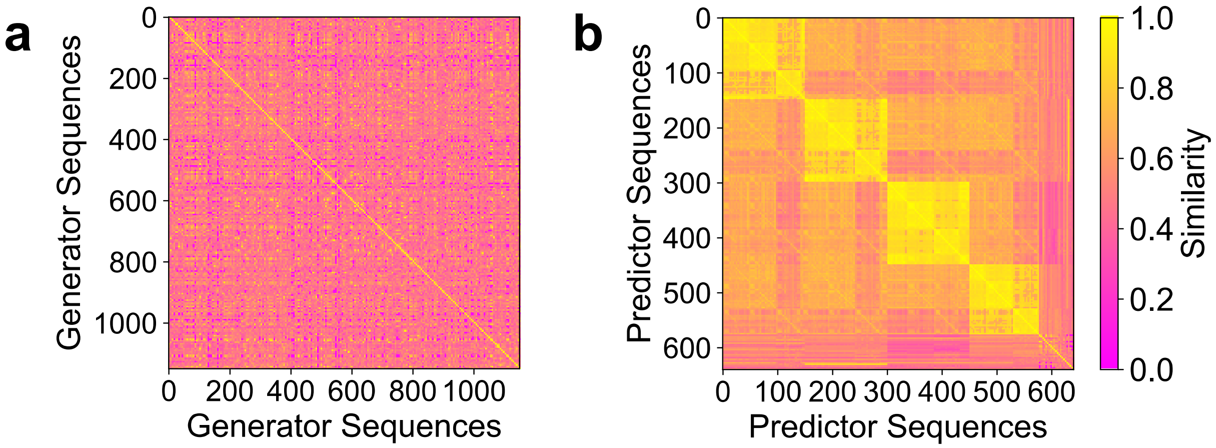

**Fig. S2. Similarity of sequences used in the training of generator and predictor**. Each sequence used in training of **(a)** generator (Nested LSTM) and **(b)** predictor (Convolutional Neural Network based model) is compared with the rest of respective training dataset. The mean similarities of the sequences are 47% and 66% for the generator and predictor respectively. The heatmap for the predictor sequences have a modular pattern owing to the combinatorial nature of the library. Jaro-Winkler distance was used as the metric to assess the similarity between two sequences.

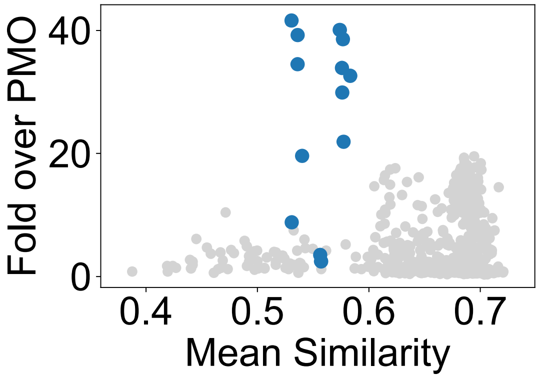

**Fig. S3. Similarity and experimental normalized MFI of Mach and training sequences.** Mach sequences are novel and high-performing in comparison to the sequences used in the training of the predictor. For each Mach sequence, Jaro-Winkler distance with rest of the predictor training dataset was averaged. For the rest of the training dataset, the mean similarity was calculated by averaging over the similarity with rest of the library. The mean similarities for all Mach sequences is less than 60%.

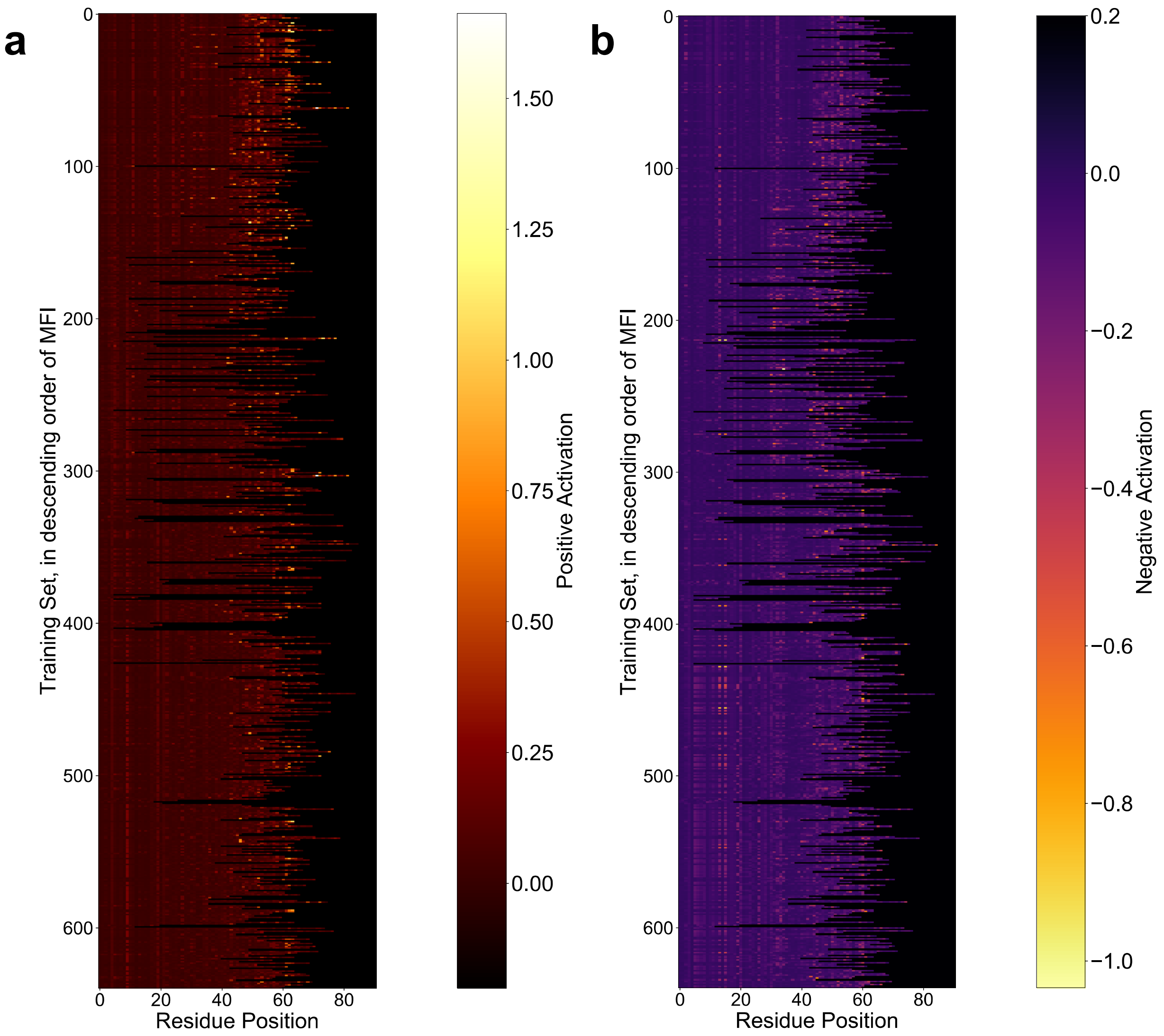

**Fig. S4. Activation map of predictor training set relative to amino acid position.** Gradient activations for sequences are arranged in descending order of experimental normalized MFI for **(a)** positive and **(b)** negative activation averaged over residue position from C-terminus. The positive activation for C-terminal residues decreases with decrease in normalized MFI values. The most active sequences have a highly positively activated C-terminus and a sparsely negatively activated C-terminus.

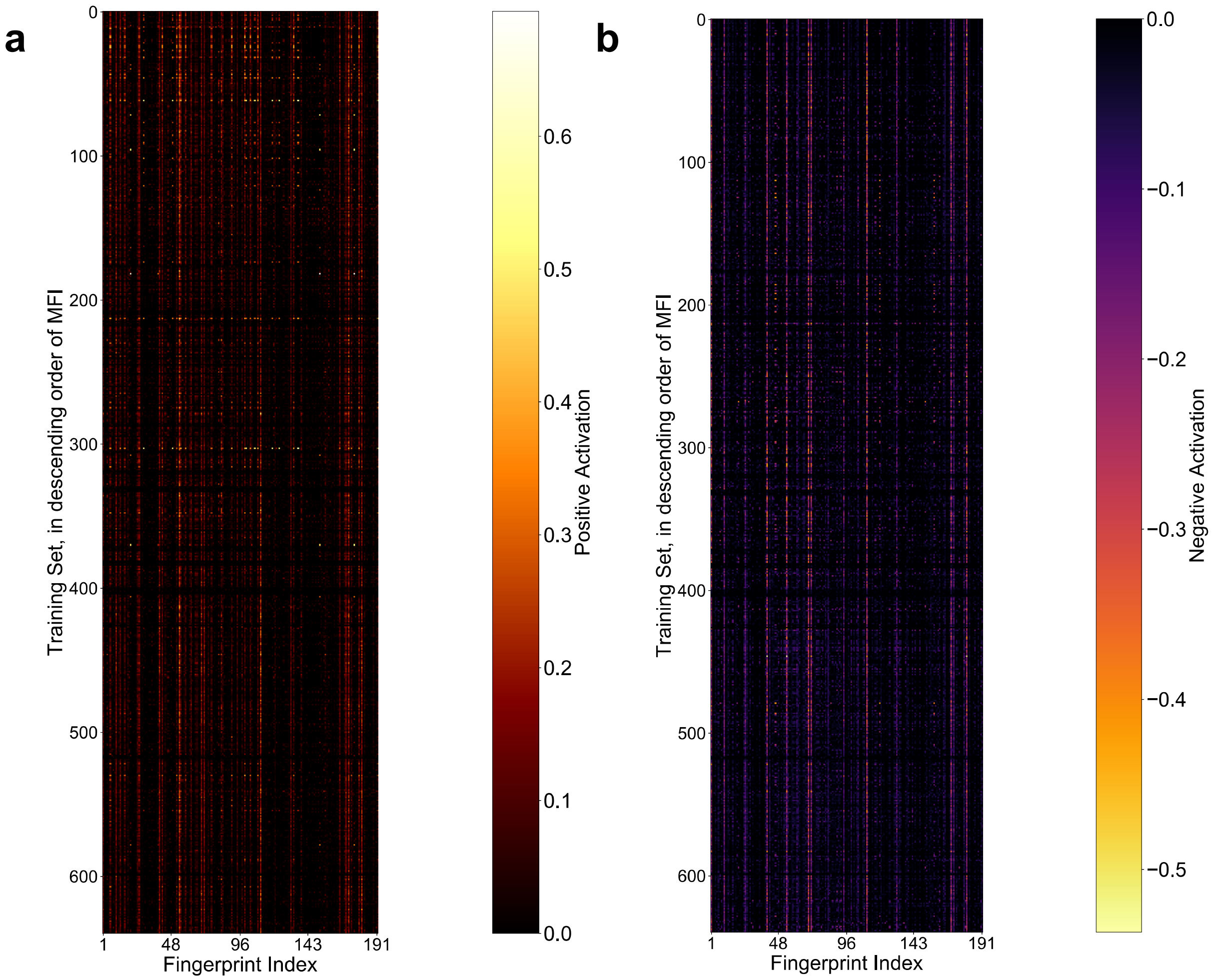

**Fig. S5. Activation map of predictor training dataset relative to fingerprint index.** Gradient activations for sequences are arranged in descending order of normalized MFI for **(a)** positive and **(b)** negative activation averaged over fingerprint index. The most positively activated substructures by residue are for aminohexanoic acid, β-alanine, aspartic acid, threonine and serine.

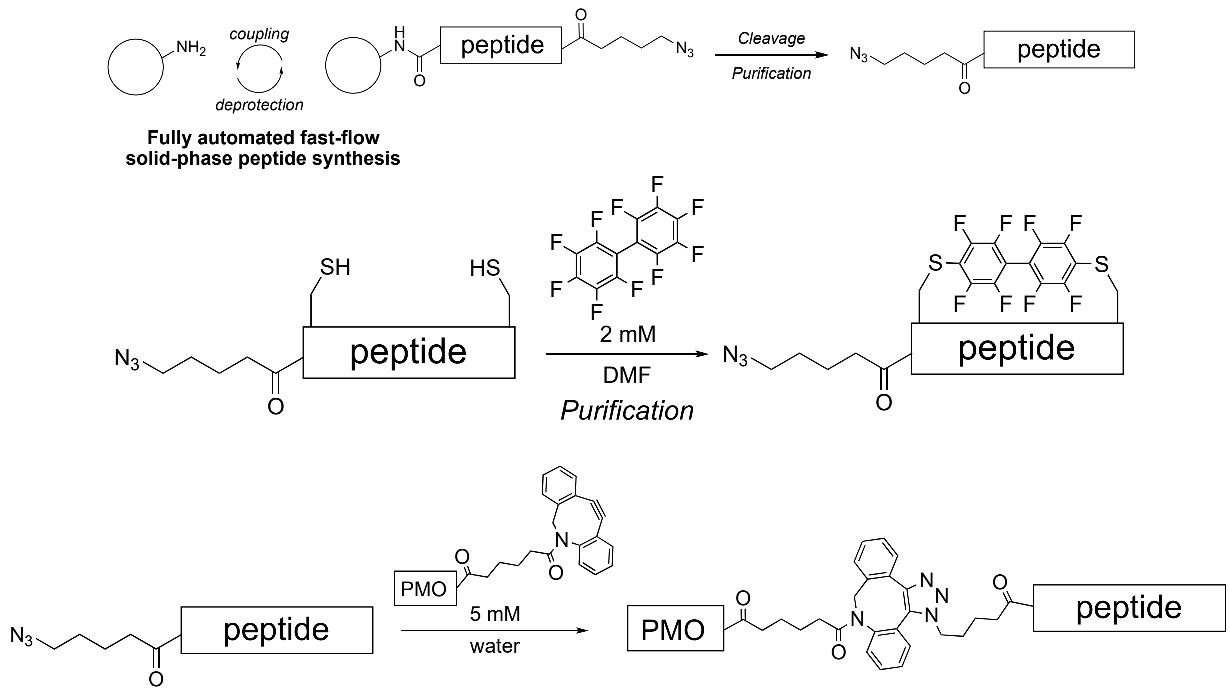

**Fig. S6. Synthesis route to Mach peptides.** Predicted sequences were synthesized by fully automated SPPS, cyclized, and conjugated to PMO. Our synthesis technology can reliably synthesize long polycationic peptides in one-shot. If the predicted peptide contains a Cys macrocycle, cleaved and purified peptides were cyclized before being attached to PMO via copper-free click chemistry.

**Table S3. List of Mach peptides**. *Peptide 10 was found to degrade in solution, so its analysis was discontinued. 'X' is 6-amino hexanoic acid, and 'B' is β-alanine. **C** residues are linked through decafluorobiphenyl.

| | | FOLD OVER PMO | % ARG | PPMO MW | NET CHARGE |
|---|---|---|---|---|---|
| **MACH 1** | ALKBRSAAKAVRWPKKKIKQASK KVAKYALXXXRKKKAASKXWLQ LHWPRW | 45 | 8 | 12,645 | 18 |
| **MACH 2** | PPLRNAKKKNLKNNLKMDPKFTK KVKQGALKLNRRKKNRGPKGPX KHWTT | 27 | 8 | 12,499 | 18 |
| **MACH 3** | QKKRKSKANKKNWPKGKLSIHAK DYKQGPKAKXRKQRXR | 39 | 10 | 11,324 | 17 |
| **MACH 4** | KKGKKQNKKKHRWPKKKVPQPK KMFKQGABXRX | 25 | 6 | 10,622 | 16 |
| **MACH 5** | AKKKIAKAKKHRGPNBGIHAPVS KIKDPLKXXX | 3 | 8 | 10,222 | 11 |
| **MACH 6** | ALKBRSAAKAVRWPKKAIKQASK KVAKYALKXXRKKKAASKXWLQ LHWPRW | 43 | 8 | 12,603 | 18 |
| **MACH 7** | XKHPXAVQBAARAWKVPAAALW KKKRLKKSSKQKKKWLWKARSA XKYXRLI | 36 | 8 | 12,645 | 18 |
| **MACH 8** | BKGKNLLAKIRRGPNGGNBQGSQ GYLLYLLXRXRRQRXXYPWWRX KHXRWXXRXRGHXRRRRQXLKP DRXRGGKGSVS | 39 | 21 | 15,929 | 22 |
| **MACH 9** | KKKKNLNBKSRRGPNGGALQPSQ GYLQPLNXRXRRQRXXYPWWRX KHXRWRXRYHXRRRRQXLKPG | 38 | 21 | 14,845 | 22 |
| **MACH 11** | TSNLKLHLAPPVKKKALKKPLYK AKKKKKVVSPTWXTDQEW | 4 | 0 | 11,423 | 11 |
| **MACH 12** | KGGKNLAKKIRRGPNGGALQPSQ GYLLYLBXRXRRQRXXGPXWRX KHXRWXXXXXRPTHXRRRRQXL **C**PGRXRP**C**RGSVS | 40 | 20 | 16,285 | 22 |
| **MACH 13** | AKKKKLGBKALRWPNGK**C**PQPK EK**C**PKYLLGRXRRKRXRYPWWR XKHRRW | 30 | 18 | 13,228 | 20 |

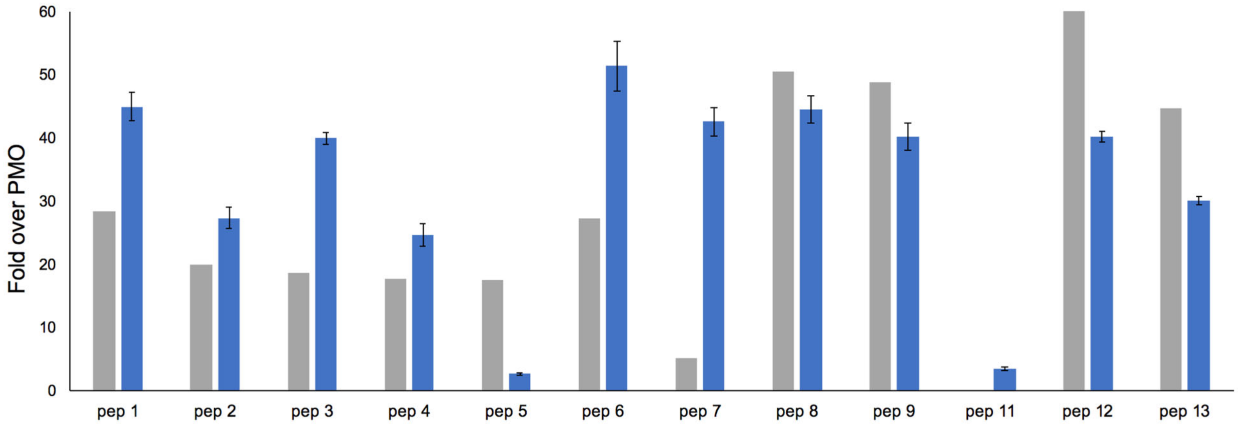

**Fig. S7. Experimental vs Predicted activity of Mach peptides.** Mach peptides enhance delivery of PMO by 40-50 fold as determined by the HeLa 654 assay. Experimental activity (blue) is comparable to predicted activity (grey). Mach12 predicted activity is off the scale, at 140. Each bar represents group mean ± SD, N = 3.

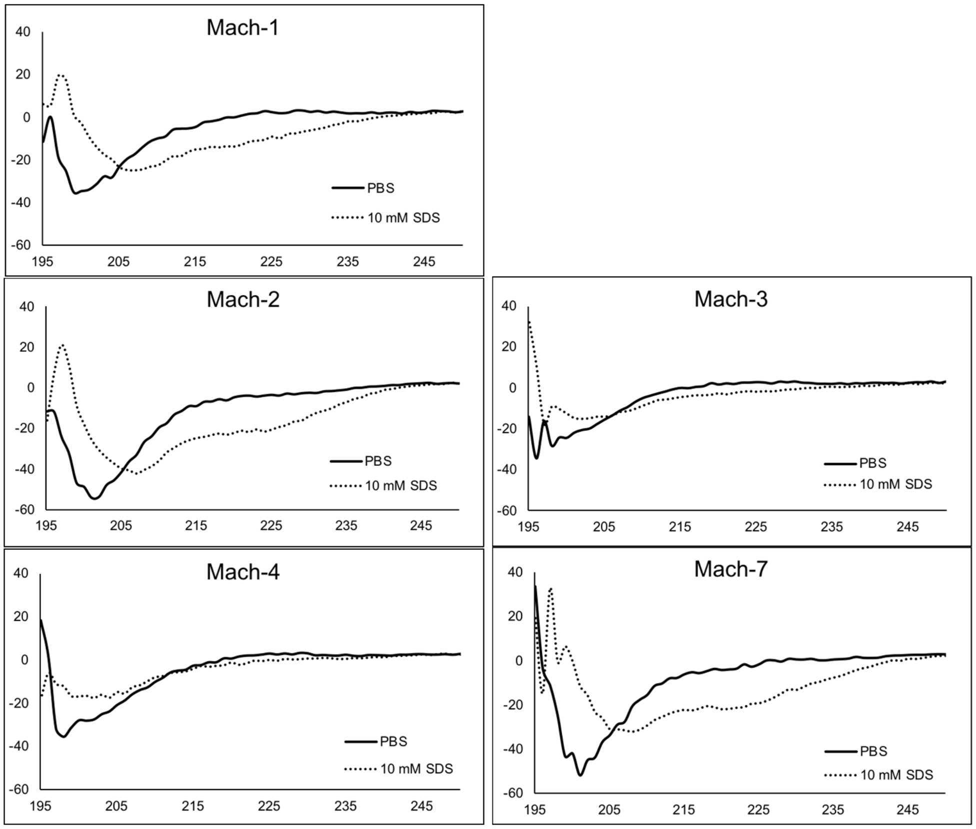

**Fig. S8. Circular dichroism of azide-Mach peptides.** 20 µM Mach peptides were either incubated in PBS or 10 mM SDS before analysis using circular dichroism. In buffer, these peptides do not exhibit secondary structure. In a lipid environment, Mach1, Mach2, and Mach7 exhibit partial alpha helicity.

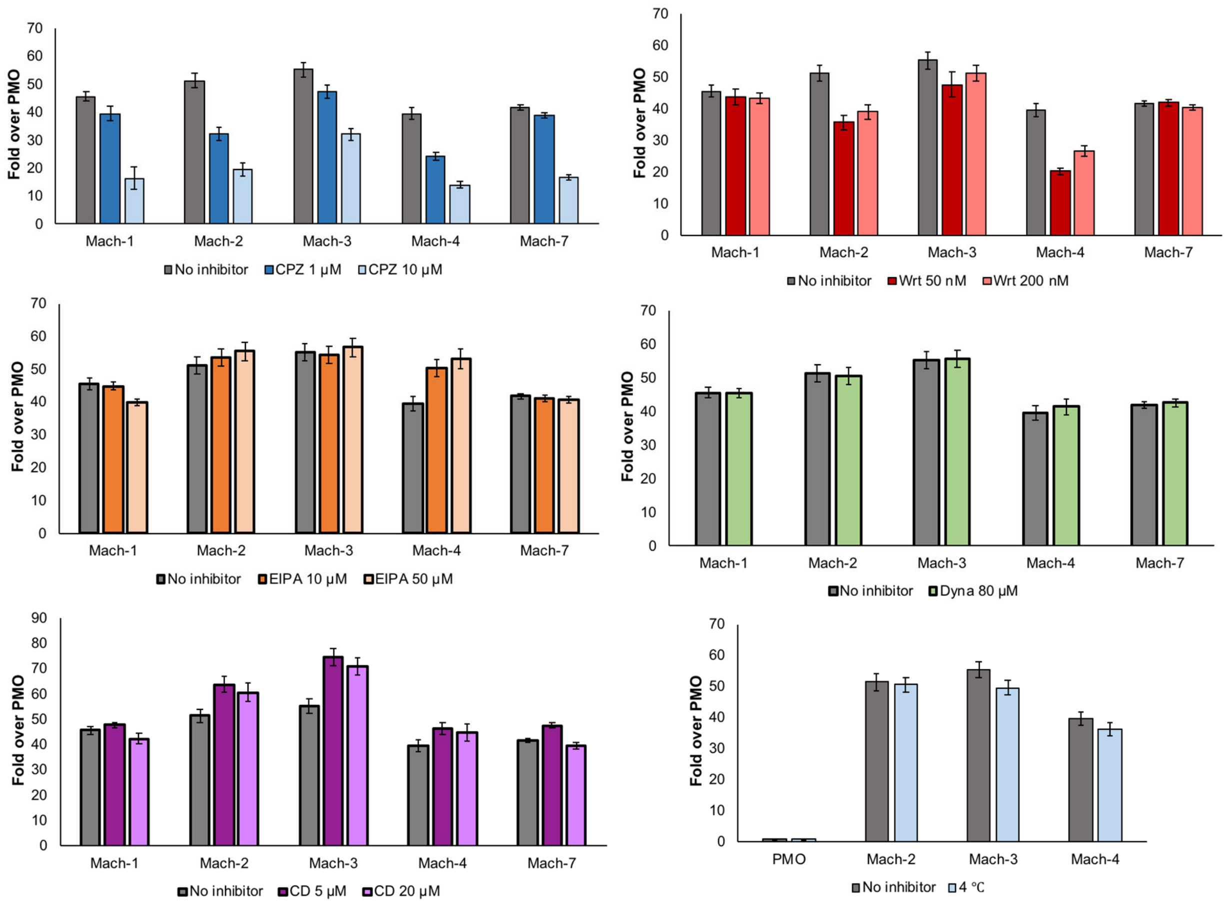

**Fig. S9. PMO-Mach peptides enter cells by energy-dependent endocytosis.** PMO activity of Mach constructs when treated with various endocytosis inhibitors. Chlorpromazine (CPZ) has a dose-dependent effect on PMO activity for each of the Mach constructs, indicating that constructs may enter via clathrin-mediated endocytosis. Each bar represents group mean ± SD, n = 3.

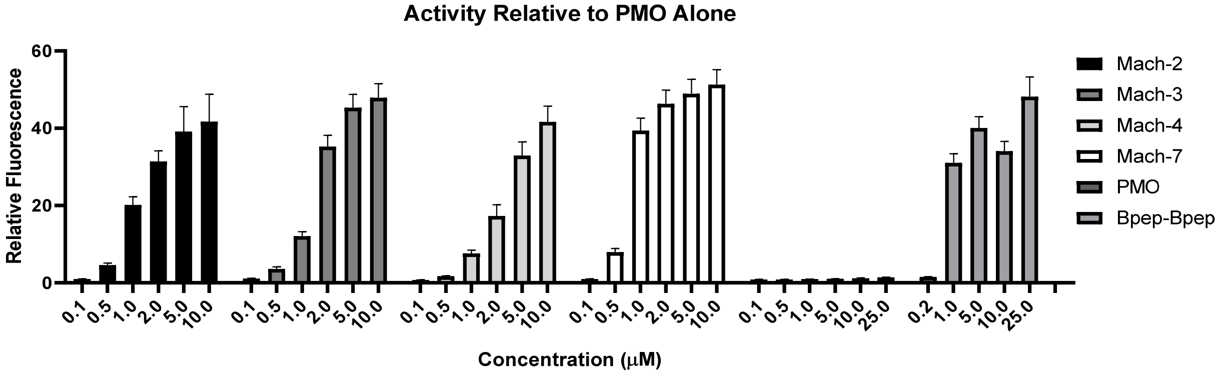

**Fig. S10. Dose-response in HeLa 654 cells (Activity).** PMO-Mach constructs elicit a dose-dependent increase in EGFP fluorescence. Included here is chimera PMO-Bpep-Bpep, a previously reported high-performing PMO-peptide. Each bar represents group mean ± SD. For Mach2 and Bpep-Bpep n = 2; PMO, Mach3, Mach4, Mach7 n = 4.

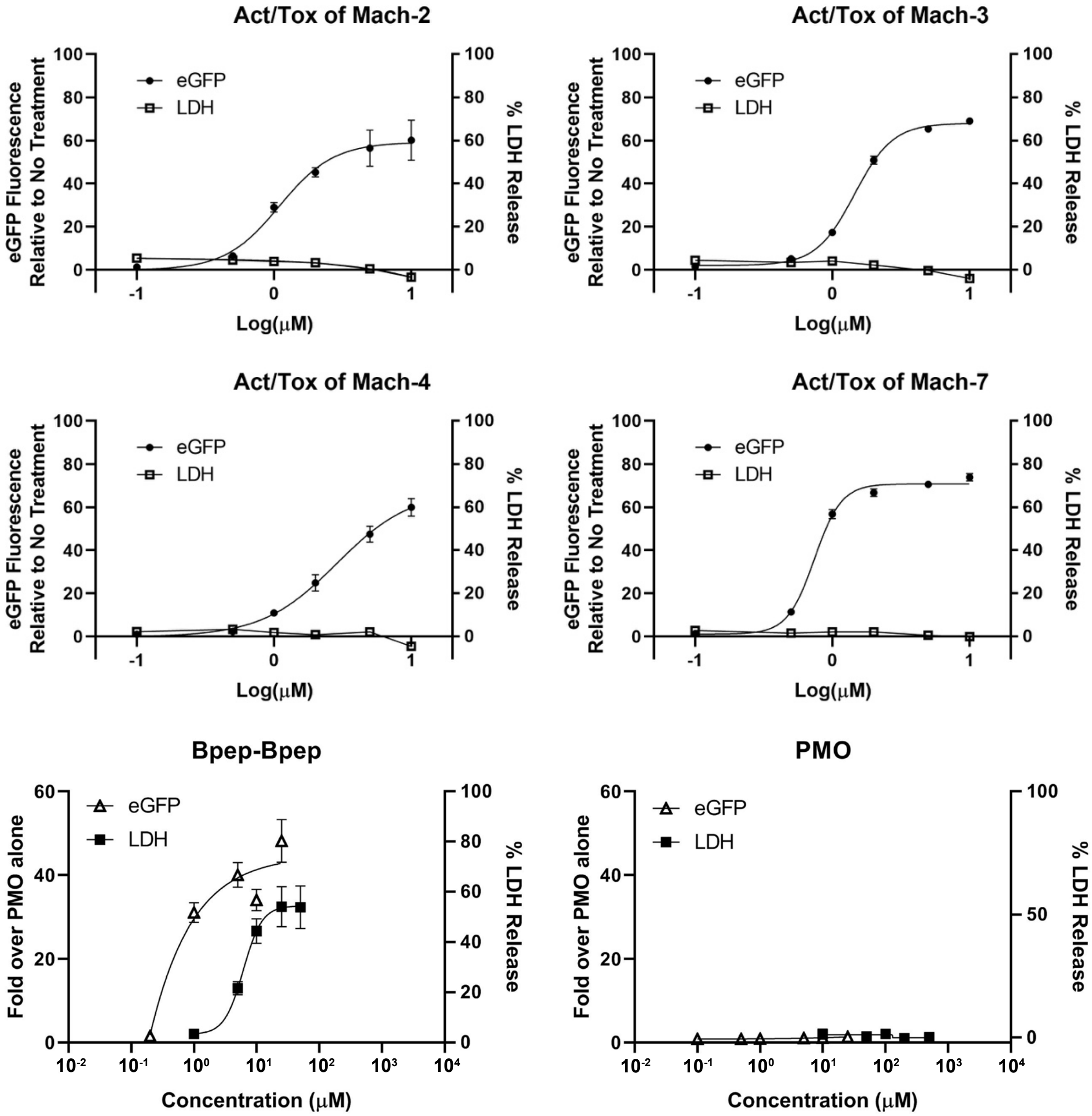

**Fig. S11. Dose response curves corresponding to activity and toxicity in HeLa 654.** HeLa 654 cells were treated with varying concentrations of PMO-Mach constructs for 22 h. Following treatment, cell supernatant was removed and tested for LDH release, reported as % LDH release relative to full lysis control. The cells were then analyzed for EGFP fluorescence by flow cytometry.

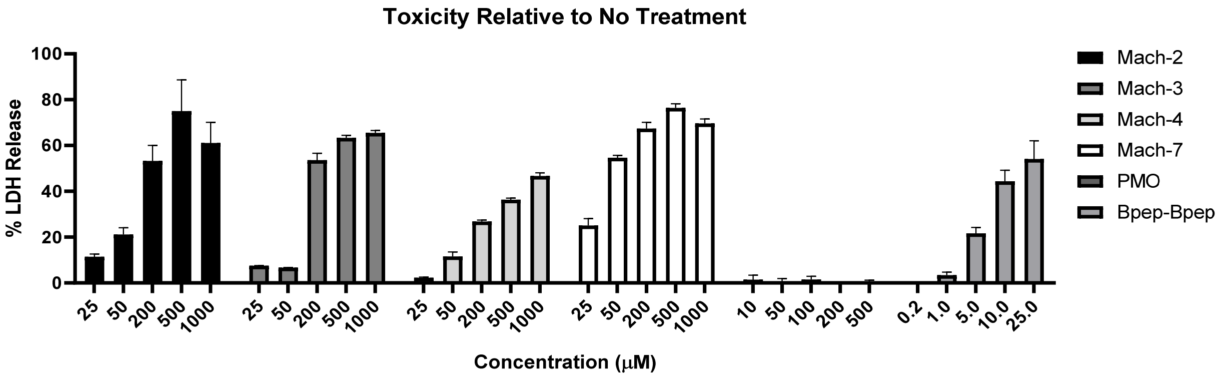

**Fig. S12. Dose-response in RPTEC (Toxicity).** PMO-Mach constructs elicit a dose-dependent increase in membrane toxicity as measured by LDH release assay. LC50 of PMO-Mach constructs are between 100-200 μM, in contrast to PMO-Bpep-Bpep, which has a significantly lower LC50 near 10 μM. Each bar represents group mean ± SD. For Mach2 n = 3; Mach3 and Mach7 n = 5; Mach4, PMO, Bpep-Bpep n = 6.

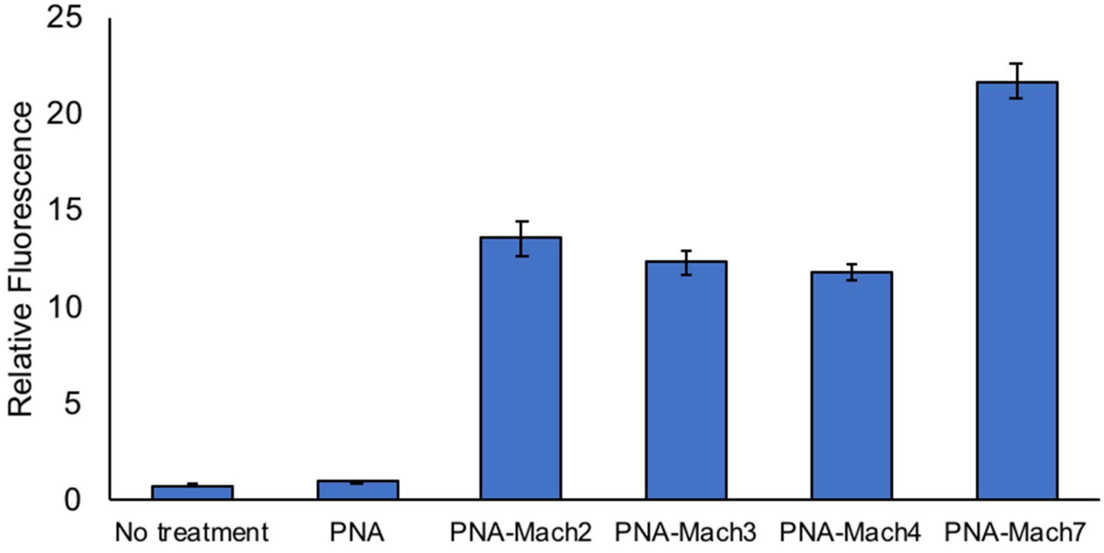

**Fig. S13. Mach peptides enhance delivery of peptide nucleic acid (PNA).** PNA-Mach constructs were evaluated at 5 μM in the HeLa EGFP 654 assay. Each bar represents group mean ± SD, n = 3.

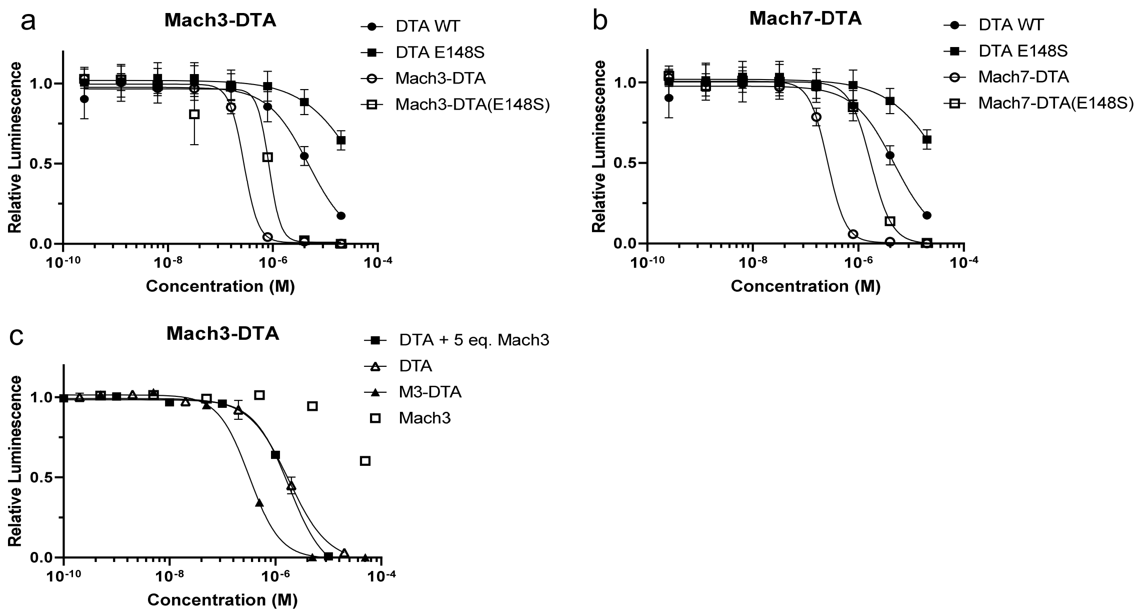

**Fig. S14. Mach-DTA conjugates produce dose-dependent toxicity in HeLa cells**. Attachment of WT DTA to (a) Mach3 or (b) Mach7 produces significantly greater activity than attachment to DTA (E148S) which has 300-fold lower activity than wild-type. (c) Covalent attachment of Mach3 is required for DTA constructs to be delivered to the cytosol. DTA alone has the same toxicity as DTA co-incubated with 5 equivalents of Mach3 peptide. Each point represents group mean ± SD, n = 3.

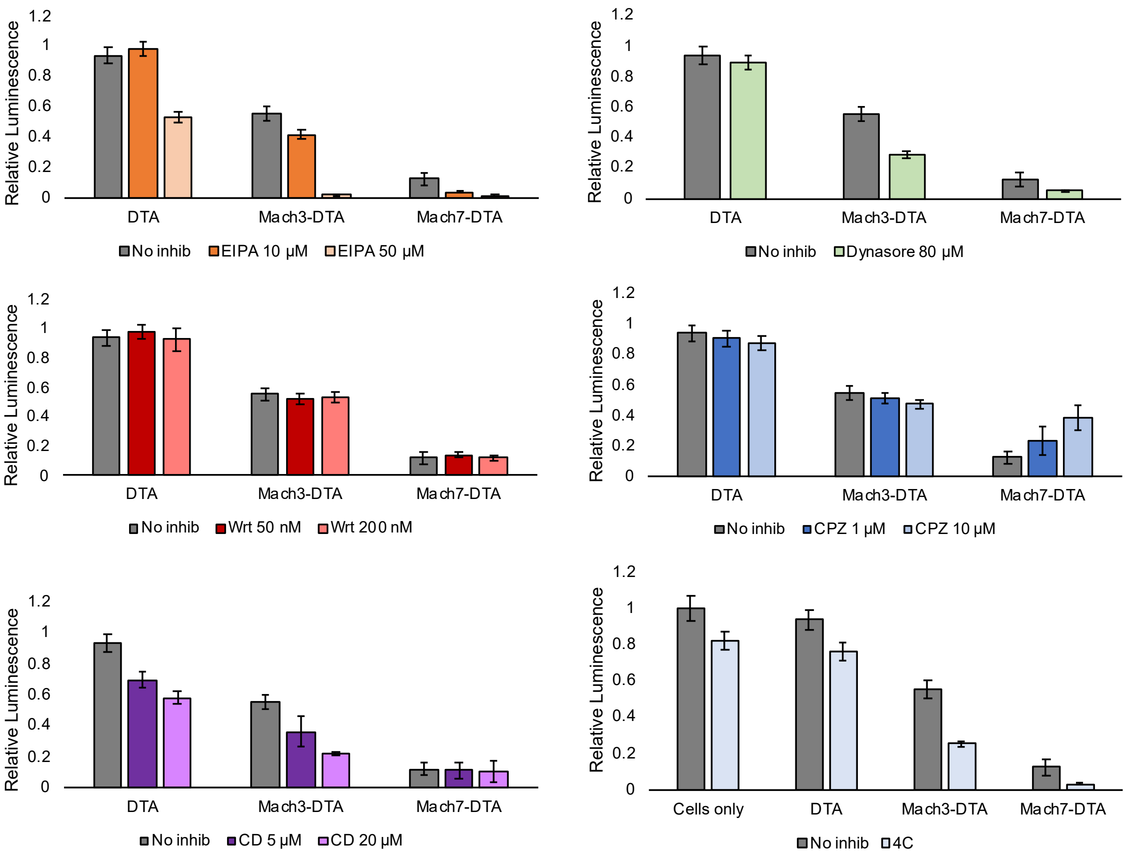

**Fig. S15. Endocytosis inhibition of Mach-DTA constructs** was analyzed at 1 µM**.** An increase in relative luminescence would indicate increased cell viability and therefore reduced cytosolic delivery of the construct. Mach7-DTA co-incubated with chlorpromazine (CPZ) displays rescued cell viability compared to its no treatment control, indicating similar mechanism of delivery as PMO-Mach constructs. Each bar represents group mean ± SD, n = 3.

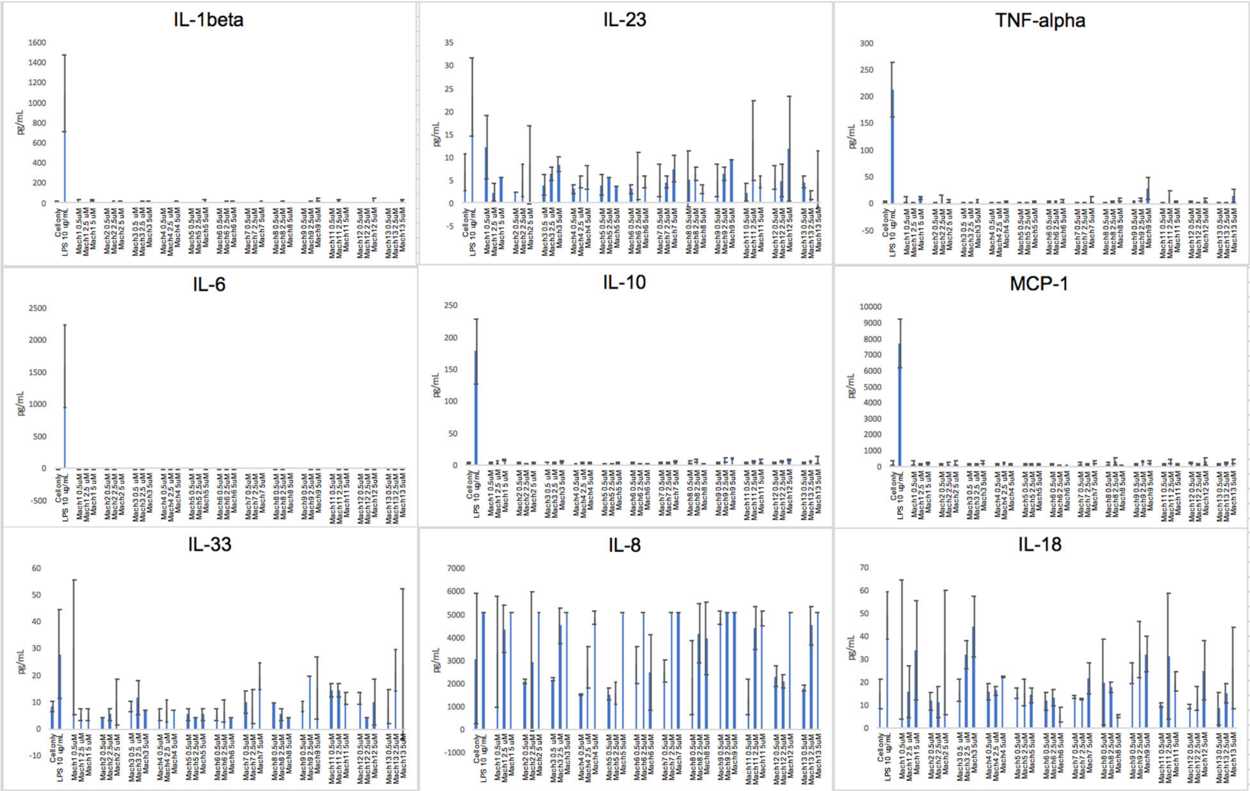

**Fig. S16. PMO-Mach constructs are nonimmunogenic in vitro**. Inflammation panel results of cytokines that were detected in human monocyte-derived macrophages. IL-1B, TNF-a, IL-6, IL-10, and MCP-1 are all released after treatment with lipopolysaccharide (LPS), but exhibit no significant increase after treatment with PMO-Mach constructs. Each bar represents group mean ± SD, n = 2.

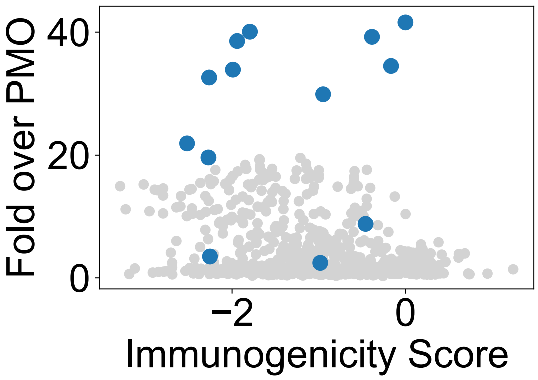

**Fig. S17. In silico immunogenicity score for Mach and training sequences.** Predicted immunogenicity for Mach sequences is within the range of the predicted immunogenicity for the sequences used in training of predictor. Mach sequences have a substantially higher experimental normalized MFI within the same range of immunogenicity, in comparison to the sequences used in the training of the predictor. The immunogenicity scores are the likelihood of being a T-cell epitope. The values are calculated using an online predictor.(*51*)

#### Appendix II: LC-MS Characterization

<u>PMO-DBCO (Method A)</u>
Mass Expected: 6527.9 Da
Mass Observed: 6527.9 Da
PMO sequence: GCT ATT ACC TTA ACC CAG

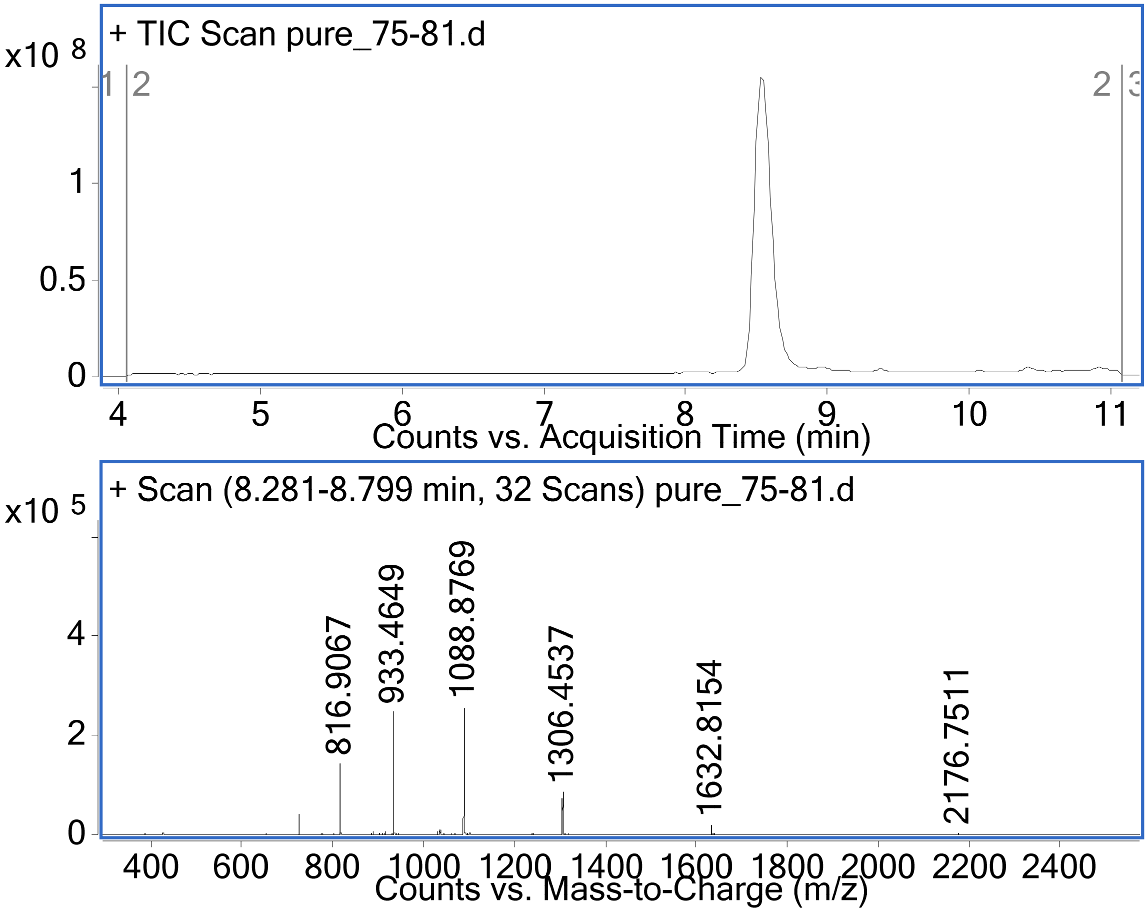

PMO-Mach1 (Method B)
Mass Expected: 12645.4 Da
Mass Observed: 12645.6 Da
Peptide sequence:
ALKBRSAAKAVRWPKKKIKQASKKVAKYALXXXRKKKAASKXWLQLHWPRW

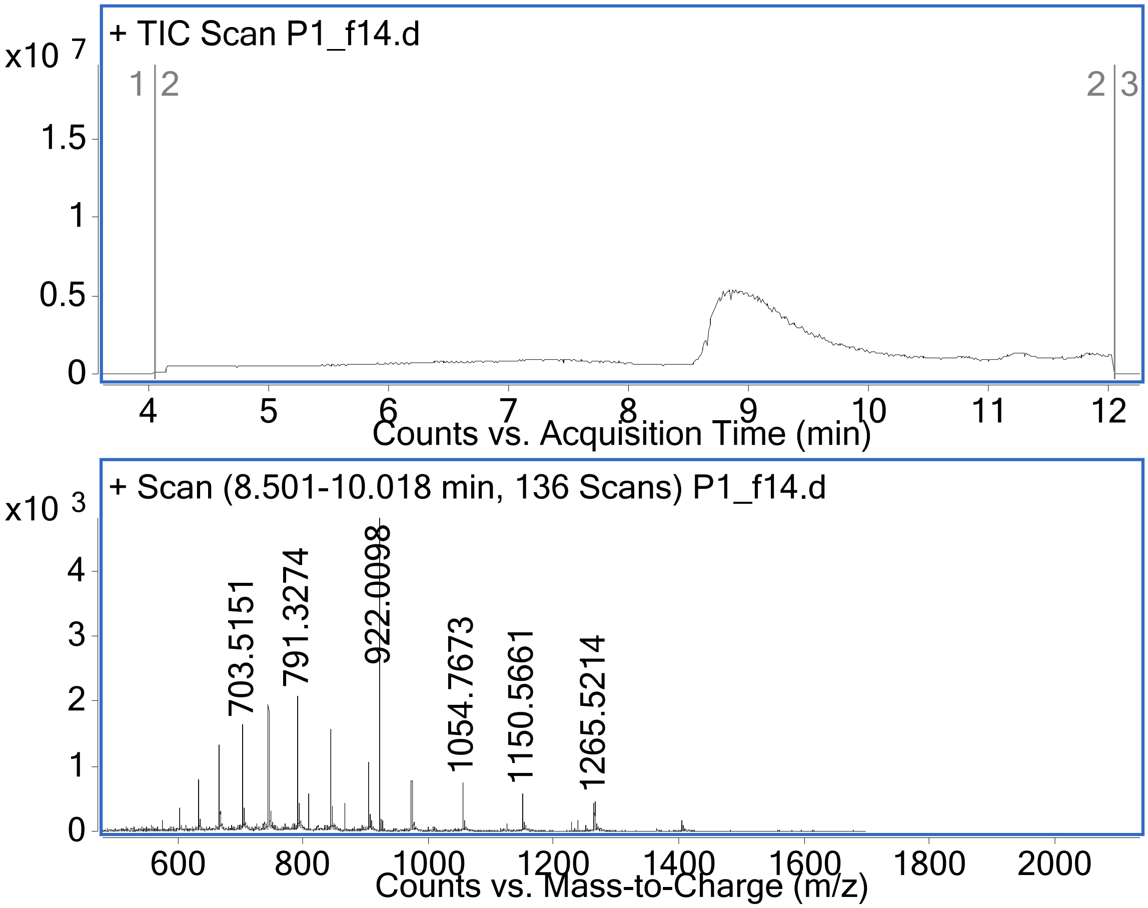

PMO-Mach2 (Method B)
Mass Expected: 12499.1 Da
Mass Observed: 12499.2 Da
Peptide sequence:
PPLRNAKKKNLKNNLKMDPKFTKKVKQGALKLNRRKKNRGPKGPXKHWTT

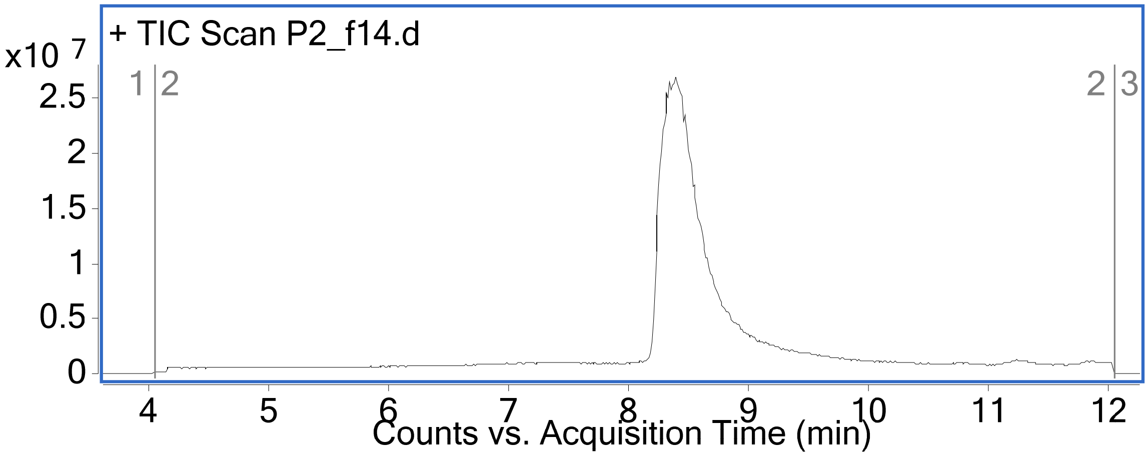

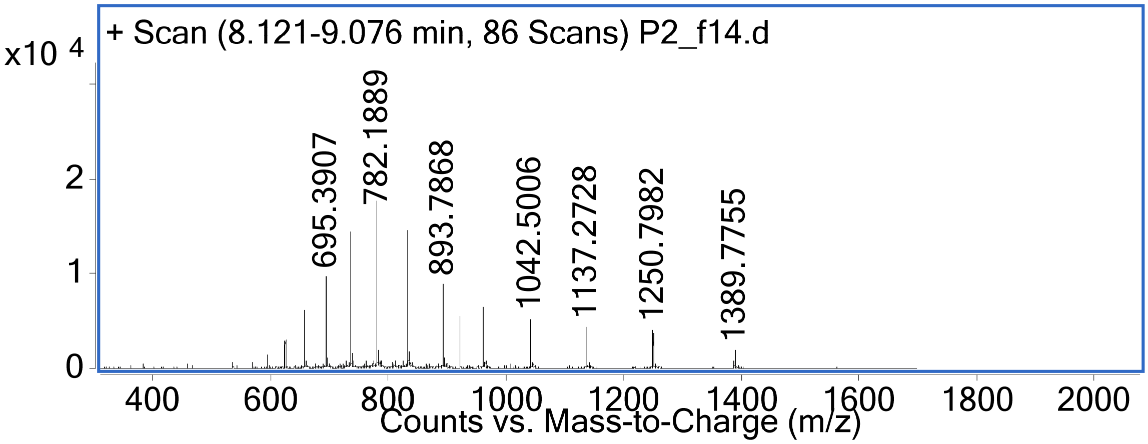

<u>PMO-Mach3 (Method A)</u>
Mass Expected: 11323.6 Da
Mass Observed: 11324.3 Da
Peptide sequence: QKKRKSKANKKNWPKGKLSIHAKDYKQGPKAKXRKQRXR

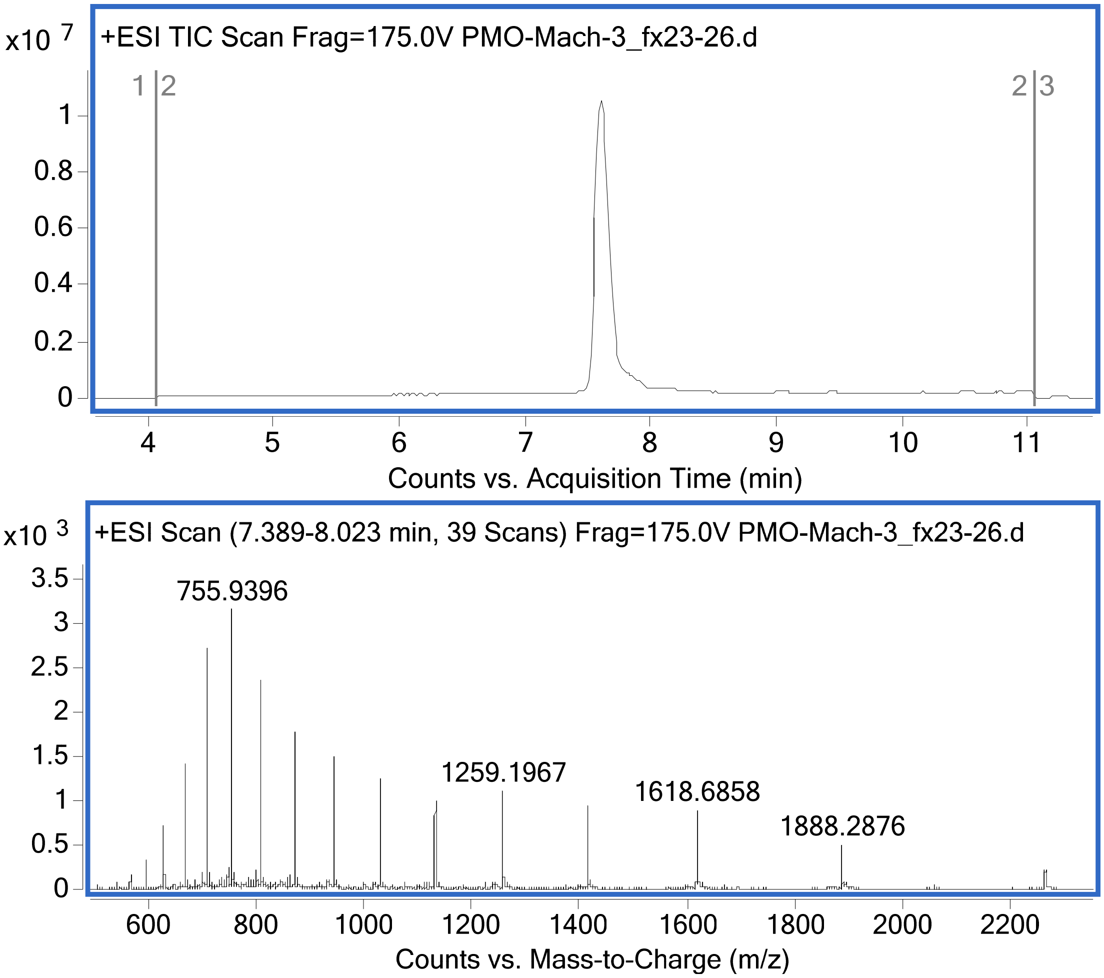

<u>PMO-Mach4 (Method A)</u>
Mass Expected: 10622.0 Da
Mass Observed: 10622.5 Da
Peptide sequence: KKGKKQNKKKHRWPKKKVPQPKKMFKQGABXRX

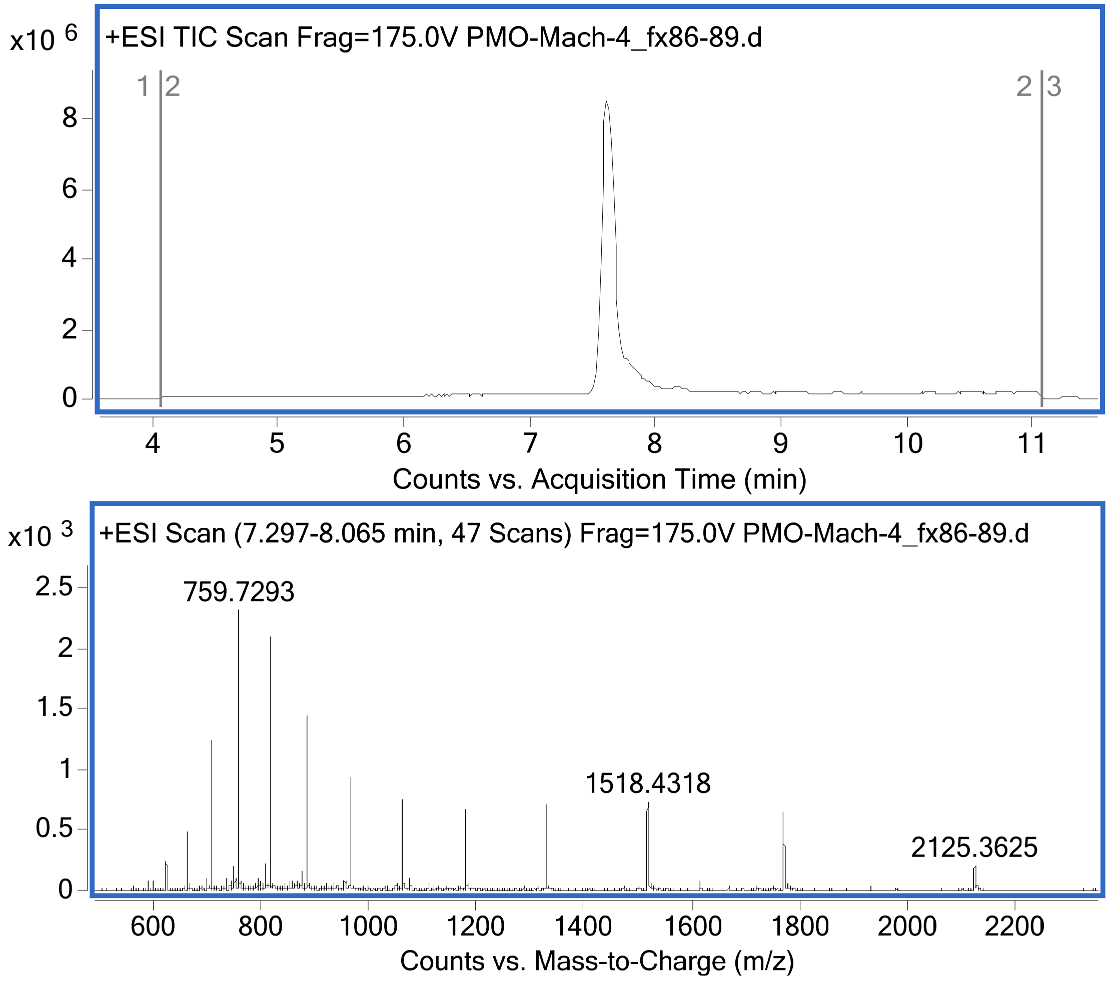

PMO-Mach5 (Method A)
Mass Expected: 10222.5 Da
Mass Observed: 10222.5 Da
Peptide sequence: AKKKIAKAKKHRGPNBGIHAPVSKIKDPLKXXX

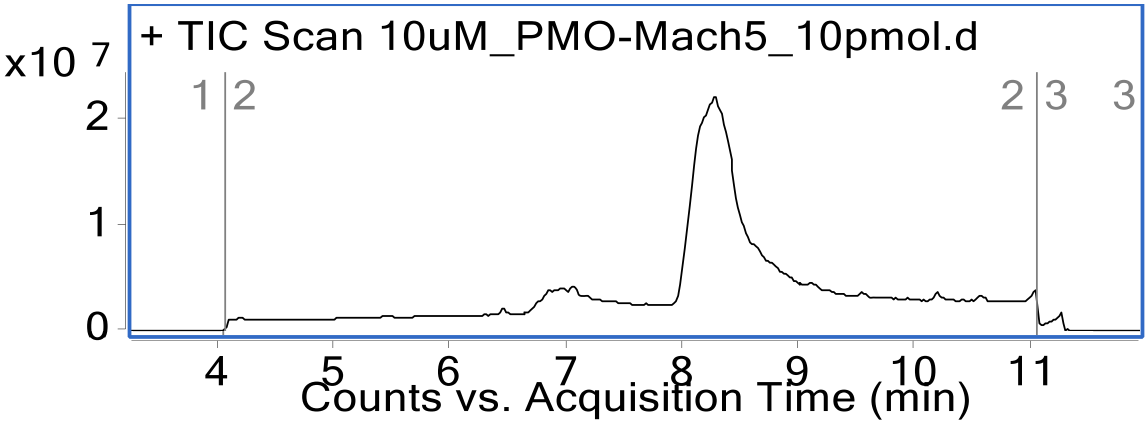

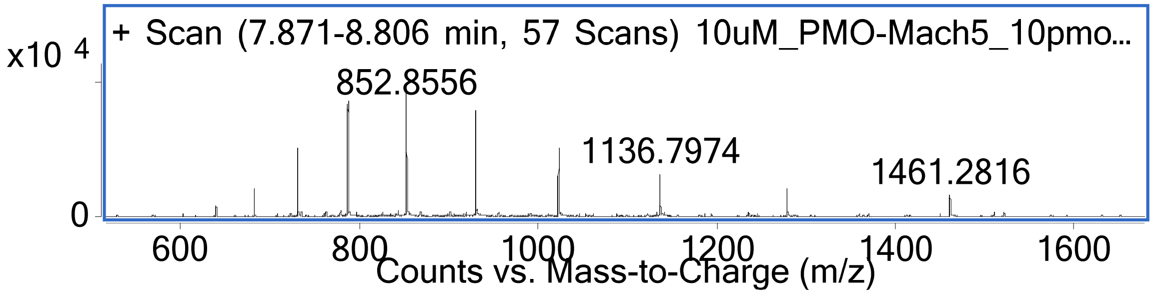

PMO-Mach6 (Method B)
Mass Expected:12603.4 g/mol
Mass Observed: 12603.4 g/mol
Peptide sequence:
ALKBRSAAKAVRWPKKAIKQASKKVAKYALKXXRKKKAASKXWLQLHWPRW

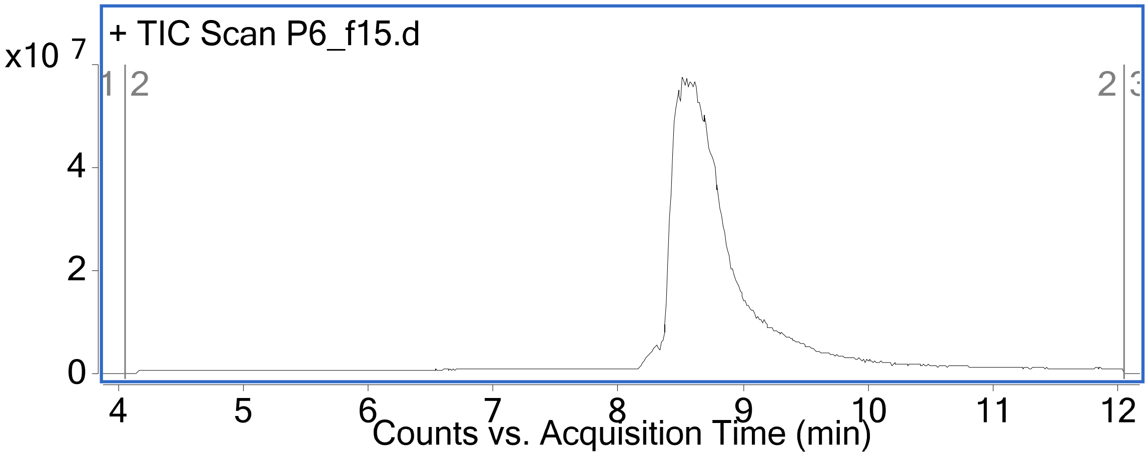

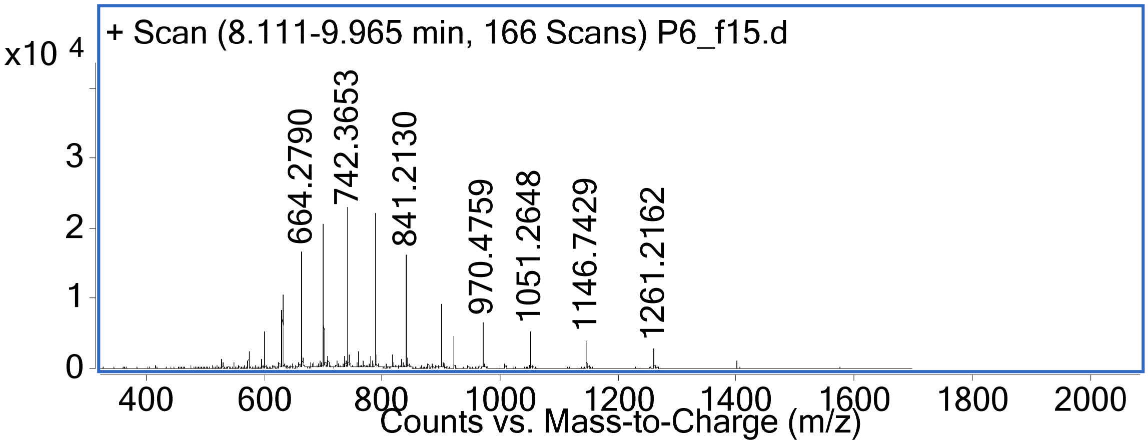

PMO-Mach7 (Method A)
Mass Expected: 12645.4 Da
Mass Observed: 12645.9 Da
Peptide sequence:
XKHPXAVQBAARAWKVPAAALWKKKRLKKSSKQKKKWLWKARSAXKYXRLI

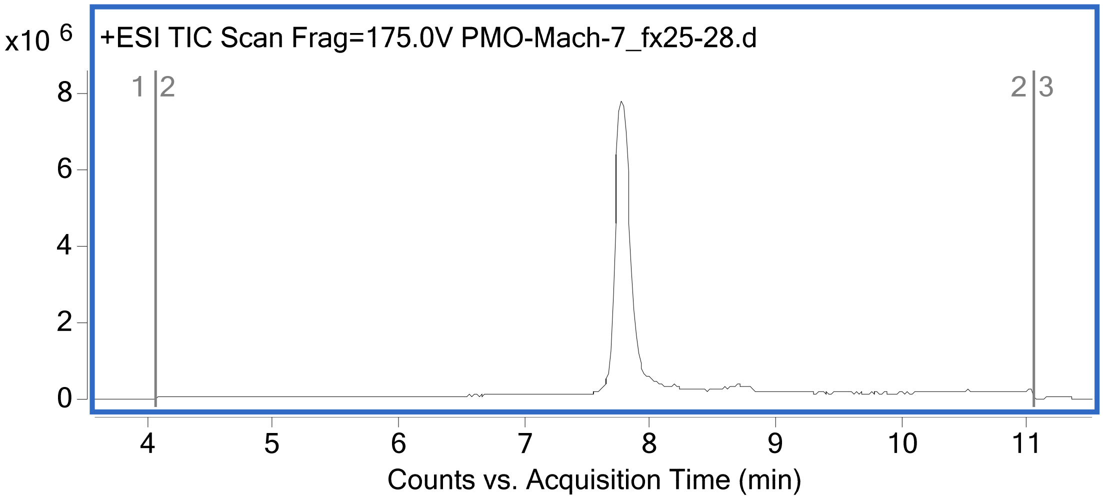

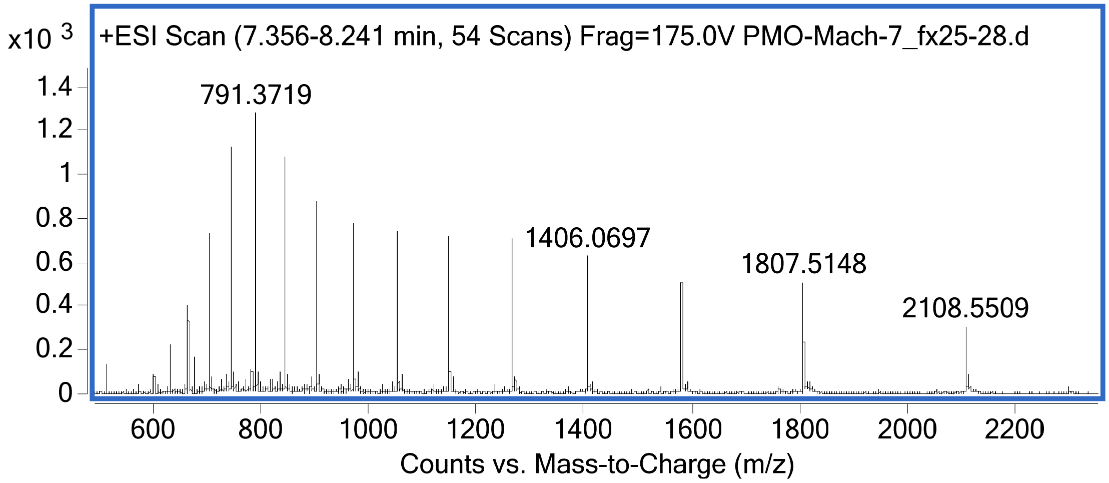

PMO-Mach8 (Method B)
Mass Expected: 15929.1 Da
Mass Observed: 15929.3 Da
Peptide sequence:
BKGKNLLAKIRRGPNGGNBQGSQGYLLYLLXRXRRQRXXYPWWRXKHXRWXXRXRG
HXRRRRQXLKPDRXRGGKGSVS

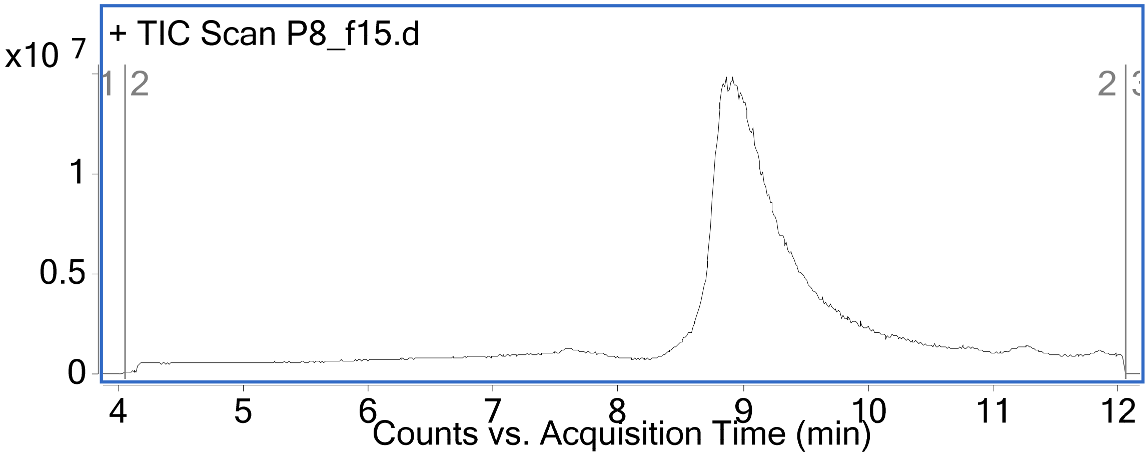

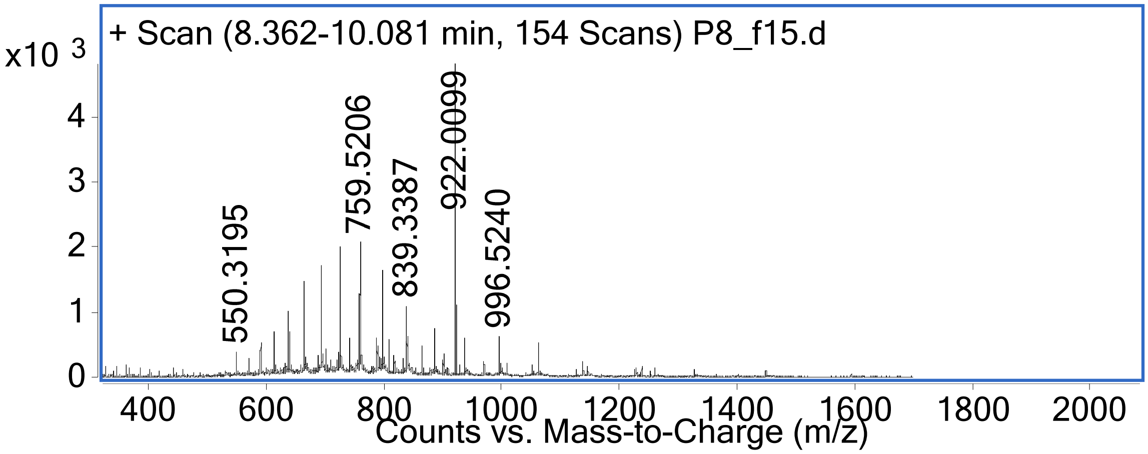

PMO-Mach9 (Method B)
Mass Expected: 14844.8 Da
Mass Observed: 14845.0 Da
Peptide sequence:
KKKKNLNBKSRRGPNGGALQPSQGYLQPLNXRXRRQRXXYPWWRXKHXRWRXRYH
XRRRRQXLKPG

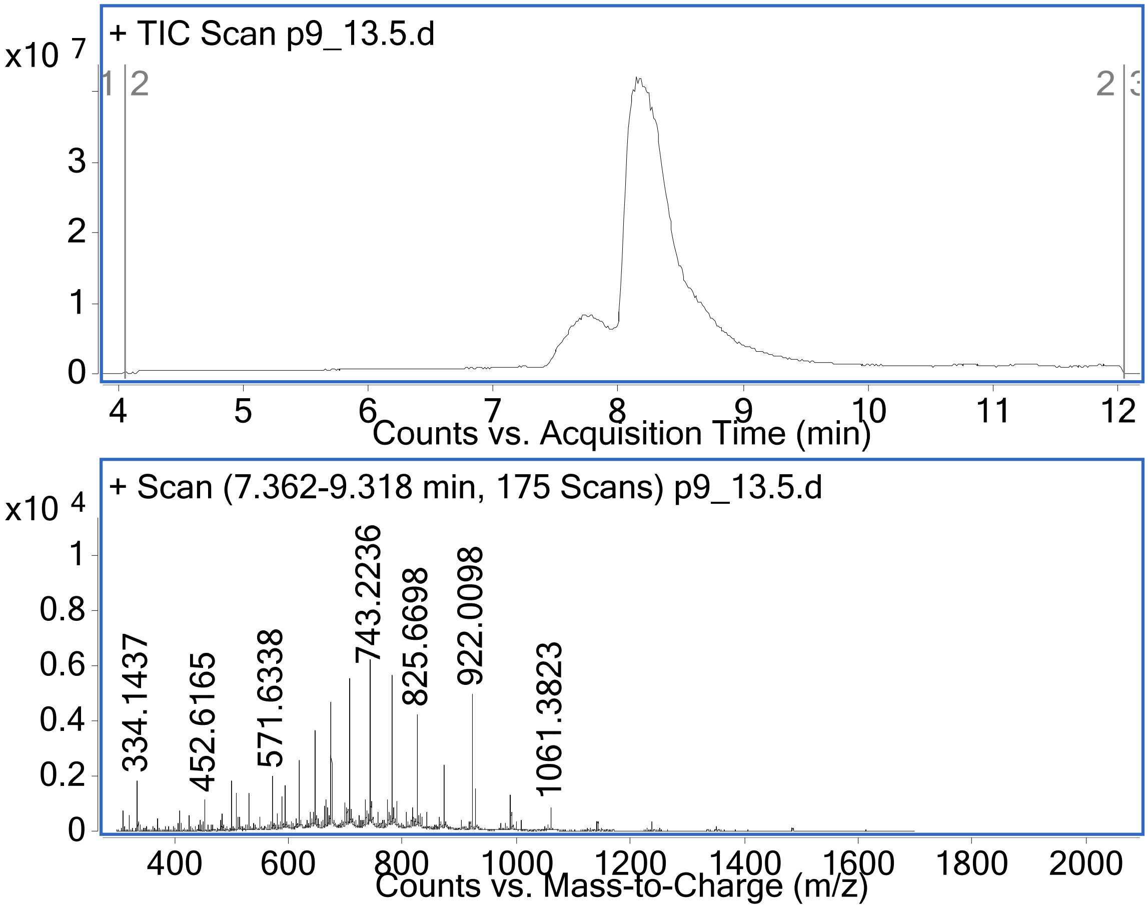

PMO-Mach11 (Method B)
Mass Expected: 11422.8 Da
Mass Observed: 11422.8 Da
Peptide sequence: TSNLKLHLAPPVKKKALKKPLYKAKKKKKVVSPTWXTDQEW

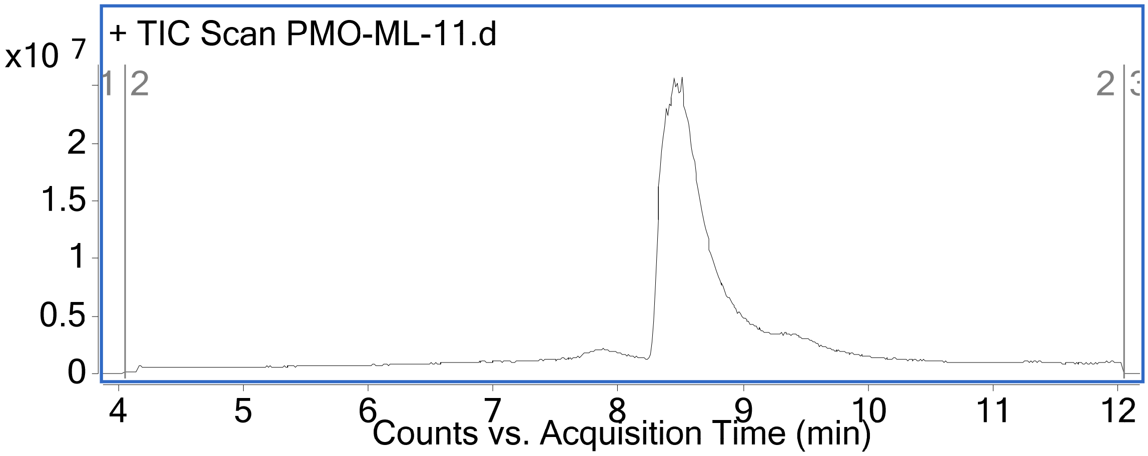

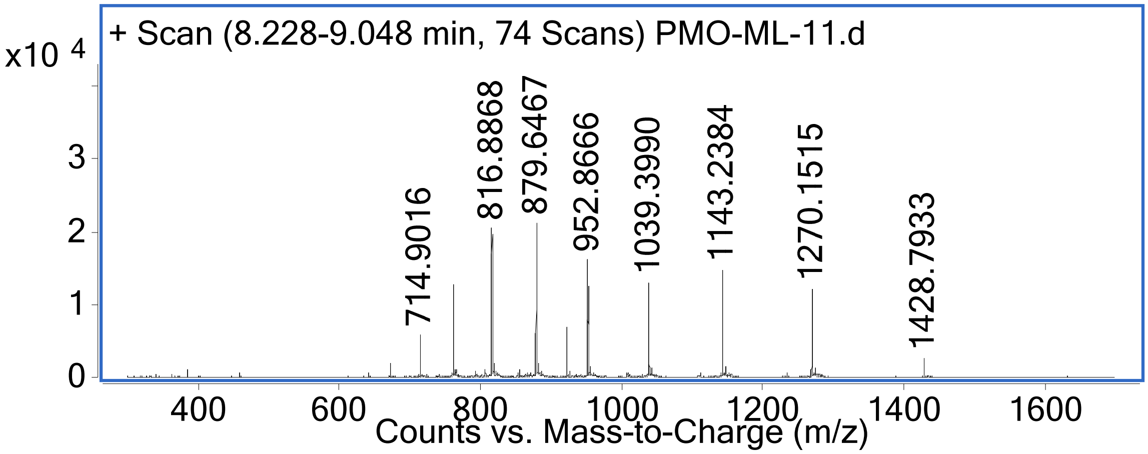

PMO-Mach12 (Method B)
Mass Expected: 16284.5 Da
Mass Observed: 16284.7 Da
Peptide sequence:
KGGKNLAKKIRRGPNGGALQPSQGYLLYLBXRXRRQRXXGPXWRXKHXRWXXXXXXR
PTHXRRRRQXL**C**PGRXRP**C**RGSVS

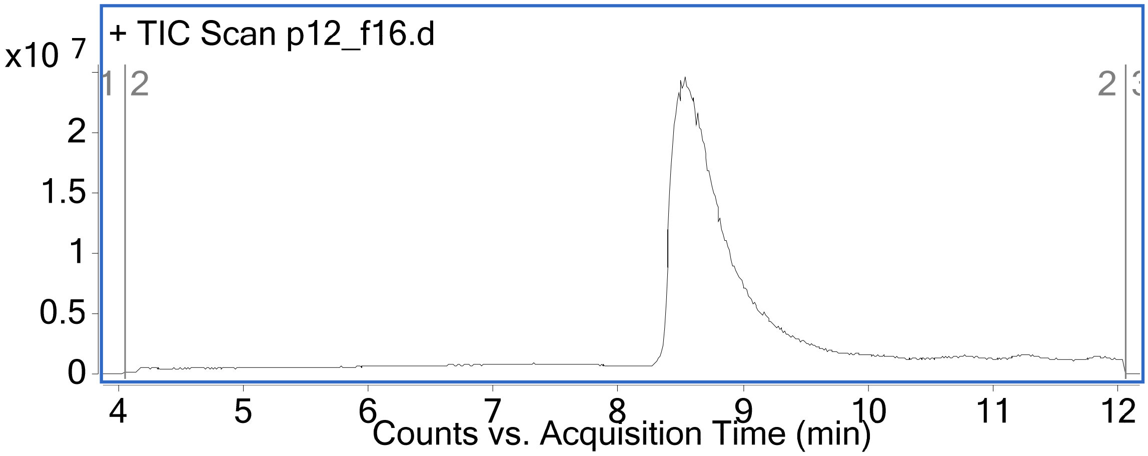

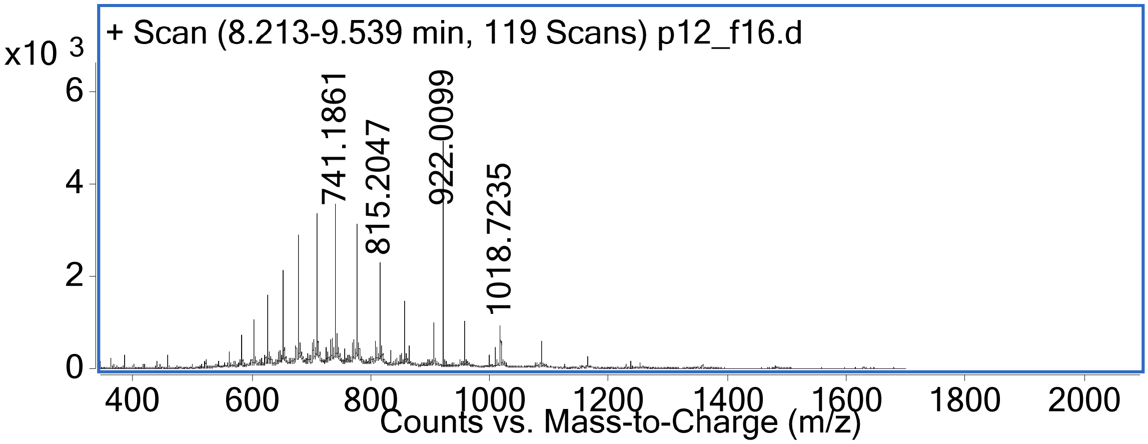

<u>PMO-Mach13 (Method B)</u>
Mass Expected: 13227.8 Da
Mass Observed: 13228.0 Da
Peptide sequence:
AKKKKLGBKALRWPNGK**<u>C</u>**PQPKEK**<u>C</u>**PKYLLGRXRRKRXRYPWWRXKHRRW

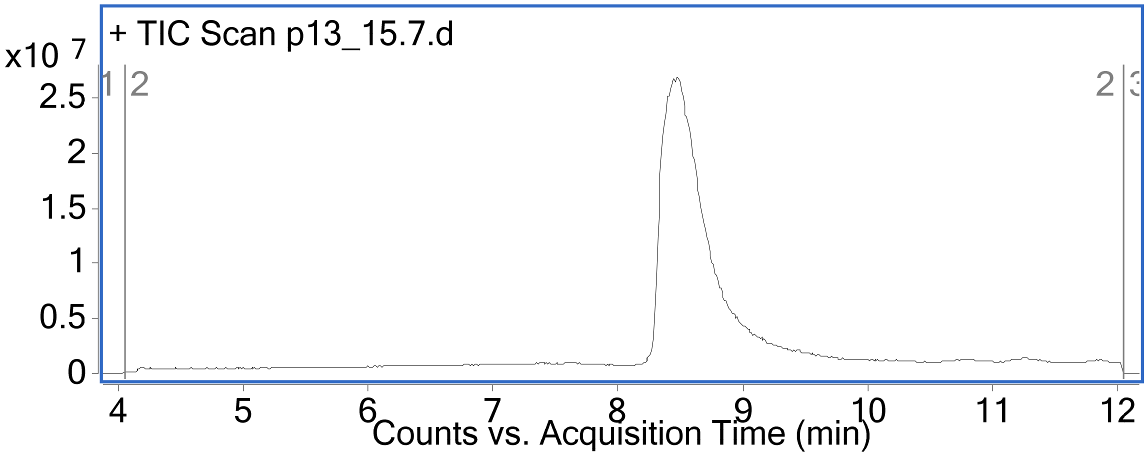

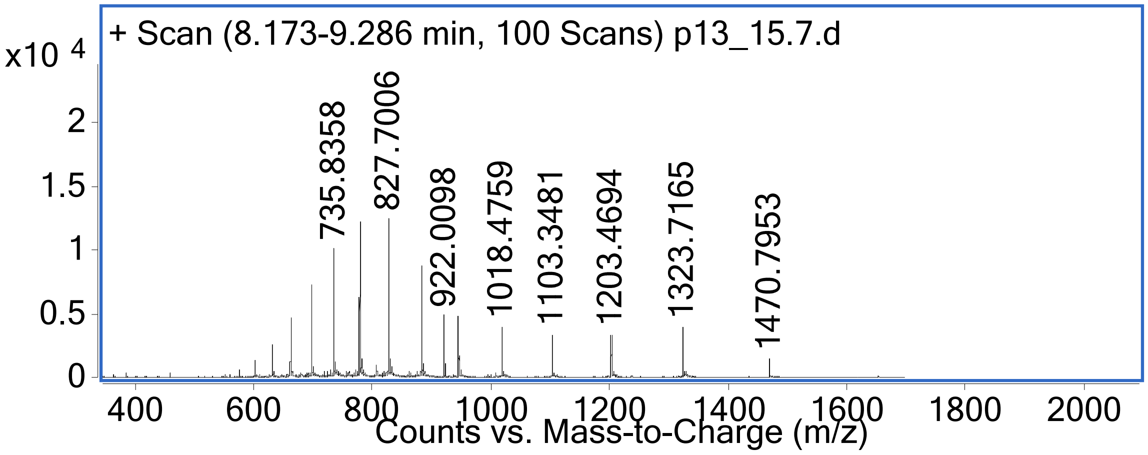

PNA-Mach2 (Method B)
Mass Expected: 11375.5 Da
Mass Observed: 11374.9 Da

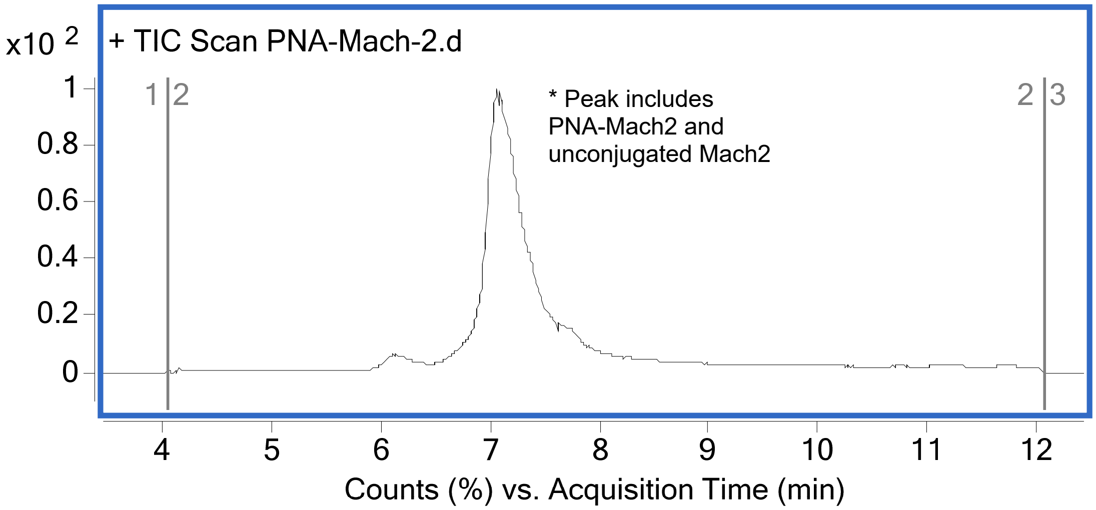

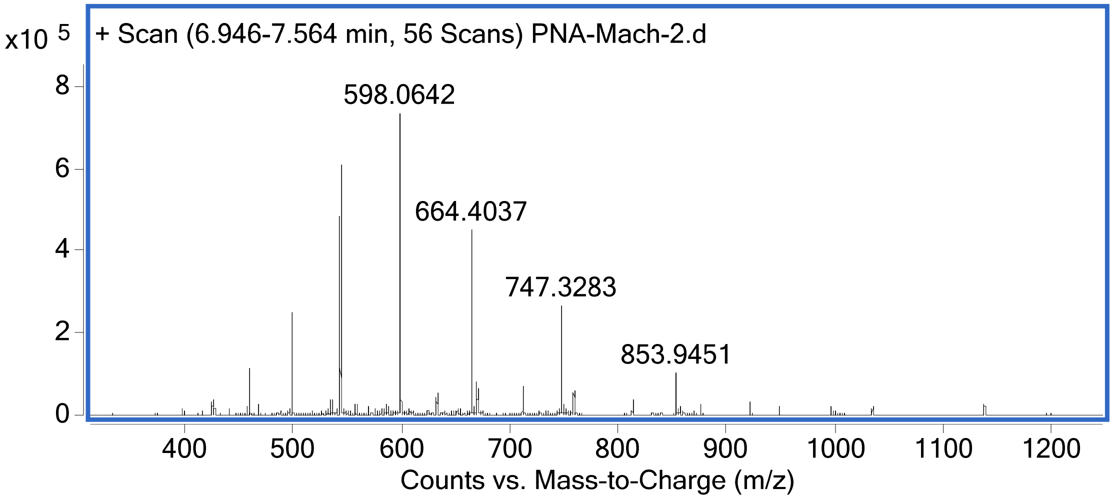

PNA-Mach3 (Method A)
Mass Expected:10200.0 Da
Mass Observed: 10200.6 Da

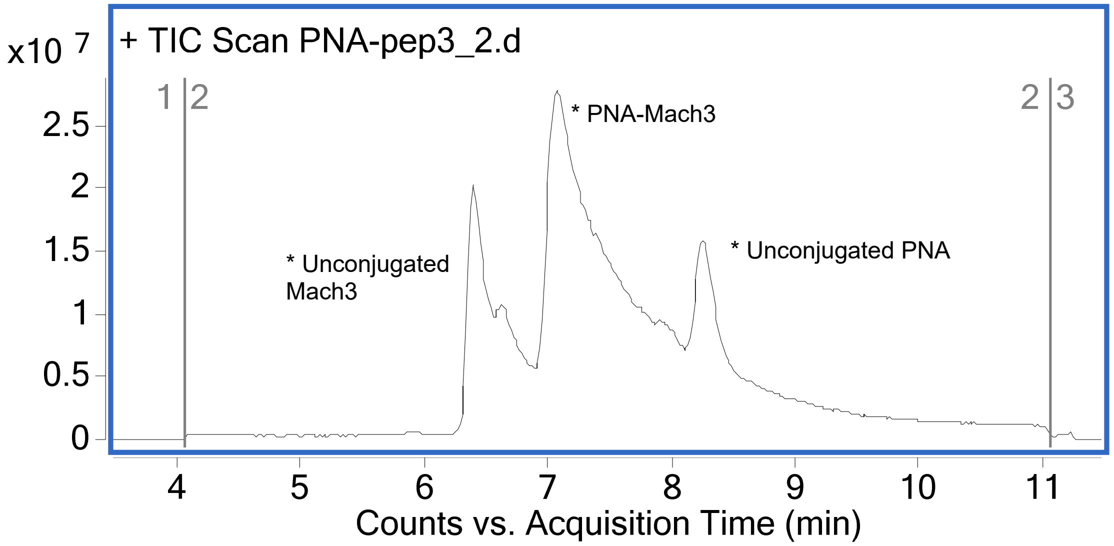

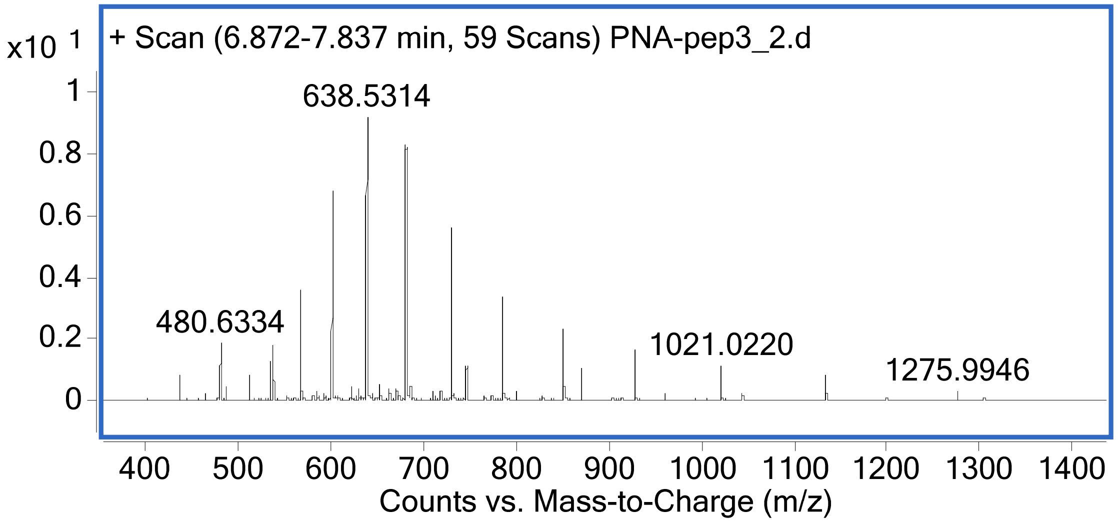

PNA-Mach4 (Method B)
Mass Expected: 9498.4 Da
Mass Observed: 9497.8 Da

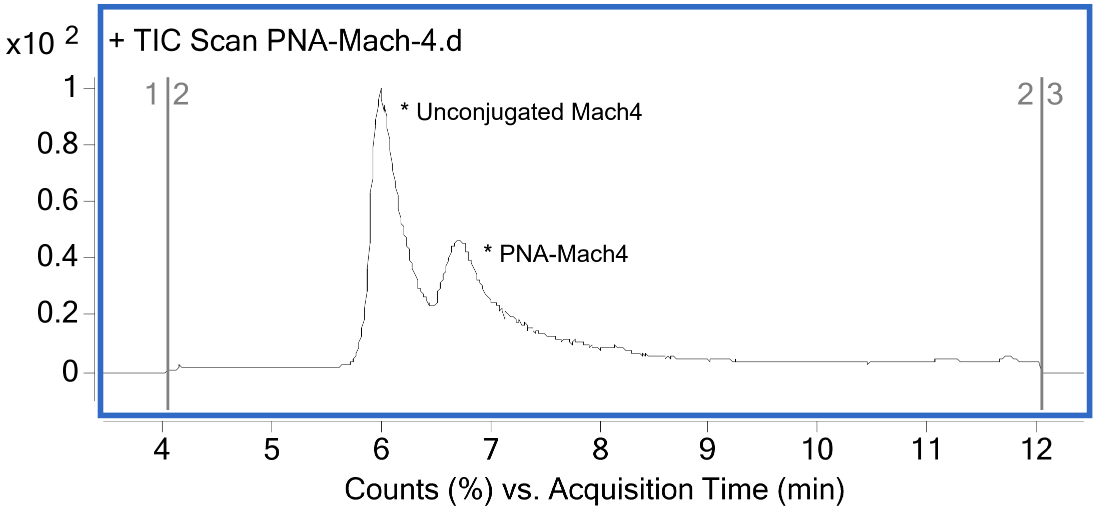

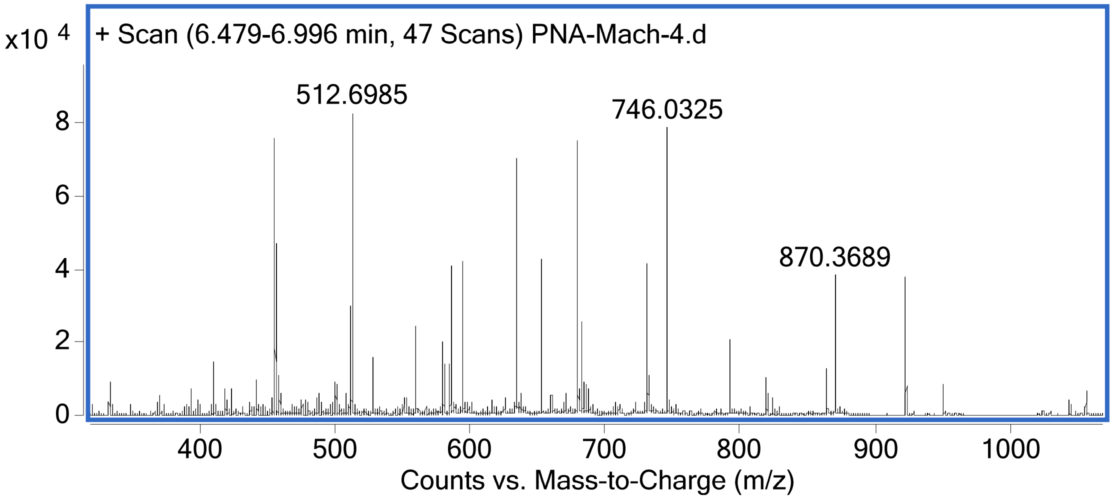

PNA-Mach7 (Method B)
Mass Expected: 11521.8 Da
Mass Observed: 11521.3 Da

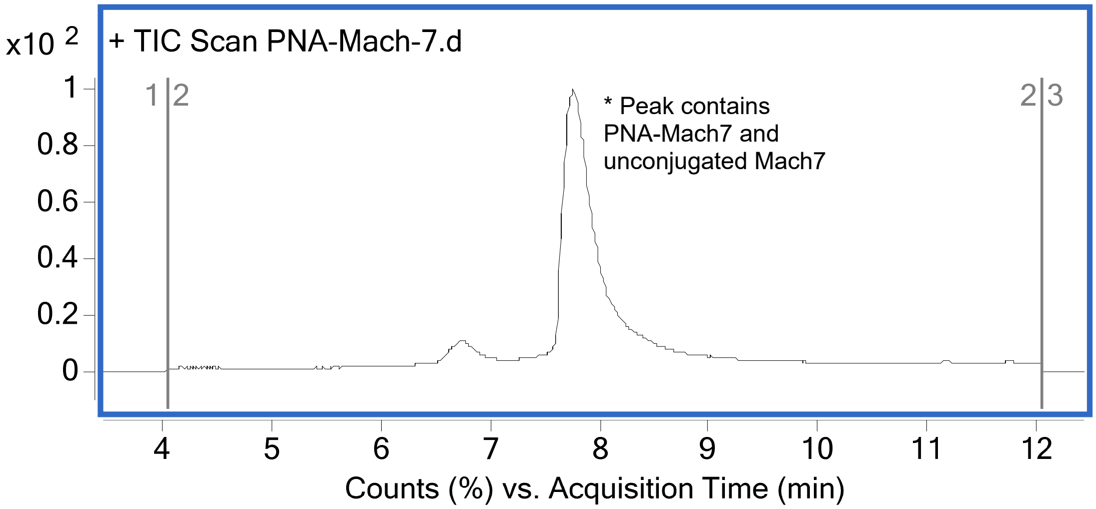

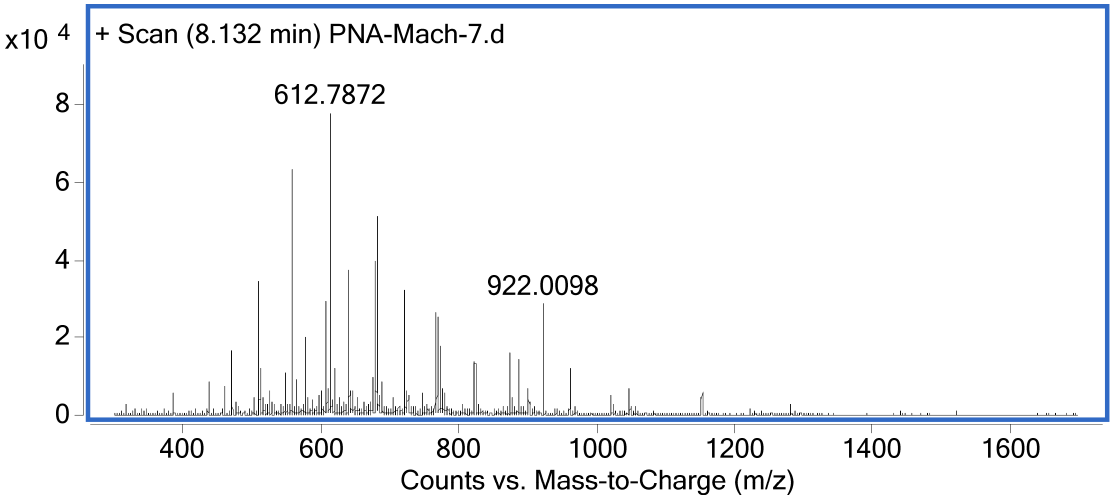

G5-DTA(C186S) (Method A)
Mass Expected: 21376.8 Da
Mass Observed: 21377.2 Da

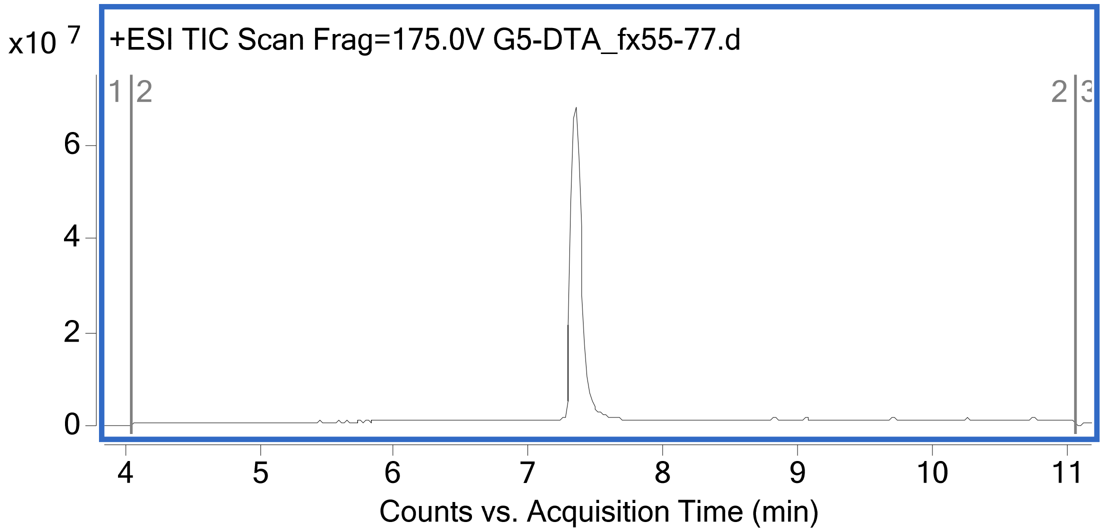

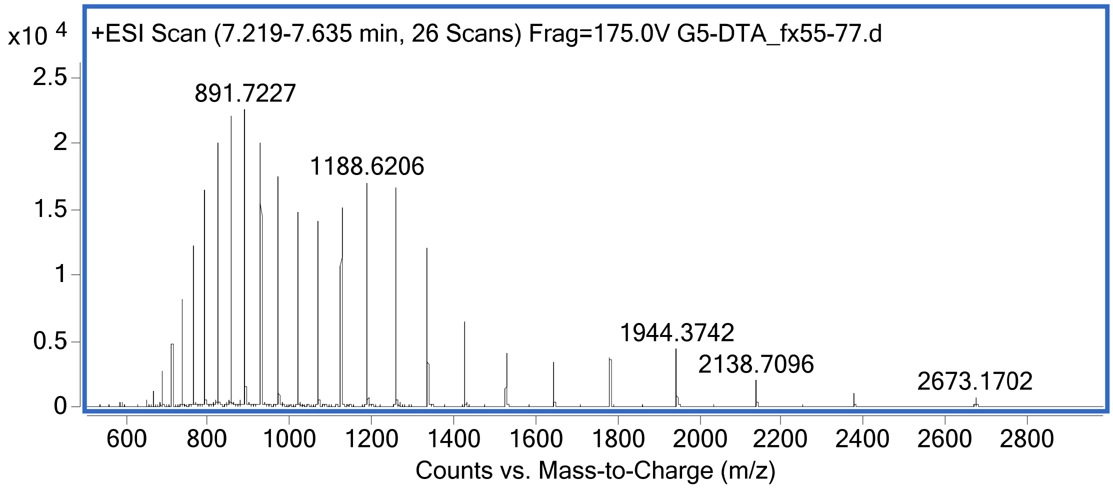

G5-DTA(C186S, E148S) (Method A)
Mass Expected: 21334.6 Da
Mass Observed: 21335.3 Da

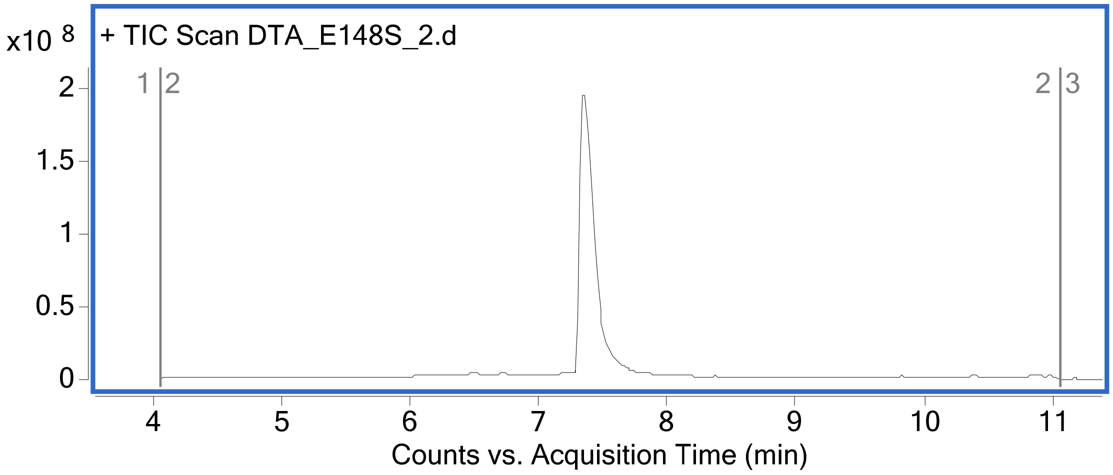

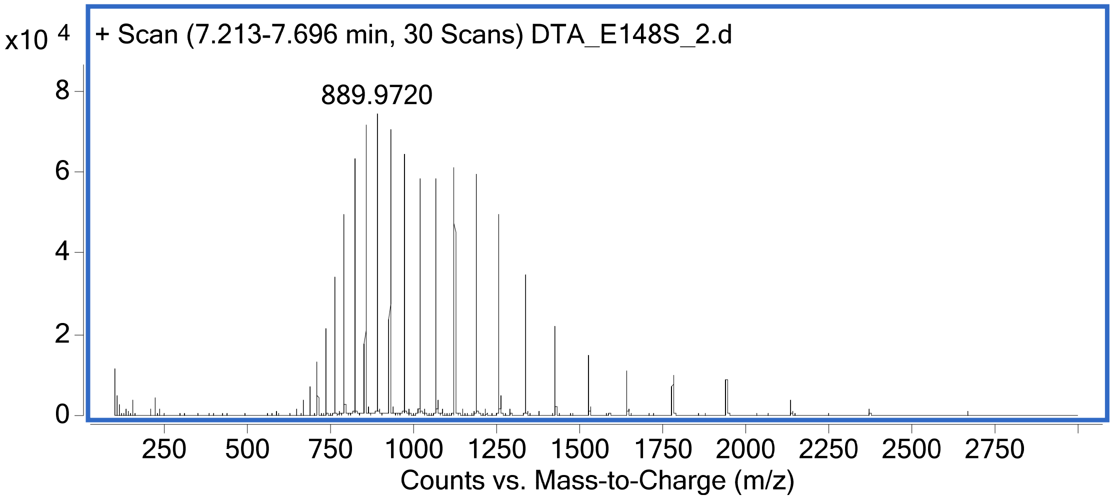

Mach3-DTA(C186S) (Method A)
Mass Expected: 26428.7 Da
Mass Observed: 26432.0 Da

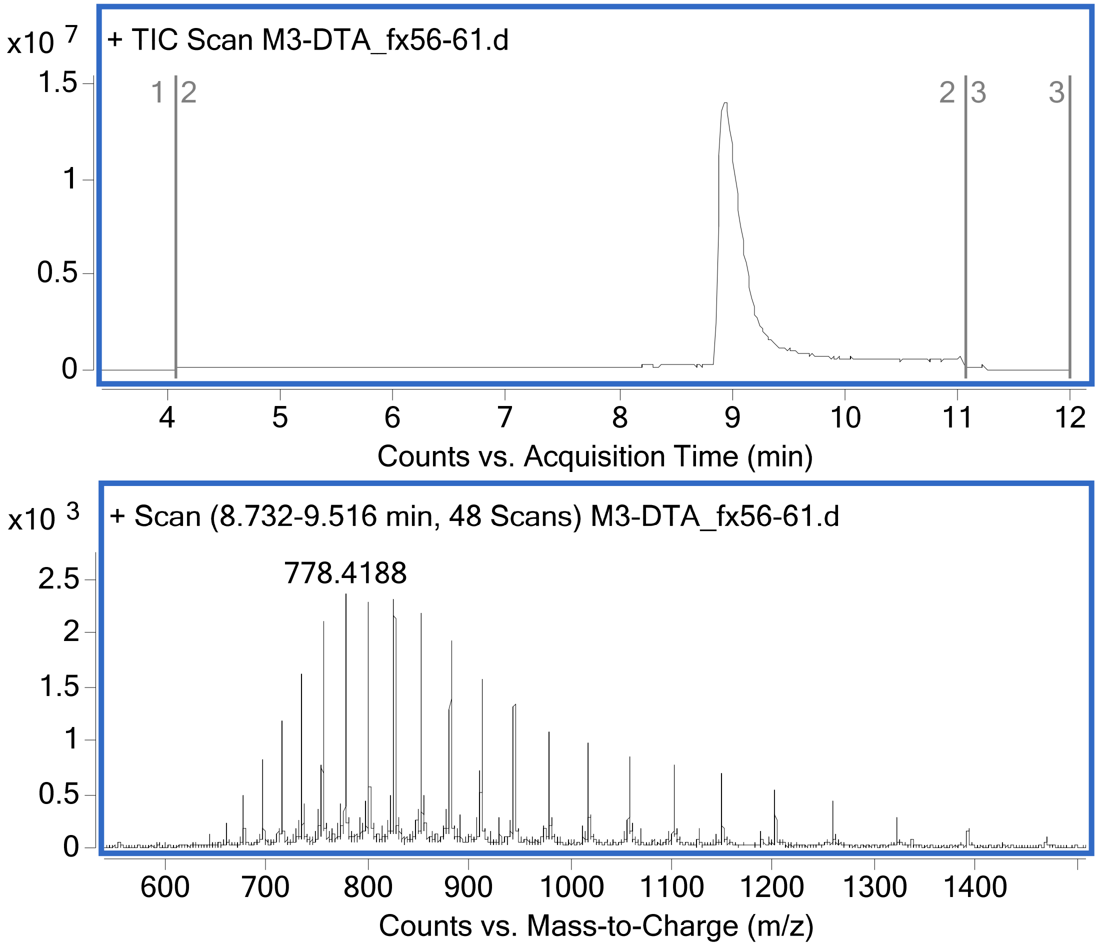

Mach3-DTA(C186S, E148S) (Method B)
Mass Expected: 26386.7 Da
Mass Observed: 26388.2 Da

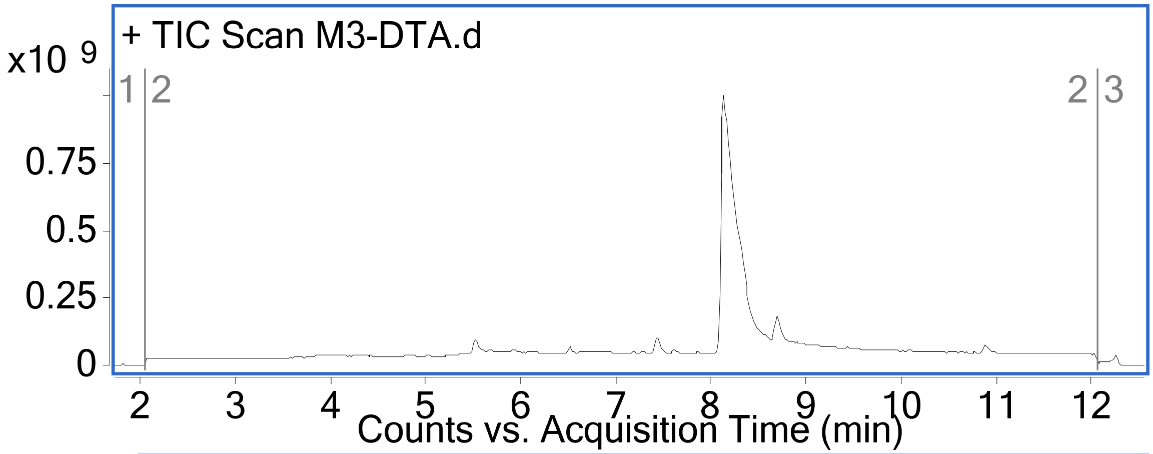

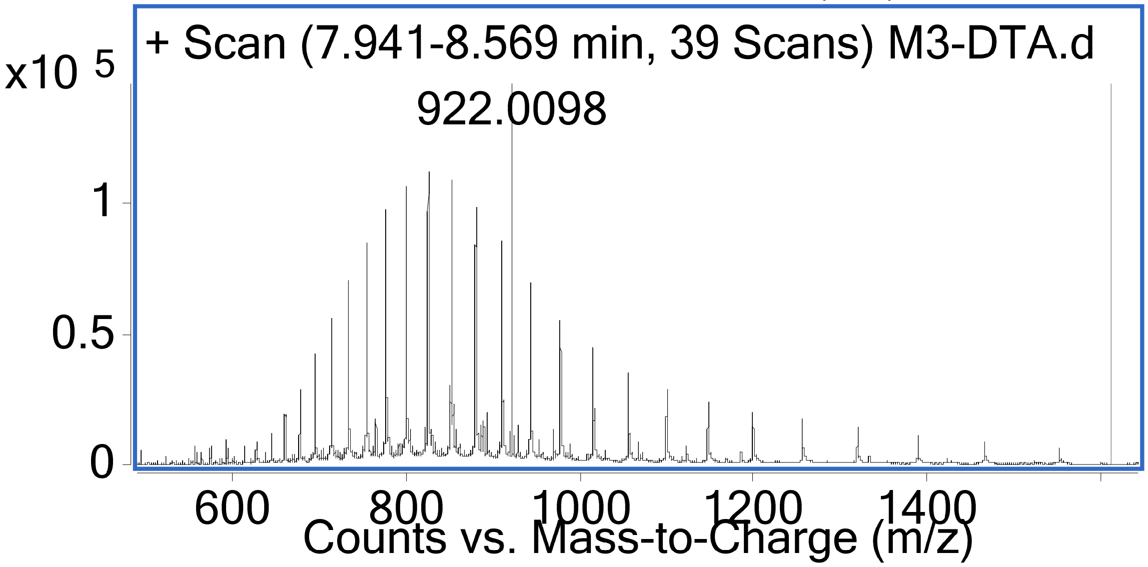

Mach7-DTA(C186S) (Method A)
Mass Expected: 27750.5 Da
Mass Observed: 27755.1 Da

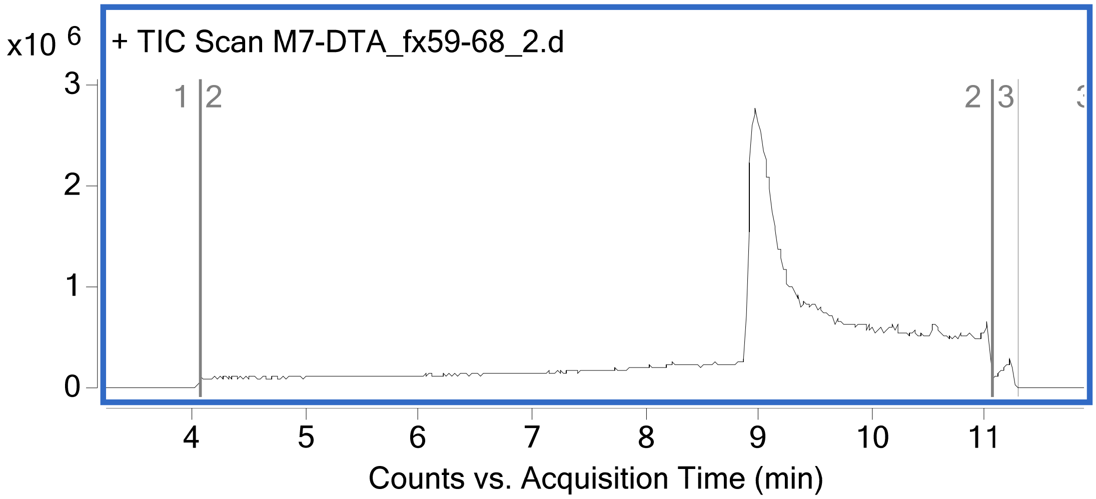

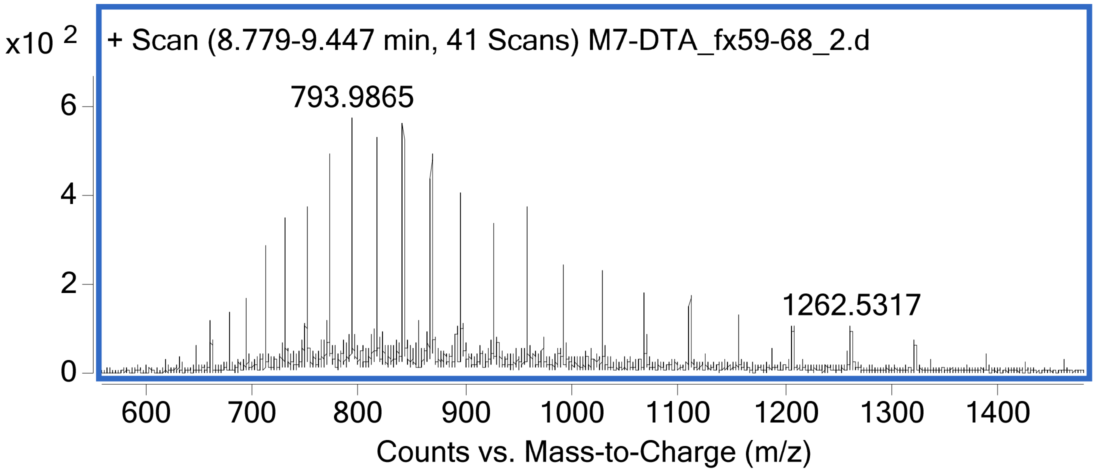

Mach7-DTA(C186S, E148S) (Method B)
Mass Expected: 27708.5 Da
Mass Observed: 27710.1 Da

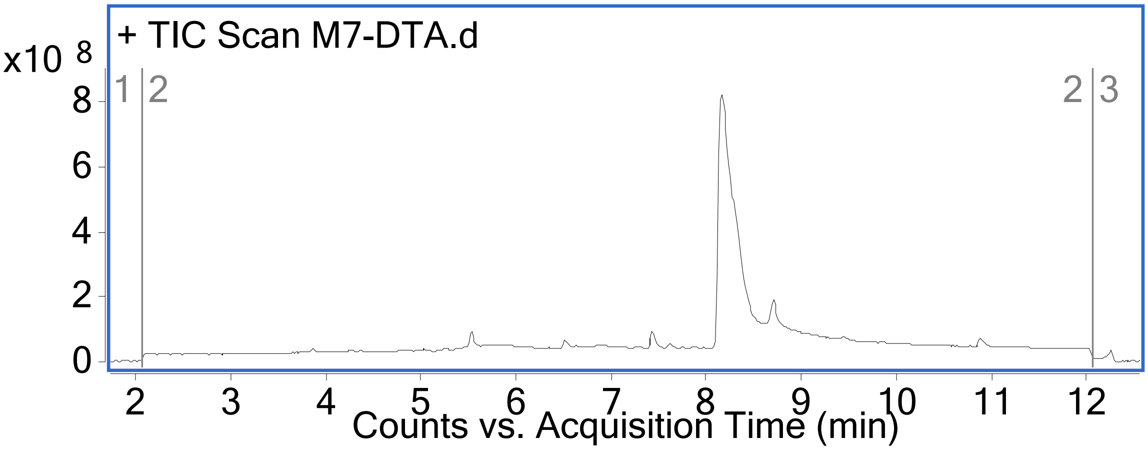

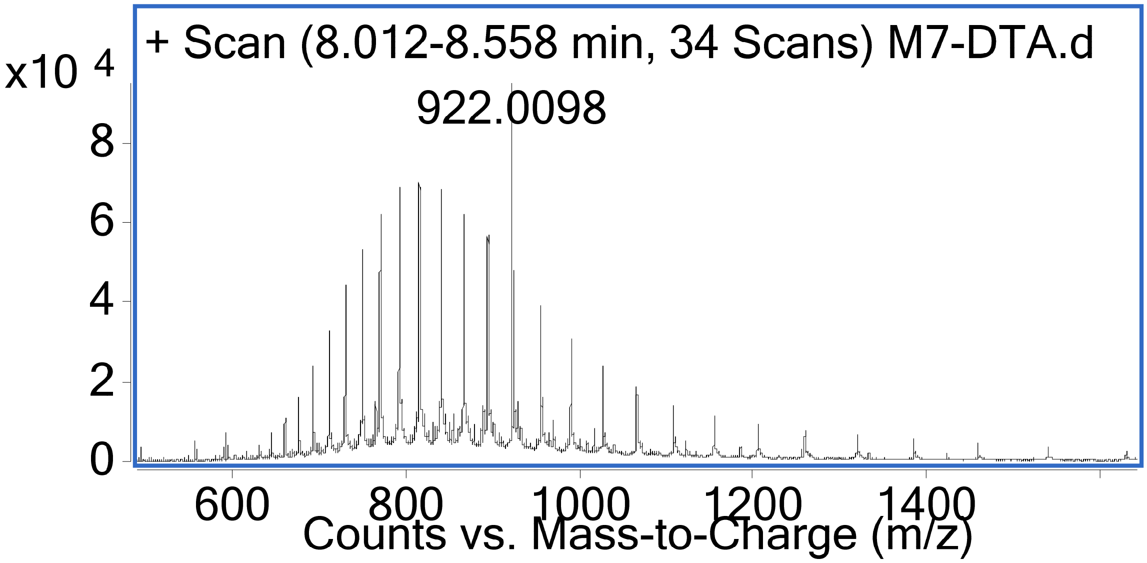

G5-EGFP (Method A)
Mass Expected: 28754.4 Da
Mass Observed: 28754.8 Da

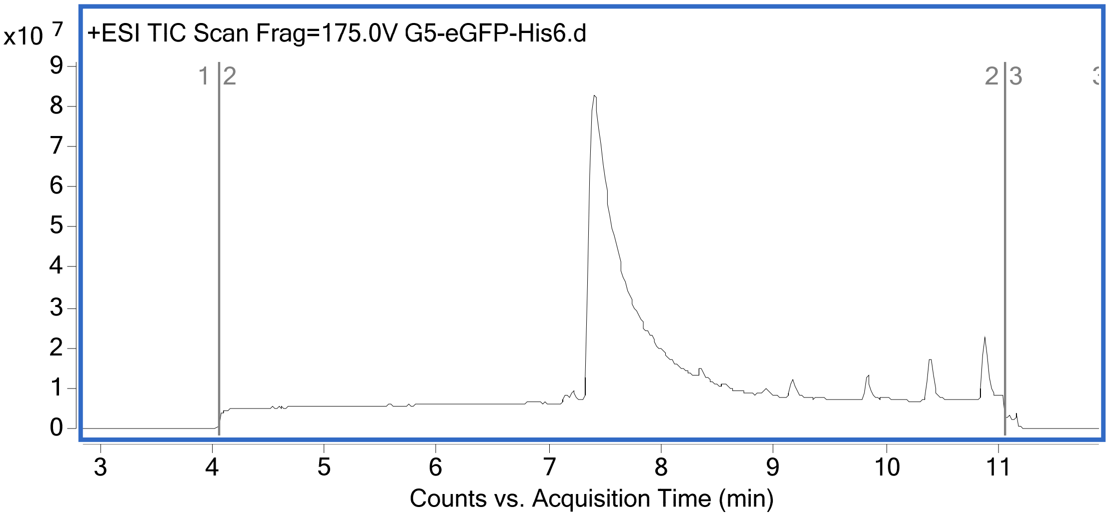

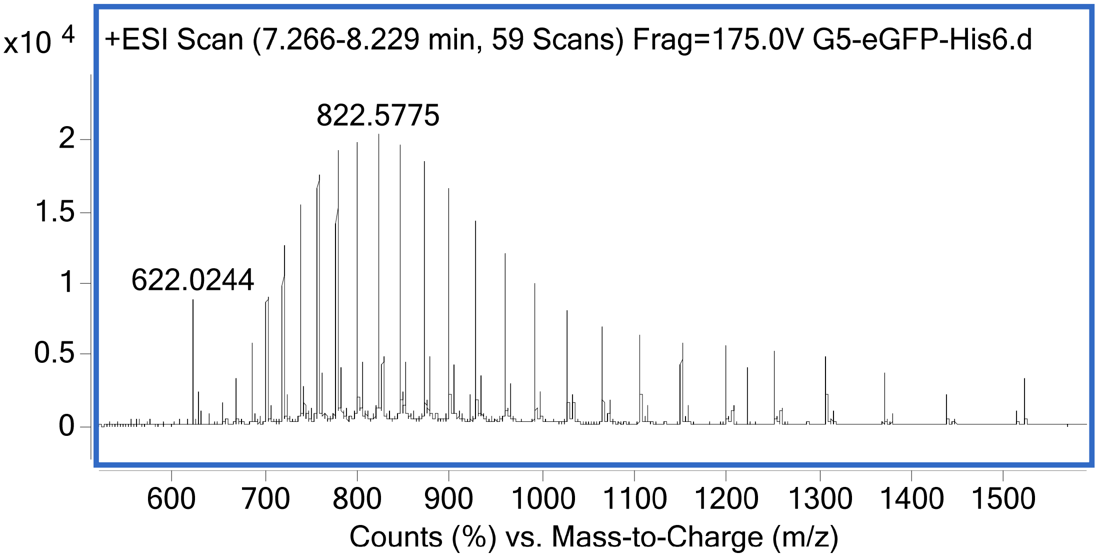

<u>Mach3-EGFP (Method B)</u>
Mass Expected: 33806.5 Da
Mass Observed: 33807.3 Da

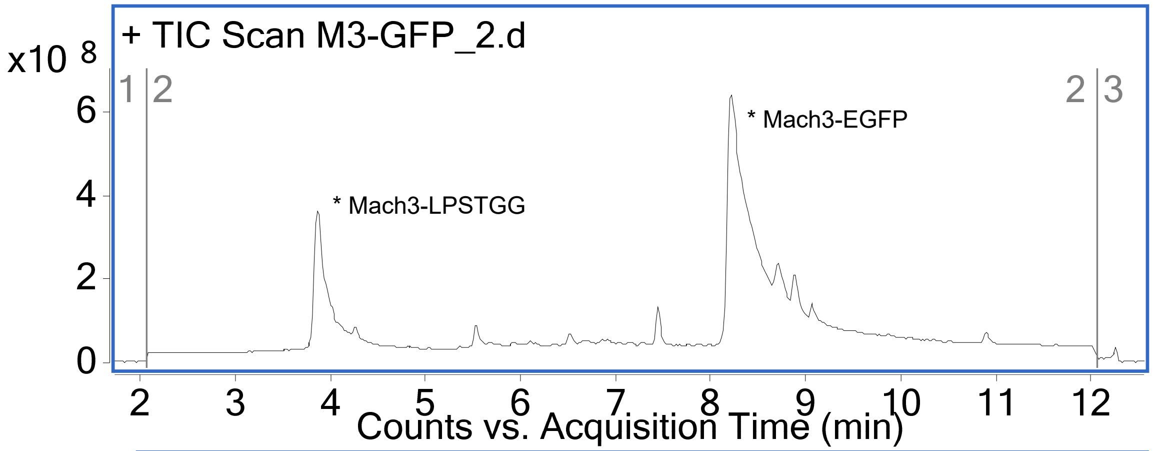

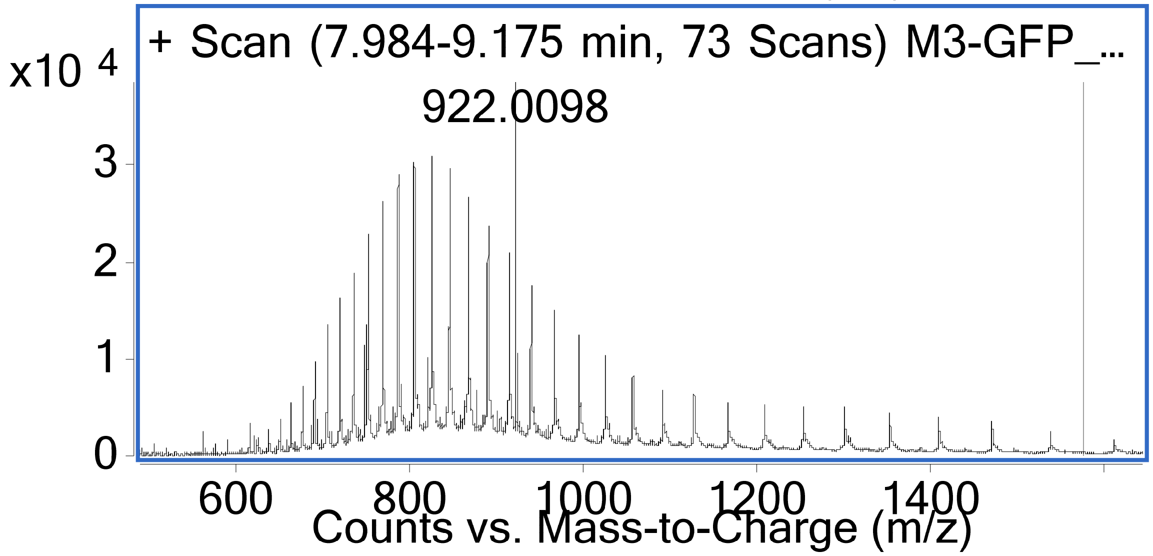

Mach7-EGFP (Method B)
Mass Expected: 35128.3 Da
Mass Observed: 35130.3 Da

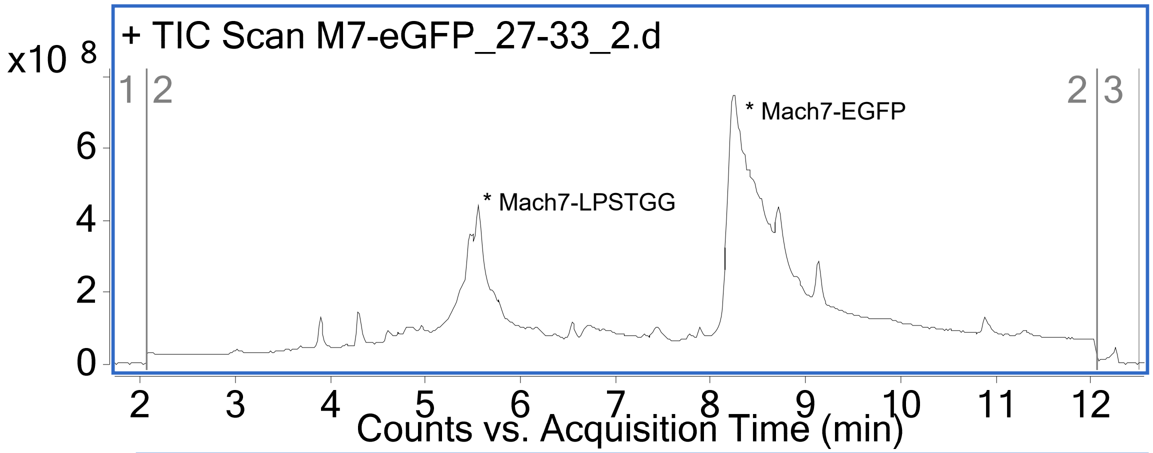

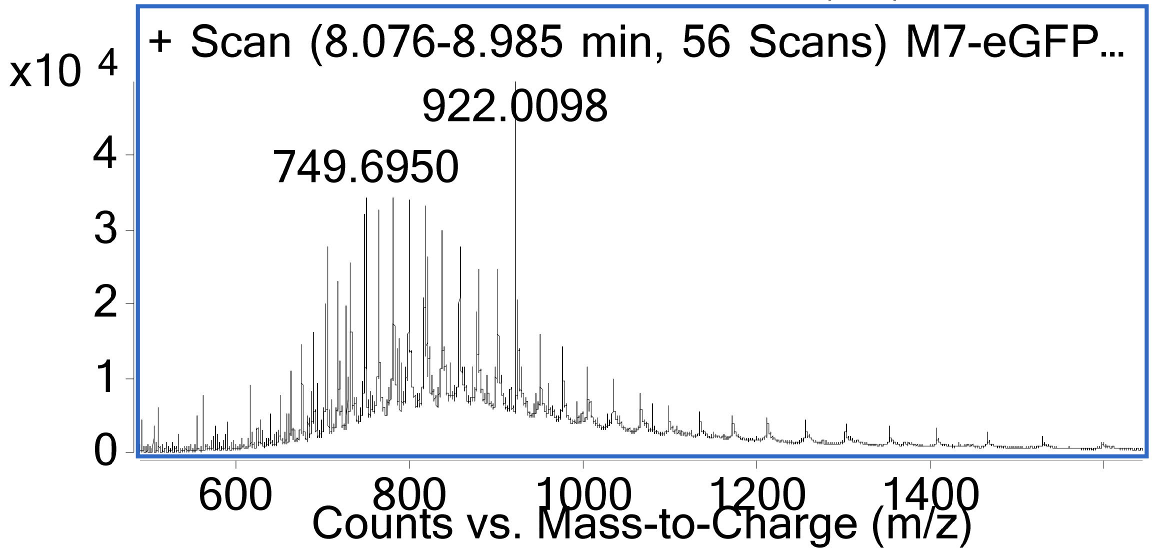

### Appendix III: Topological fingerprints

CB_Index = Condensed Bit-vector index, also used in the main text figures
TF_Index = Topological Fingerprint index, for the corresponding CB_index

All ON bits out of the 2048-bits have been represented in the following set of figures. The radius of exploration goes from 0 (atom, itself) to 3 nearest neighbors. The coloring scheme denotes the node atom in blue, atoms which are a part of an aromatic ring in yellow, connected neighbors as a part of the topological exploration in black, and the unexplored neighboring atoms and nodes in gray.

| CB_Index | TF_Index | CB_Index | TF_Index | CB_Index | TF_Index | CB_Index | TF_Index |
|---|---|---|---|---|---|---|---|
| 1 | 1 | 51 | 585 | 101 | 1114 | 151 | 1693 |
| 2 | 11 | 52 | 623 | 102 | 1117 | 152 | 1719 |
| 3 | 22 | 53 | 625 | 103 | 1127 | 153 | 1731 |
| 4 | 27 | 54 | 650 | 104 | 1139 | 154 | 1736 |
| 5 | 32 | 55 | 667 | 105 | 1141 | 155 | 1737 |
| 6 | 67 | 56 | 671 | 106 | 1143 | 156 | 1750 |
| 7 | 70 | 57 | 680 | 107 | 1145 | 157 | 1751 |
| 8 | 74 | 58 | 708 | 108 | 1152 | 158 | 1752 |
| 9 | 79 | 59 | 713 | 109 | 1158 | 159 | 1754 |
| 10 | 80 | 60 | 724 | 110 | 1171 | 160 | 1758 |
| 11 | 119 | 61 | 727 | 111 | 1185 | 161 | 1773 |
| 12 | 132 | 62 | 739 | 112 | 1199 | 162 | 1778 |
| 13 | 140 | 63 | 742 | 113 | 1213 | 163 | 1783 |
| 14 | 150 | 64 | 745 | 114 | 1221 | 164 | 1785 |
| 15 | 173 | 65 | 759 | 115 | 1226 | 165 | 1791 |
| 16 | 197 | 66 | 776 | 116 | 1258 | 166 | 1794 |
| 17 | 204 | 67 | 784 | 117 | 1259 | 167 | 1805 |
| 18 | 220 | 68 | 785 | 118 | 1267 | 168 | 1840 |
| 19 | 222 | 69 | 786 | 119 | 1268 | 169 | 1844 |
| 20 | 227 | 70 | 806 | 120 | 1283 | 170 | 1847 |
| 21 | 229 | 71 | 807 | 121 | 1287 | 171 | 1849 |
| 22 | 231 | 72 | 831 | 122 | 1290 | 172 | 1873 |
| 23 | 272 | 73 | 857 | 123 | 1301 | 173 | 1876 |
| 24 | 280 | 74 | 878 | 124 | 1307 | 174 | 1879 |
| 25 | 283 | 75 | 889 | 125 | 1313 | 175 | 1882 |
| 26 | 289 | 76 | 894 | 126 | 1325 | 176 | 1898 |
| 27 | 293 | 77 | 900 | 127 | 1349 | 177 | 1910 |
| 28 | 294 | 78 | 926 | 128 | 1357 | 178 | 1911 |
| 29 | 295 | 79 | 931 | 129 | 1380 | 179 | 1912 |
| 30 | 305 | 80 | 955 | 130 | 1388 | 180 | 1917 |
| 31 | 310 | 81 | 966 | 131 | 1427 | 181 | 1926 |
| 32 | 321 | 82 | 971 | 132 | 1431 | 182 | 1928 |
| 33 | 328 | 83 | 981 | 133 | 1451 | 183 | 1937 |
| 34 | 329 | 84 | 983 | 134 | 1452 | 184 | 1946 |
| 35 | 362 | 85 | 989 | 135 | 1459 | 185 | 1947 |
| 36 | 364 | 86 | 1014 | 136 | 1462 | 186 | 1969 |
| 37 | 368 | 87 | 1017 | 137 | 1507 | 187 | 1970 |
| 38 | 376 | 88 | 1019 | 138 | 1517 | 188 | 2006 |
| 39 | 378 | 89 | 1022 | 139 | 1544 | 189 | 2013 |
| 40 | 389 | 90 | 1027 | 140 | 1547 | 190 | 2022 |
| 41 | 394 | 91 | 1028 | 141 | 1558 | 191 | 2042 |
| 42 | 412 | 92 | 1031 | 142 | 1564 | | |
| 43 | 420 | 93 | 1034 | 143 | 1573 | | |
| 44 | 425 | 94 | 1057 | 144 | 1601 | | |
| 45 | 473 | 95 | 1066 | 145 | 1602 | | |
| 46 | 482 | 96 | 1072 | 146 | 1607 | | |
| 47 | 545 | 97 | 1082 | 147 | 1633 | | |
| 48 | 553 | 98 | 1088 | 148 | 1656 | | |
| 49 | 561 | 99 | 1104 | 149 | 1661 | | |
| 50 | 575 | 100 | 1110 | 150 | 1685 | | |

Linker 2

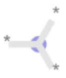

1

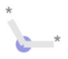

80

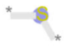

116

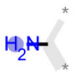

140

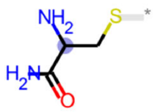

210

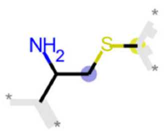

222

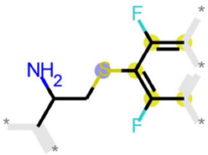

302

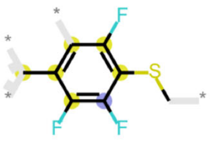

319

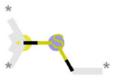

341

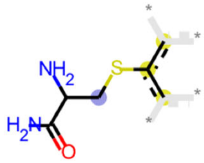

480

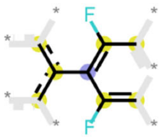

551

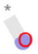

650

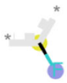

699

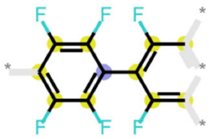

773

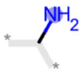

786

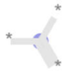

807

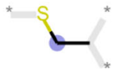

1093

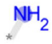

1171

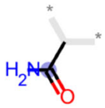

1212

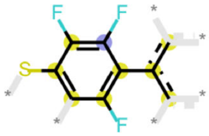

1234

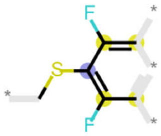

1305

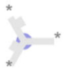

1380

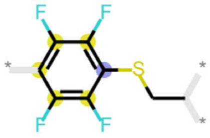

1436

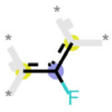

1445

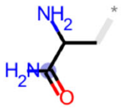

1654

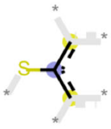

1716

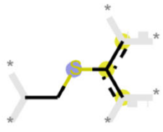

1733

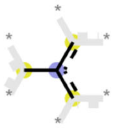

1747

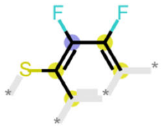

1821

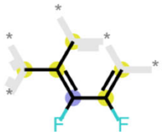

1848

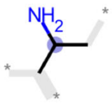

1849

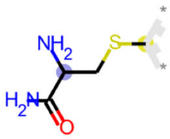

1855

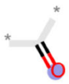

1917

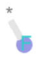

1928

Linker 3

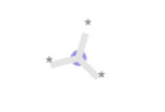
1

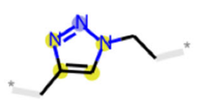
72

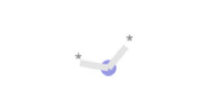
80

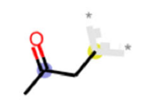
109

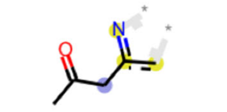
124

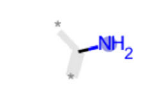
140

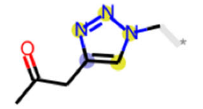
186

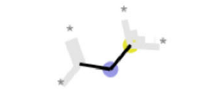
325

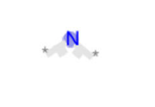
378

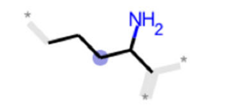
412

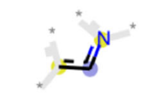
464

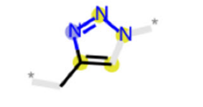
488

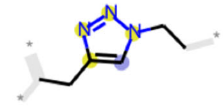
567

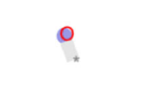
650

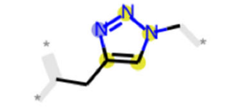
669

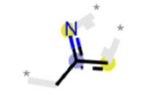
724

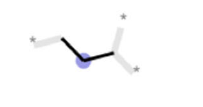
739

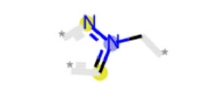
747

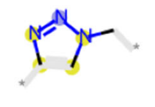
781

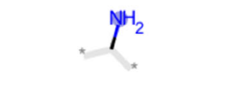
786

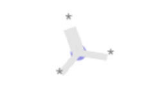
807

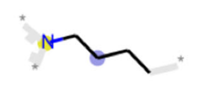
893

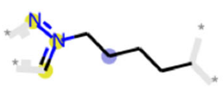
913

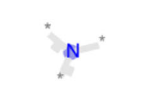
935

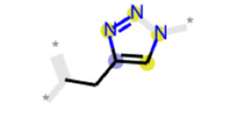
980

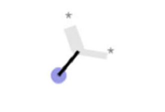
1017

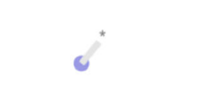
1057

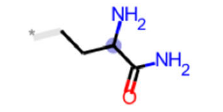
1127

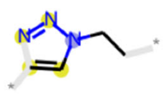
1141

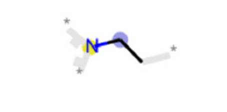
1145

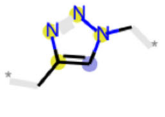
1155

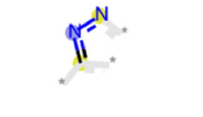
1164

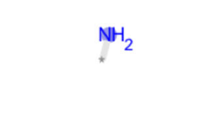
1171

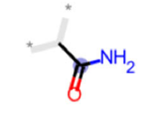
1212

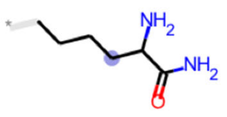
1214

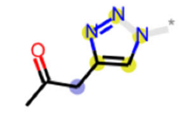

1259

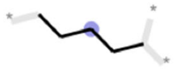

1283

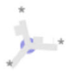

1380

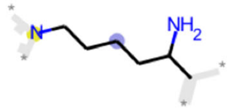

1536

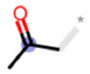

1550

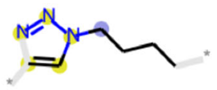

1582

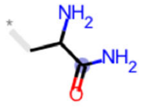

1654

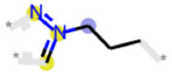

1689

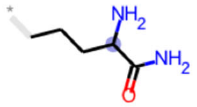

1728

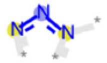

1783

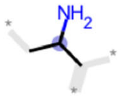

1849

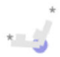

1873

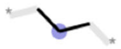

1911

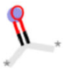

1917

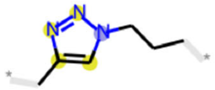

1971

Alanine

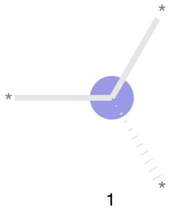
1

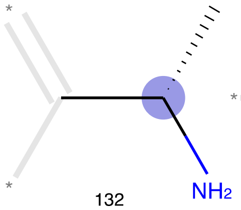
132

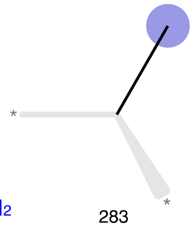
283

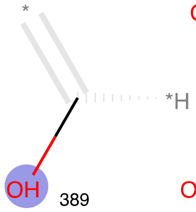
389

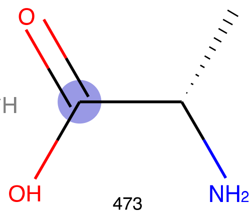
473

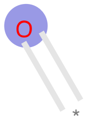
650

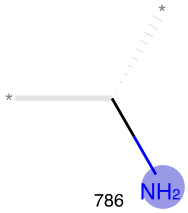
786

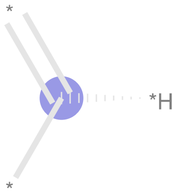
807

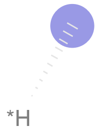
1057

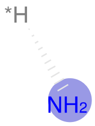
1171

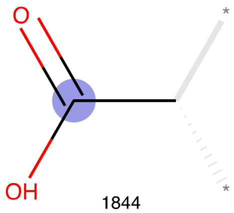
1844

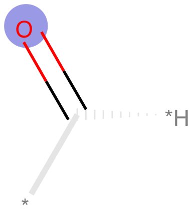
1917

Beta-Alanine

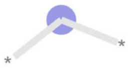

80

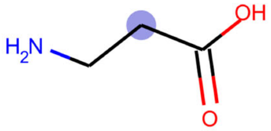

173

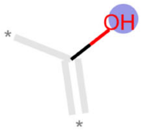

389

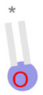

650

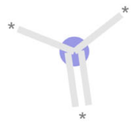

807

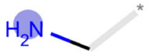

981

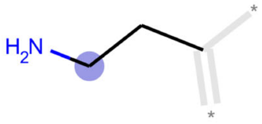

1088

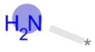

1171

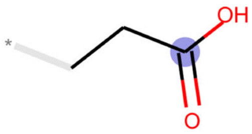

1287

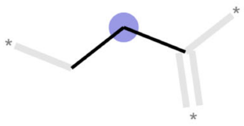

1564

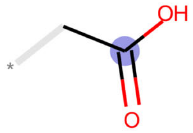

1737

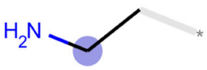

1840

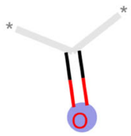

1917

Aminohexanoic acid

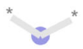

80

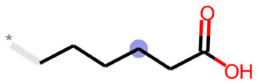

295

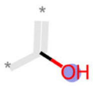

389

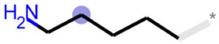

561

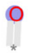

650

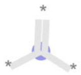

807

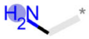

981

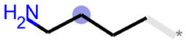

1082

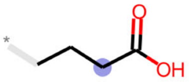

1110

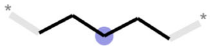

1143

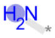

1171

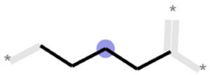

1267

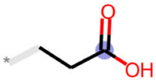

1287

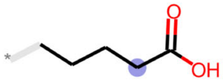

1301

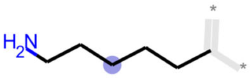

1462

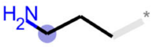

1517

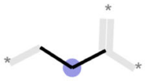

1564

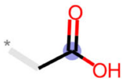

1737

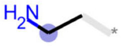

1840

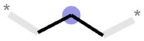

1911

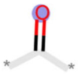

1917

Arginine

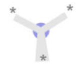
1

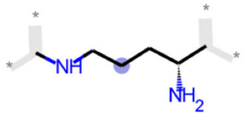
67

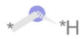
80

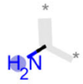
140

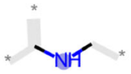
197

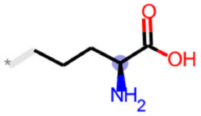
289

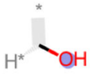
389

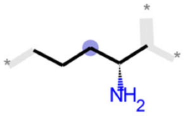
412

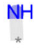
623

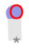
650

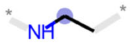
667

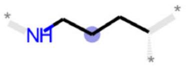
708

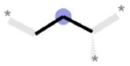
739

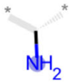
786

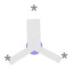
807

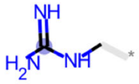
894

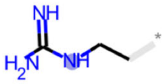
983

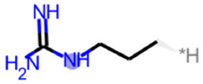
1027

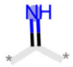
1104

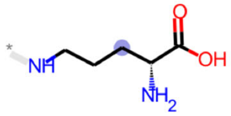
1127

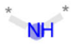
1152

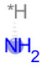
1171

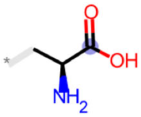
1427

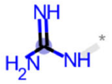
1451

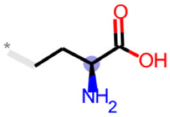
1791

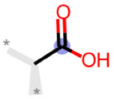
1844

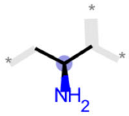
1849

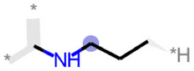
1876

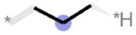
1911

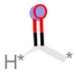
1917

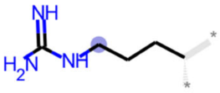
2042

Asparagine

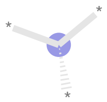

1

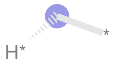

80

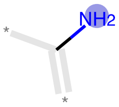

140

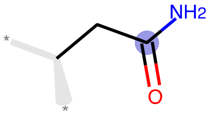

376

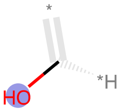

389

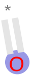

650

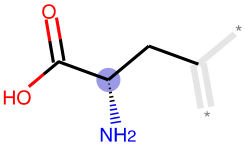

776

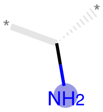

786

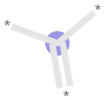

807

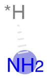

1171

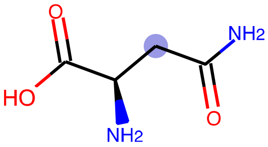

1213

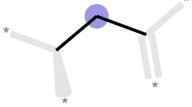

1290

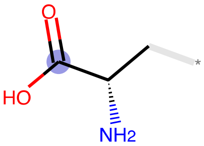

1427

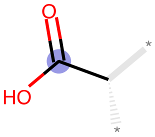

1844

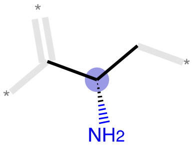

1849

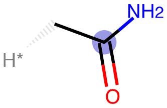

1898

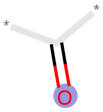

1917

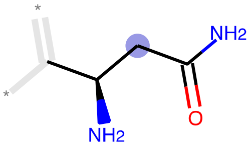

1946

Aspartic acid

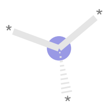

1

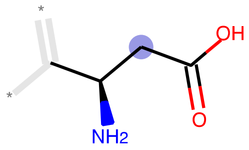

70

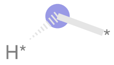

80

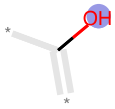

389

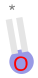

650

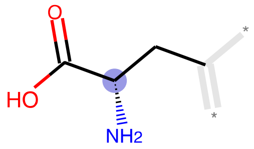

776

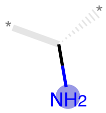

786

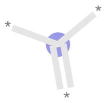

807

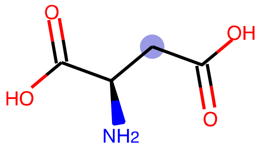

989

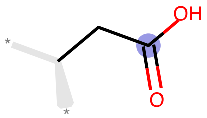

1141

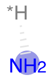

1171

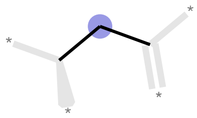

1290

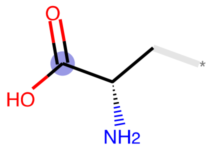

1427

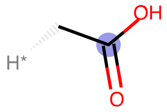

1737

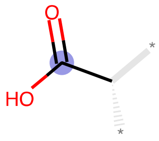

1844

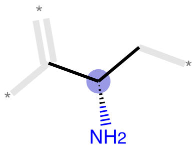

1849

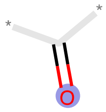

1917

Cysteine

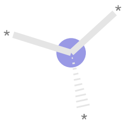

1

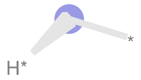

80

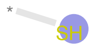

229

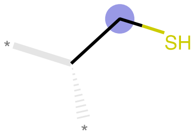

321

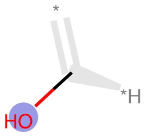

389

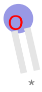

650

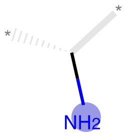

786

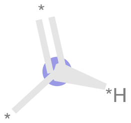

807

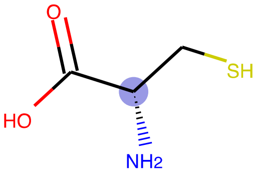

1072

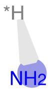

1171

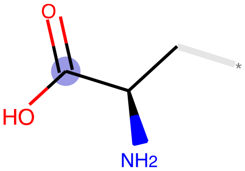

1427

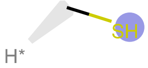

1459

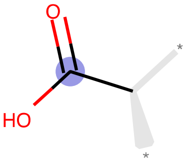

1844

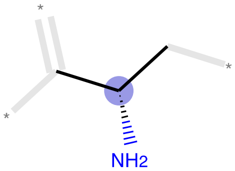

1849

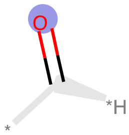

1917

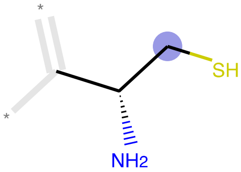

1926

### Glutamic acid

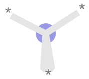

1

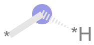

80

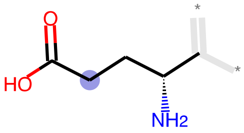

293

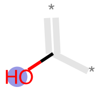

389

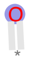

650

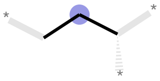

739

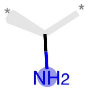

786

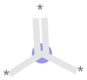

807

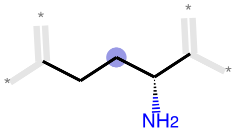

900

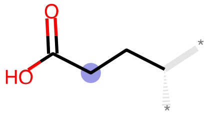

955

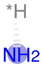

1171

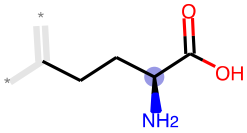

1258

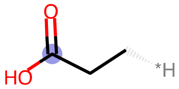

1287

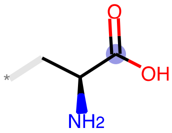

1427

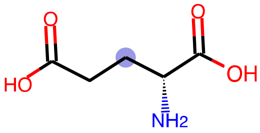

1547

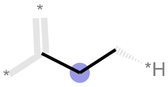

1564

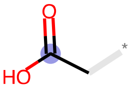

1737

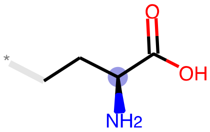

1791

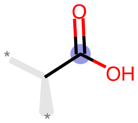

1844

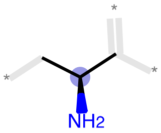

1849

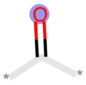

1917

Glutamine

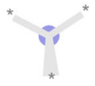

1

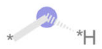

80

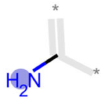

140

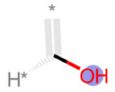

389

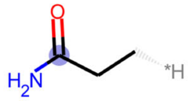

545

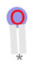

650

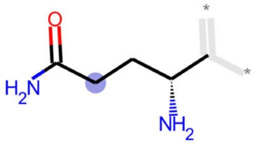

671

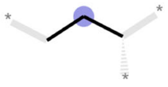

739

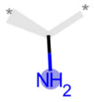

786

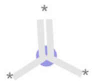

807

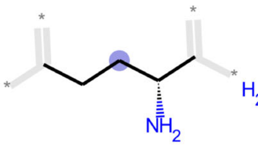

900

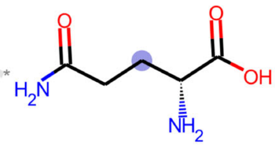

971

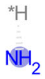

1171

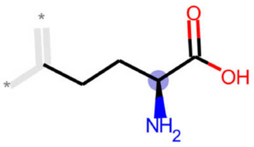

1258

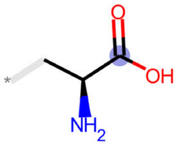

1427

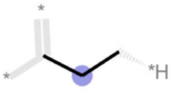

1564

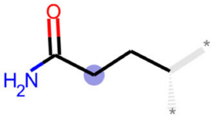

1752

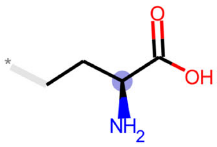

1791

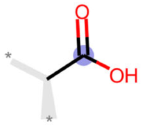

1844

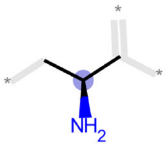

1849

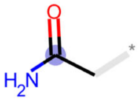

1898

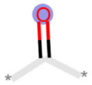

1917

Glycine

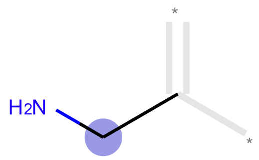

27

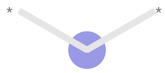
80

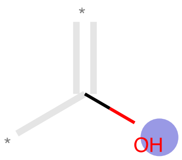

389

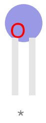

650

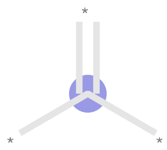
807

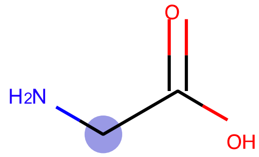

966

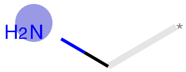

981

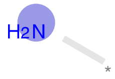

1171

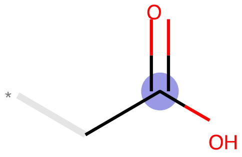

1737

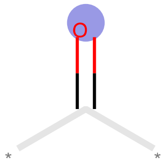

1917

Histidine

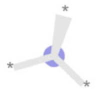
1

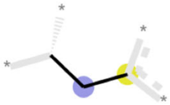
79

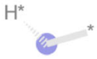
80

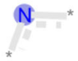
378

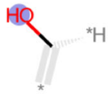
389

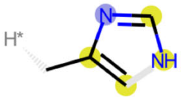
575

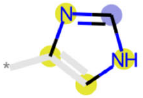
625

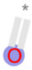
650

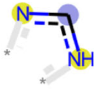
713

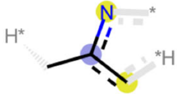
724

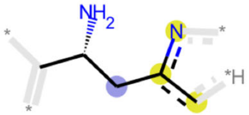
785

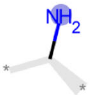
786

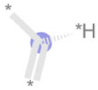
807

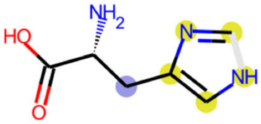
889

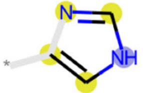
931

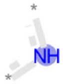
1114

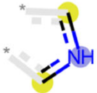
1145

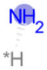
1171

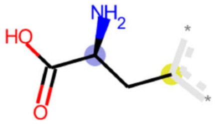
1221

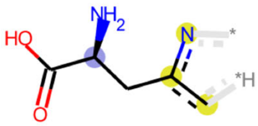
1259

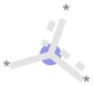
1380

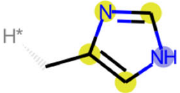
1388

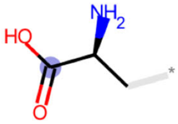
1427

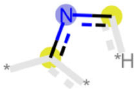
1452

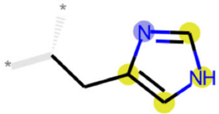
1601

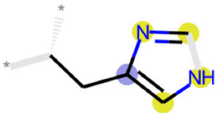
1633

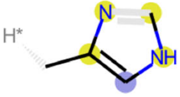
1758

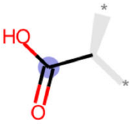
1844

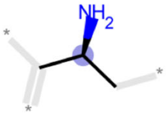
1849

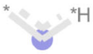
1873

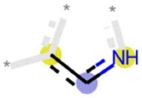
1879

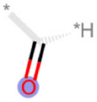
1917

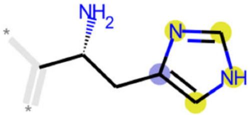
1969

Isoleucine

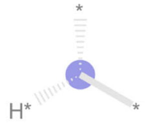

1

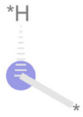

80

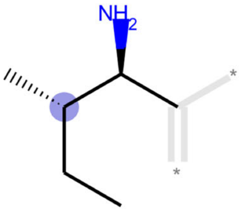

280

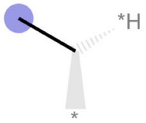

283

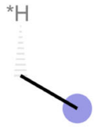

294

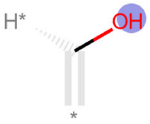

389

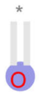

650

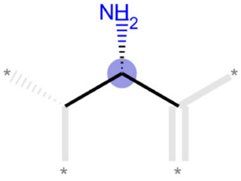

727

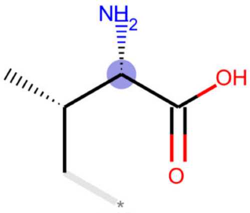

784

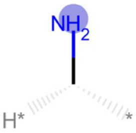

786

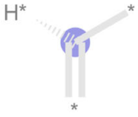

807

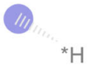

1057

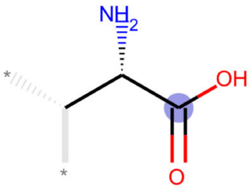

1139

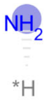

1171

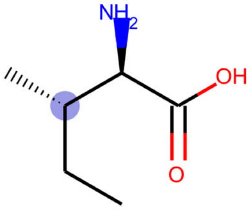

1185

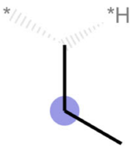

1544

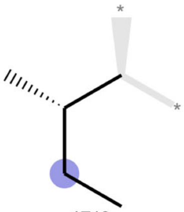

1719

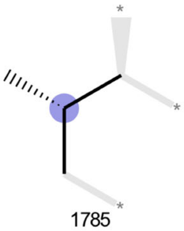

1785

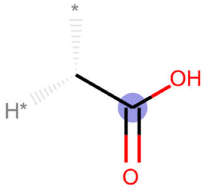

1844

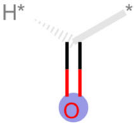

1917

Leucine

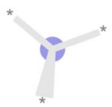
1

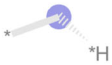
80

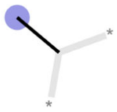
283

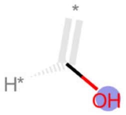
389

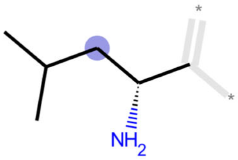
425

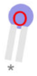
650

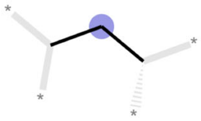
680

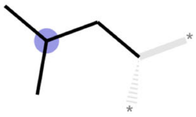
759

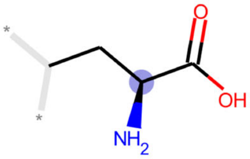
776

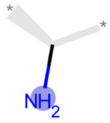
786

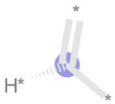
807

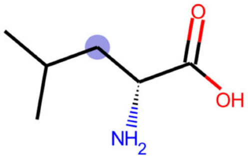
878

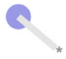
1057

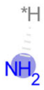
1171

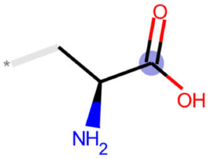
1427

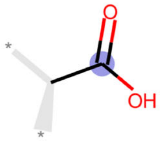
1844

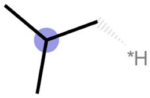
1847

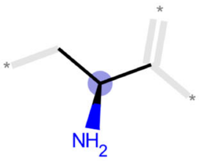
1849

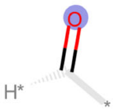
1917

Lysine

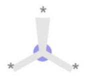
1

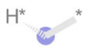
80

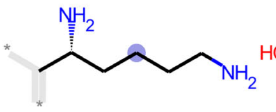
220

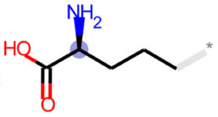
289

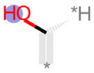
389

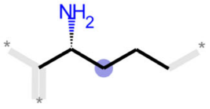
412

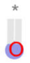
650

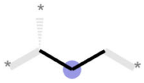
739

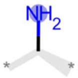
786

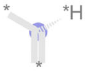
807

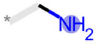
981

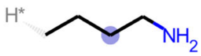
1082

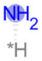
1171

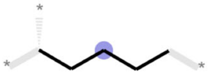
1283

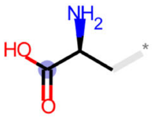
1427

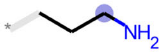
1517

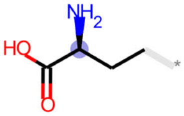
1791

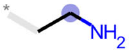
1840

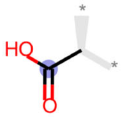
1844

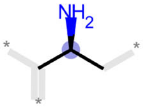
1849

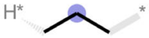
1911

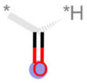
1917

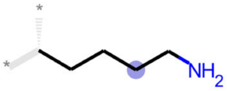
1928

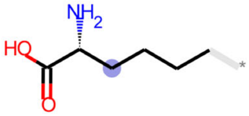
2006

Methionine

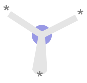
1

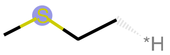
11

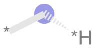
80

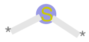
116

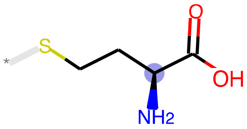
368

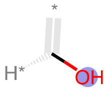
389

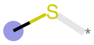
394

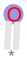
650

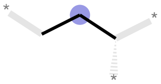
739

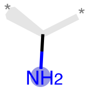
786

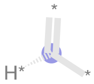
807

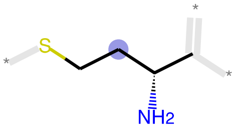
1021

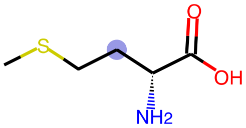
1034

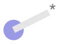
1057

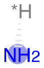
1171

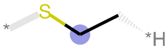
1182

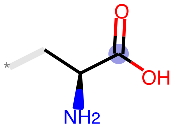
1427

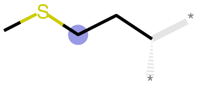
1558

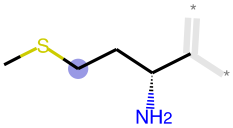
1661

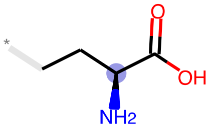
1791

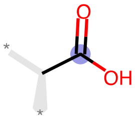
1844

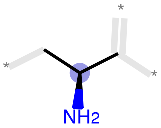
1849

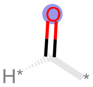
1917

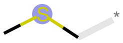
1942

Phenylalanine

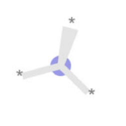

1

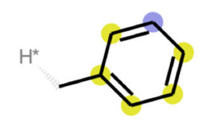

32

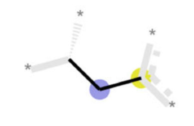

79

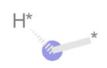

80

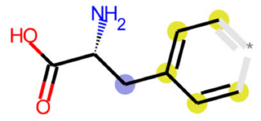

150

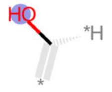

389

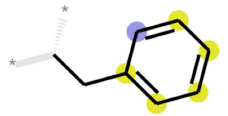

420

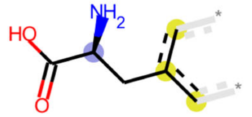

585

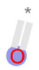

650

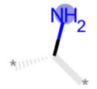

786

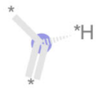

807

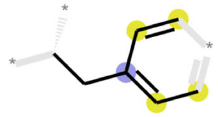

857

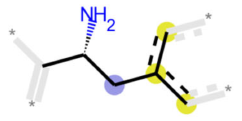

1017

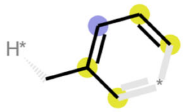

1066

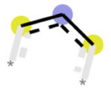

1088

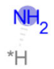

1171

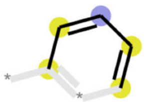

1199

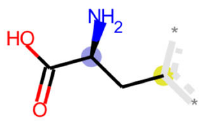

1221

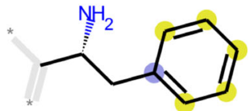

1307

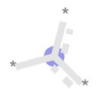

1380

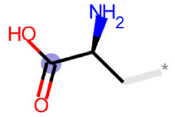

1427

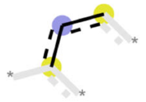

1750

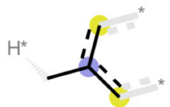

1754

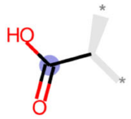

1844

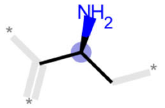

1849

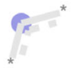

1873

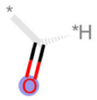

1917

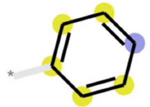

1947

Proline

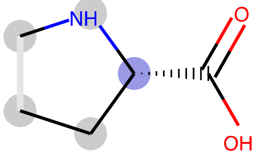
74

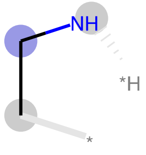
305

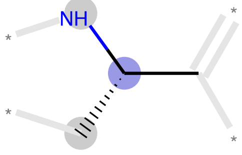
362

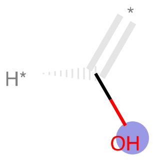
389

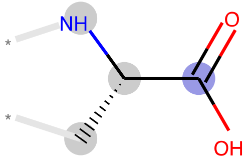
553

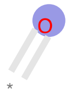
650

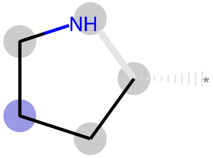
742

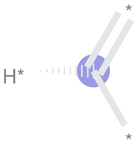
807

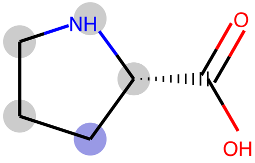
831

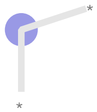
926

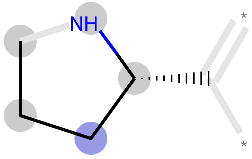
1014

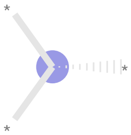
1019

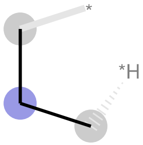
1028

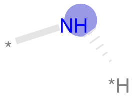
1114

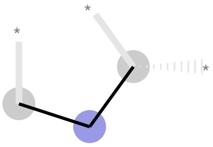
1325

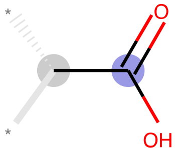
1431

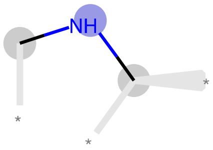
1507

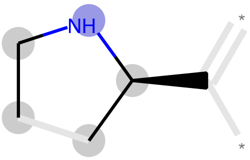
1736

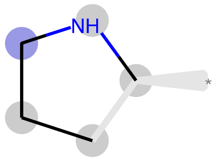
1912

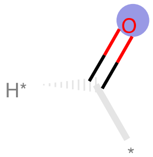
1917

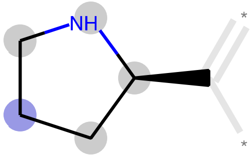
2022

Serine

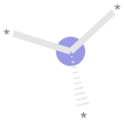

1

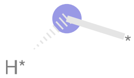

80

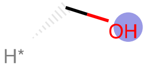

222

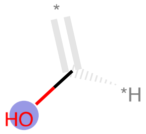

389

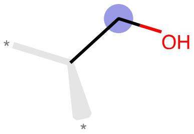

482

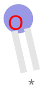

650

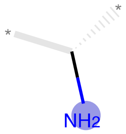

786

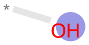

807

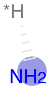

1171

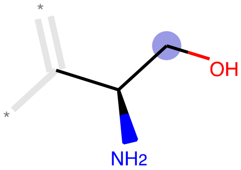

1268

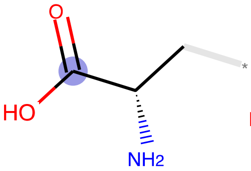

1427

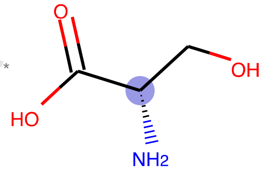

1751

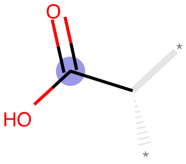

1844

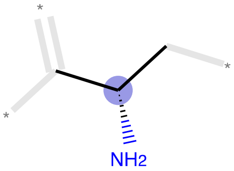

1849

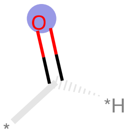

1917

Threonine

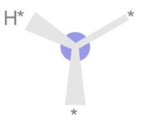

1

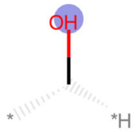

227

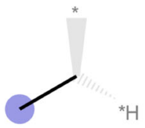

283

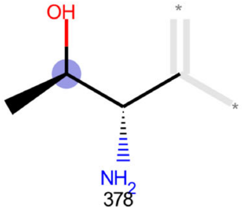

378

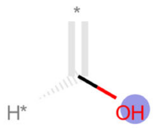

389

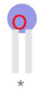

650

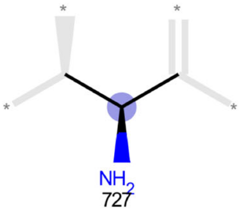

727

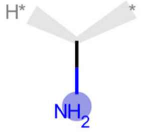

786

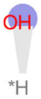

807

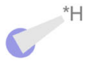

1057

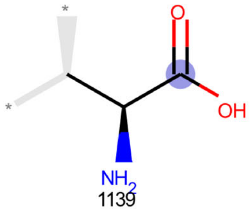

1139

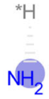

1171

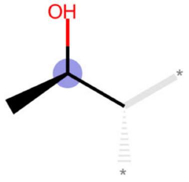

1693

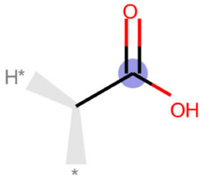

1844

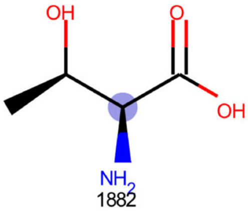

1882

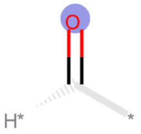

1917

Tryptophan

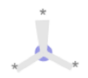
1

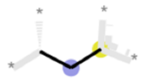
79

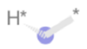
80

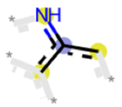
119

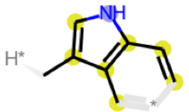
140

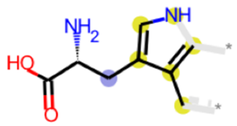
204

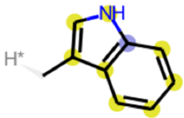
272

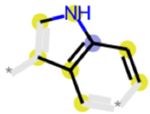
328

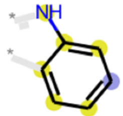
329

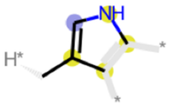
364

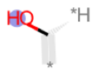
389

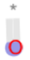
650

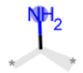
786

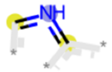
806

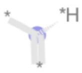
807

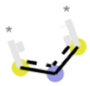
1088

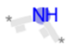
1114

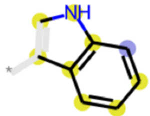
1158

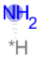
1171

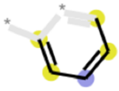
1199

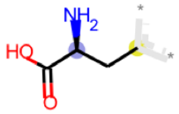
1221

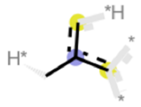
1349

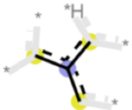
1357

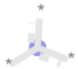
1380

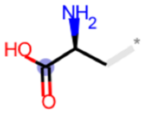
1427

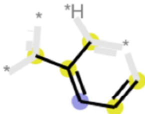
1573

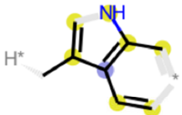
1607

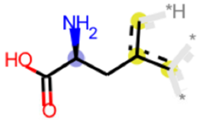
1656

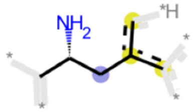
1685

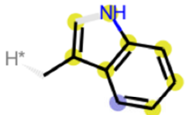
1731

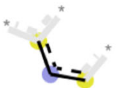
1750

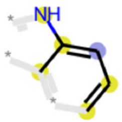
1783

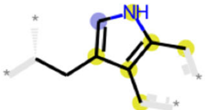
1794

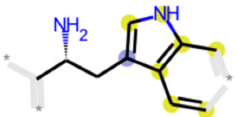
1805

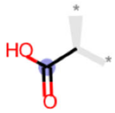
1844

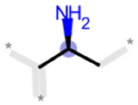

1849

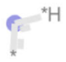

1873

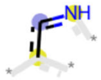

1879

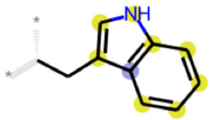

1910

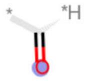

1917

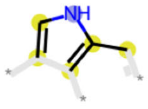

1937

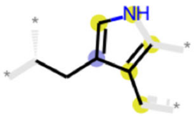

1970

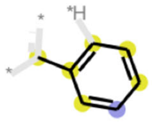

2013

Tyrosine

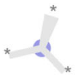

1

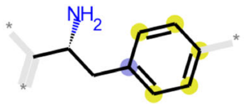

22

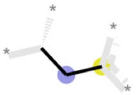

79

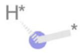

80

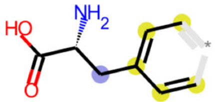

150

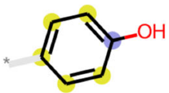

231

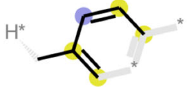

310

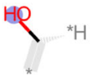

389

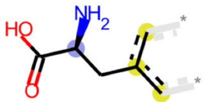

585

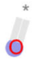

650

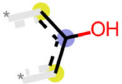

745

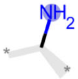

786

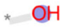

807

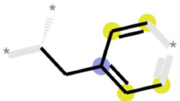

857

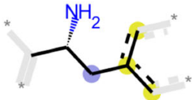

1017

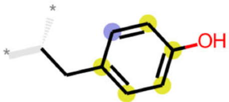

1117

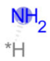

1171

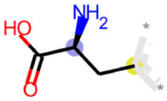

1221

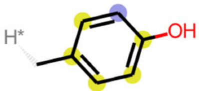

1226

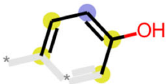

1313

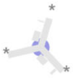

1380

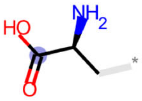

1427

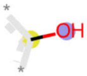

1602

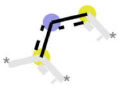

1750

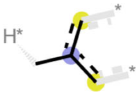

1754

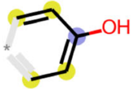

1778

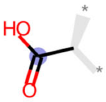

1844

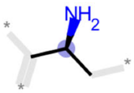

1849

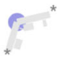

1873

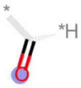

1917

Valine

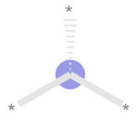
1

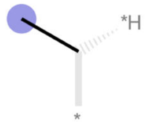
283

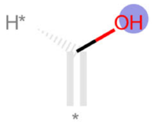
389

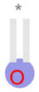
650

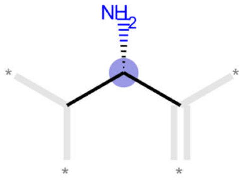
727

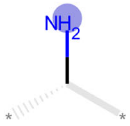
786

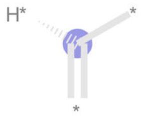
807

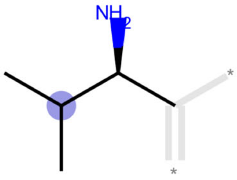
1022

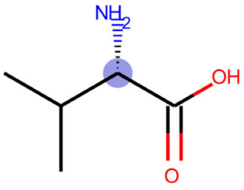
1031

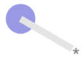
1057

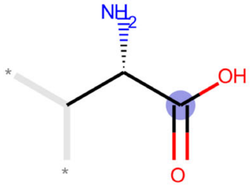
1139

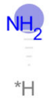
1171

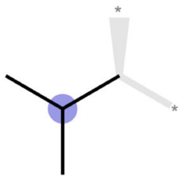
1773

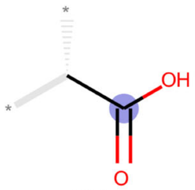
1844

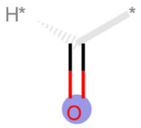
1917
